# Gene- and domain-aware calibration increases the clinical utility of variant effect predictors

**DOI:** 10.64898/2026.02.17.706269

**Authors:** Yile Chen, Shawn Fayer, Shantanu Jain, Mariam Benazouz, Yuriy Sverchkov, Jeremy Stone, Himanshu Sharma, Timothy Bergquist, Ross Stewart, Sean D. Mooney, Mark Craven, Predrag Radivojac, Lea M. Starita, Douglas M. Fowler, Vikas Pejaver

## Abstract

The utility of clinical genetic testing is limited because around 90% of missense variants in ClinVar remain of uncertain clinical significance. Variant effect predictors (VEPs) can score any missense variant, potentially empowering variant classification. Realizing this potential requires calibration to translate predictor scores into evidence. However, genome-wide calibration ignores predictor heterogeneity across genes, causing evidence misassignment. We developed an automated, data-adaptive framework that optimizes two complementary approaches: gene-specific calibration for genes with enough variants for calibration, and domain-aggregate calibration for other disease-associated genes, which groups variants from protein domains with similar predictor score distributions for calibration. Applied to three predictors across 2,769 genes, this framework assigned evidence to 10.6% more variants on average while generally improving evidence accuracy compared to genome-wide calibration. These calibrations and the resulting calibrated computational evidence are available through the PredictMD portal. Our framework substantially increases the clinical utility of VEPs for variant classification.

## Introduction

Clinical genetic testing has revealed millions of variants, yet approximately 90% of missense variants in ClinVar are variants of uncertain significance (VUS)^1^. VUS represent a major challenge for genomic medicine as they cannot be used to guide medical management and can prolong diagnostic odysseys for patients and their families. VUS result from missing or weak evidence during variant classification, the process by which a variant’s association with disease is determined by collecting, weighting and combining different lines of evidence^2^. These lines of evidence include the patient’s phenotype, segregation of the variant with disease in families, the frequency of the variant in the population, the effect of the variant in experimental assays and the results of computational variant effect predictors (VEPs), among others.

Variant effect predictors are statistical or machine learning models that integrate protein sequence, structure, evolutionary conservation, and other genomic features to predict whether a variant does or does not disrupt function. Because variant effect predictors can predict the outcome of most types of possible single nucleotide variants, they are often the only type of evidence available for most variants. Moreover, predictor accuracy has dramatically increased in recent years as a consequence of more sophisticated models, increased availability of training data, and the adoption of strategies to improve generalizability and reduce circularity and bias^3–6^. Thus, variant effect predictors could play a major role in reducing VUS, enhancing genomic medicine and delivering value to patients.

However, predictors have had limited impact on variant classification because guidelines dictated they could only generate the weakest level of evidence toward benign or pathogenic classifications^2,7^. This restriction reflects longstanding difficulties in variant effect predictor calibration, the process of transforming predictor scores into evidence of pathogenicity or benignity based on their ability to discriminate known pathogenic and benign variants. Ideally, predictions for each gene would be calibrated individually, but currently most genes do not have enough pathogenic or benign ClinVar control variants for robust calibration. Recent genome-wide calibration efforts overcame this limitation through a key innovation: aggregating control pathogenic and benign variants across all disease-associated genes into a single pooled set of control variants. This genome-wide approach provided sufficient power to establish universal score thresholds that were then applied uniformly across genes for evidence assignment, demonstrating for the first time that predictors can provide strong evidence, and thus, initiating a paradigm shift in clinical use of variant effect predictors^8,9^.

Nonetheless, predictors behave variably across genes. Predictors’ score distributions and discriminatory power to discern pathogenic and benign variants vary with protein structure and function, evolutionary constraint, disease mechanism, and inheritance mode. Additionally, genes differentially tolerate amino acid substitutions leading to differing prior (baseline) probabilities of pathogenicity within a gene^10–13^ (**Box 1**). When calibration is performed using globally defined thresholds and a universal prior^8–10^, these sources of heterogeneity lead to systematic over- or under-assignment of evidence for individual variants^10,14^. Addressing this problem requires calibration strategies that (i) account for gene- or domain-specific heterogeneity in score distributions, and (ii) explicitly incorporate gene- or domain-specific prior probabilities of pathogenicity, while being applicable to disease-associated genes^15^ with a limited number of pathogenic and benign control variants.

Here, we introduce a two-pronged framework for more granular calibration than the previous genome-wide approach. First, predictor scores are calibrated for individual genes using fewer variants than needed for genome-wide calibration, while incorporating a gene-specific prior probability of pathogenicity estimated from a semi-supervised machine learning approach^16^. Second, since only ∼3% of disease-associated genes have sufficient control variants to enable gene-specific calibration, we aggregate protein domains with similar predictor score distributions, under the assumption that their control variants would also be similarly distributed. This approach pools control variants across protein domains with similar predictor score distributions, defined using the predicted scores of all possible missense variants within each domain. Domain-specific prior probabilities are estimated using the same approach used for gene-specific priors, enabling appropriate calibration in domains with limited variant counts. This framework overcomes the limitations of genome-wide calibration by enabling gene- and domain-level calibration of variant effect predictors, improving evidence point assignment given the available control variants, and providing a principled foundation for applying computational evidence within the American College of Medical Genetics/Association of Molecular Pathology (ACMG/AMP) guidelines for clinical variant classification.

Using this framework, we performed gene-specific calibration for 132 genes with adequate number of control variants and used domain-aggregate calibration for an additional 2,637 disease-associated genes. We calibrated three top-performing variant effect predictors (REVEL, MutPred2, AlphaMissense) and demonstrated consistent improvements over the genome-wide aggregation approach^6,8,9^. Gene-specific calibration increased the number of variants assigned evidence from a median of 68% to 80% across genes and predictors, improved overall discrimination, and particularly enhanced correct classification of benign variants. For genes lacking sufficient control variants, domain-aggregate calibration increased the number of variants assigned evidence from 68% to 74% and similarly improved performance. All calibrations are publicly available through a new interface for predictive evidence, PredictMD (https://igvf.mavedb.org/), designed to allow clinical users to easily access and evaluate calibrated evidence across genes. Together, these results establish a generalizable calibration framework that overcomes limitations of the genome-wide approach and substantially increases the clinical utility of variant effect predictors for variant classification.

## Box

### Calibration of evidence within the ACMG/AMP framework

The rule-based ACMG/AMP guidelines^2^ for combining population-level, computational, functional, segregation and other lines of evidence to classify a variant as *Pathogenic (P), Likely Pathogenic (LP), Variant of Uncertain Significance (VUS), Benign (B) or Likely Benign (LB)* have been adapted to a probabilistic Bayesian framework^7,17^. This framework depends upon an initial expectation of pathogenicity, the prior probability of pathogenicity, which is updated using the available lines of evidence (**Fig. 1a,b**). The following three quantities are fundamental to this Bayesian variant classification framework:

**Fig. 1:**
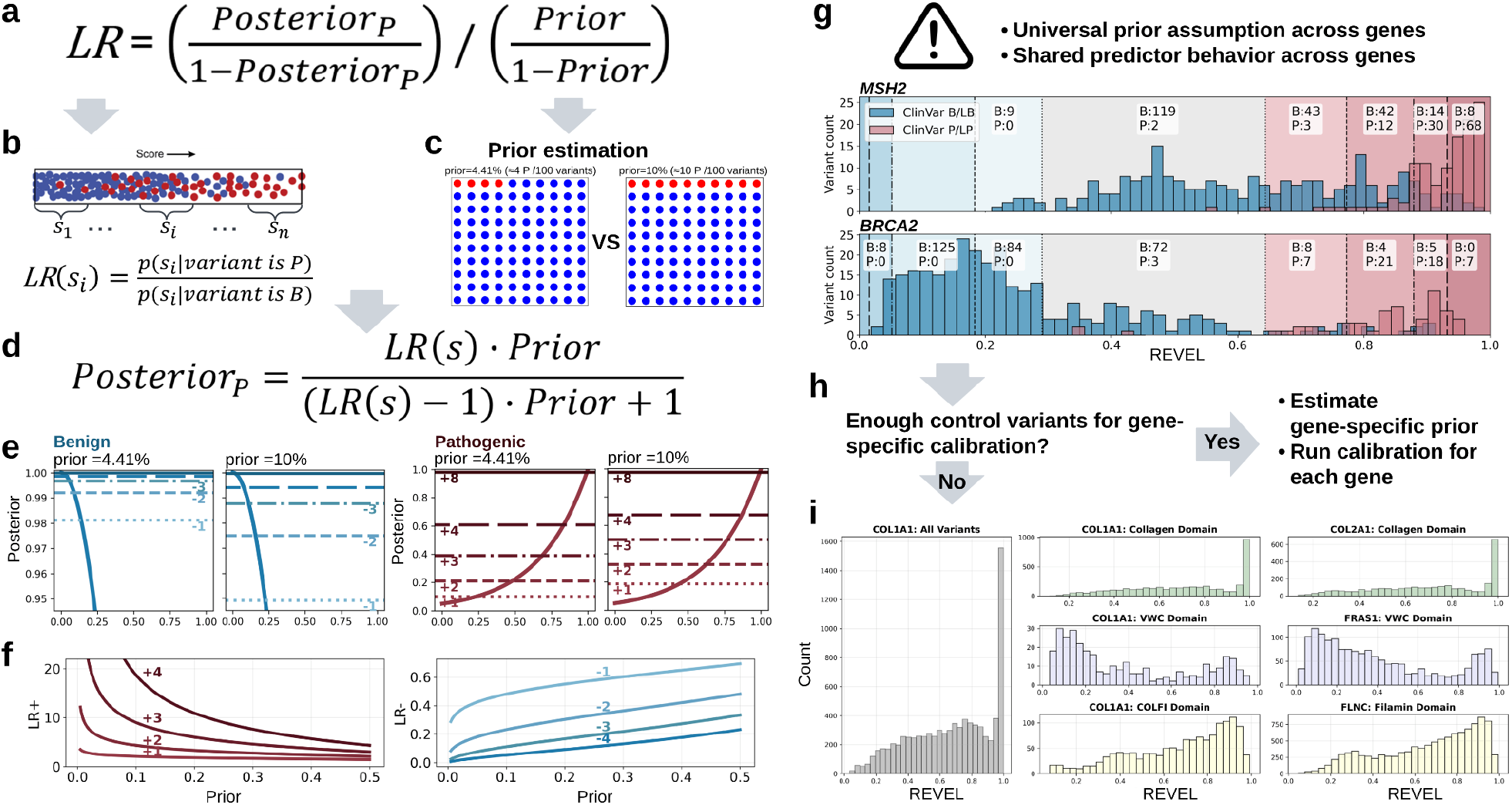
ACMG/AMP Bayesian framework for VEP calibration and limitations of genome-wide aggregation approach and proposed solutions. **(a)** Posterior odds of pathogenicity are obtained by combining **(b)** a score-specific likelihood ratio, *LR*(*s*), with **(c)** a prior probability of pathogenicity. The prior is illustrated as the expected proportion of pathogenic variants among population variants (*e*.*g*., 4.41% versus 10%). **(d)** The posterior probability as a transformation of the formula in (a), **(e)** changing the prior (left, 4.41%; right, 10%) shifts the predictor score ranges corresponding to benign (blue) and pathogenic (red) evidence strengths, demonstrating the direct dependence of evidence thresholds on the prior using simulated data. **(f)** Prior-dependent likelihood ratio (LR) for ACMG/AMP evidence strengths. Left panel shows LR+ as a function of prior for pathogenic evidence strengths (+1 to +4 points); right panel shows LR-as a function of prior for benign evidence strengths (-1 to -4 points). **(g)** Limitations of genome-wide calibration. Applying a universal prior and genome-wide calibration yields identical evidence thresholds across genes despite differences in gene-specific score distributions. As shown for REVEL in *MSH2* and *BRCA2*, the same score corresponds to different posterior probabilities of pathogenicity, resulting in misplaced evidence thresholds that overestimate pathogenic evidence in one gene while underestimating it in the other. **(h)** Proposed gene-specific calibration strategy. Each gene is first evaluated to determine whether sufficient pathogenic and benign control variants are available to support gene-specific calibration. If sufficient controls are available (Yes), gene-specific priors are estimated and calibration is performed specifically for that gene. If sufficient controls are not available (No), calibration is performed on aggregated protein domains. **(i)** Domain-based calibration for genes with limited control variants, in which protein domains with similar predictor score distributions are aggregated across genes to gather enough control variants to enable robust calibration (example shown for *COL1A1* and related domains).

1. The ***prior probability of pathogenicity*** *(prior)* establishes the baseline probability that a variant is pathogenic before any evidence specific to the variant is considered (**Fig. 1c**). Traditionally, a universal prior based on expert judgement has been applied to variants for all genes^7^. However, newer approaches estimate the prior probability for a single gene or group of genes^8,11^.
2. The evidence ***likelihood ratios*** (*LRs*; also referred to as *odds of pathogenicity*^*7,17,18*^ or *positive likelihood ratio^8,9^*) quantify the strength of a specific source of evidence (for example, a particular predictor score) by comparing how often that evidence occurs among pathogenic control variants relative to benign control variants (**Fig. 1b**). Within the ACMG/AMP Bayesian framework, an integer-valued point system from -8 to +8 has been derived to represent evidence strength more conveniently, where points from multiple lines of evidence can be added to reflect the combined evidence strength^17^. Higher positive points indicate stronger pathogenic evidence, and lower negative points indicate stronger benign evidence; for computational evidence, this scale is capped at +4 points (pathogenic) and -4 points (benign)^9,19^. Evidence points can be determined by thresholding *LR* values using the ten *LR*-thresholds for ±1, ±2, ±3, ±4 and ±8 points, defined in the ACMG/AMP Bayesian framework^5,7^. For example, on the pathogenic side, a line of evidence receives +2 points if its *LR* lies between the *LR*-thresholds for +2 and +3 points. Similarly, on the benign side a line of evidence receives -2 points if its *LR* lies between the *LR* -thresholds for -2 and -3 points. The *LR*-thresholds are prior-dependent quantities, as shown in **Fig. 1f**. They are greater than 1 for pathogenic (positive) points and less than 1 for benign (negative) points. In this work we consider two types of *LR* quantities: 1) *LR* assigned to a single variant based on its predictor score (also referred to as local *LR*^5,8^), and 2) Interval *LR*: the *LR* assigned to a group of variants with predictor scores lying in an interval.
3. The ***posterior probability of pathogenicity*** *(posterior)* updates the prior probability of a variant’s pathogenicity, after observing evidence specific to the variant. The posterior may be higher, lower or the same as the prior, depending on how strong the evidence is in favor of the variant’s pathogenicity or benignity. In this work, we consider the posterior probability of a variant as evidenced by its variant effect predictor score.

These quantities are linked through Bayes’ theorem: *LR* is equal to the ratio of the posterior and prior odds of pathogenicity (**Fig. 1a**). Because predictor scores cannot be directly expressed as probabilities or *LRs*, they cannot be used directly in this framework. Instead, the predictor score *s* must be converted to its posterior probability of pathogenicity, *posterior*_*p*_ (*s*), a process known as **calibration** (**Fig. 1e**). Once calibrated, posterior probabilities can be converted back into *LRs* (local *LRs*), which are then compared against the prior-dependent *LR*-thresholds to assign evidence points^7^. The prior plays a critical role in this process. Changing the prior alters the mapping between predictor score and posterior probability, and also alters the *LR* thresholds, and therefore shifts the posterior and score intervals corresponding to each evidence point (**Fig. 1e**). Each *LR* threshold can be mapped to a posterior threshold and a predictor score threshold.

## Results

### Gene-specific calibration is feasible for a substantial number of clinically relevant genes

Calibration begins by establishing a prior probability of pathogenicity for each gene as these influence the selection of score thresholds for ACMG/AMP evidence points. Expert-opinion-based priors are not a scalable solution and tend to be overestimated^8,20^. While data-driven priors derived solely from clinical databases such as ClinVar or biobanks, e.g., All of Us^21^ or the UK Biobank^22^, can be useful, these data tend to be insufficient for prior estimation due to the selection bias of variants (in the case of clinical databases) or the lack of variant pathogenicity status (in the case of biobanks). To overcome these limitations, we adopted DistCurve, a previously established semi-supervised machine learning approach to estimate priors from ClinVar pathogenic or likely pathogenic (P/LP) and variants in the general population using gnomAD variants^8,16,23,24^. DistCurve contrasts known pathogenic variants with variants found in the gnomAD database in a shared, high-dimensional feature space capturing evolutionary constraint, protein structure, and other biologically relevant signals to estimate the proportion of variants in gnomAD that share features with known pathogenic variants^24^ (**Extended Data Fig. 1**).

We estimated priors for 648 genes with at least 10 control P/LP variants. Estimated gene-specific priors ranged from nearly 0% to 32.2% (median 5%; **Extended Data Fig. 2**), revealing substantial heterogeneity across genes (**Supplementary Table 1**). Notably, 360 (55.6%) genes exceeded the genome-wide prior of 4.4%^8^ and 115 (17.8%) genes exceeded the 10% prior threshold proposed in the ACMG/AMP Bayesian framework^7^, underscoring pronounced differences in baseline pathogenic risk across genes, consequent to mutation.

As described above, no gold standard exists for benchmarking gene-specific priors. We compared our prior estimates with two independently generated sources of gene-specific priors: population-based priors derived from UK Biobank data and clinical priors estimated using ClinVar pathogenic and benign variants^11^; **Extended Data Fig. 3**). For genes where pathogenic variants cause readily ascertained phenotypes (e.g., *LDLR* causing hypercholesterolemia, *ATM* increasing cancer risk), our priors showed strong concordance with both population and clinical prior estimates at nearly double the ACMG/AMP default 10%. For other cancer susceptibility genes (*TP53, BRCA1, BRCA2, MLH1, MSH6, MSH2*), our gene-specific priors were largely comparable to either population or clinical prior estimates. For the remaining genes, priors varied markedly across estimation methods, potentially reflecting uncertain estimates from sparse and/or biased data for these conditions. Together, these comparisons highlight both the challenges of prior estimation for individual genes associated with rare Mendelian conditions and the ability of the DistCurve method to generate reasonable gene-specific priors in the absence of robust clinical data.

DistCurve-derived gene-specific prior estimates enabled us to explore calibration of variant effect predictor scores for individual genes. The number of control variants required for robust calibration depends on the prior, the balance between pathogenic and benign control variant classes and the model chosen to fit the control variant distributions. To assess how these factors affect calibration performance and to determine the minimum number of control variants required for each gene, we developed a clinically interpretable miscalibration metric (**Fig. 2a**) that measures deviations between calibrated and true posterior probabilities generated by simulations (**Fig. 2a**, red line vs. blue line). The difference between the calibrated and simulated true posteriors are calculated within score ranges where evidence would be assigned (**Fig. 2a**, shaded region) and is expressed in evidence points^17^. We defined successful calibration as mean miscalibration of at most one evidence point. As expected, the mean miscalibration decreased with increasing numbers of control variants in our simulations (**Fig. 2b**), but varied considerably across the 11 different methods we used to fit control variant distributions. Parametric methods such as Platt scaling^25^ and Beta calibration^26^ performed best when the number of controls were limited. More flexible approaches, particularly neural network–based methods (e.g. MonoPostNN), performed best with more control variants, but still fewer than those required for the local calibration algorithm used in genome-wide calibration. We therefore developed a data-adaptive calibration framework that identifies the best method given the prior and number and balance of control variants rather than relying on a single method for all genes (**Extended Data Figs. 4, 5 and 6**).

**Fig. 2:**
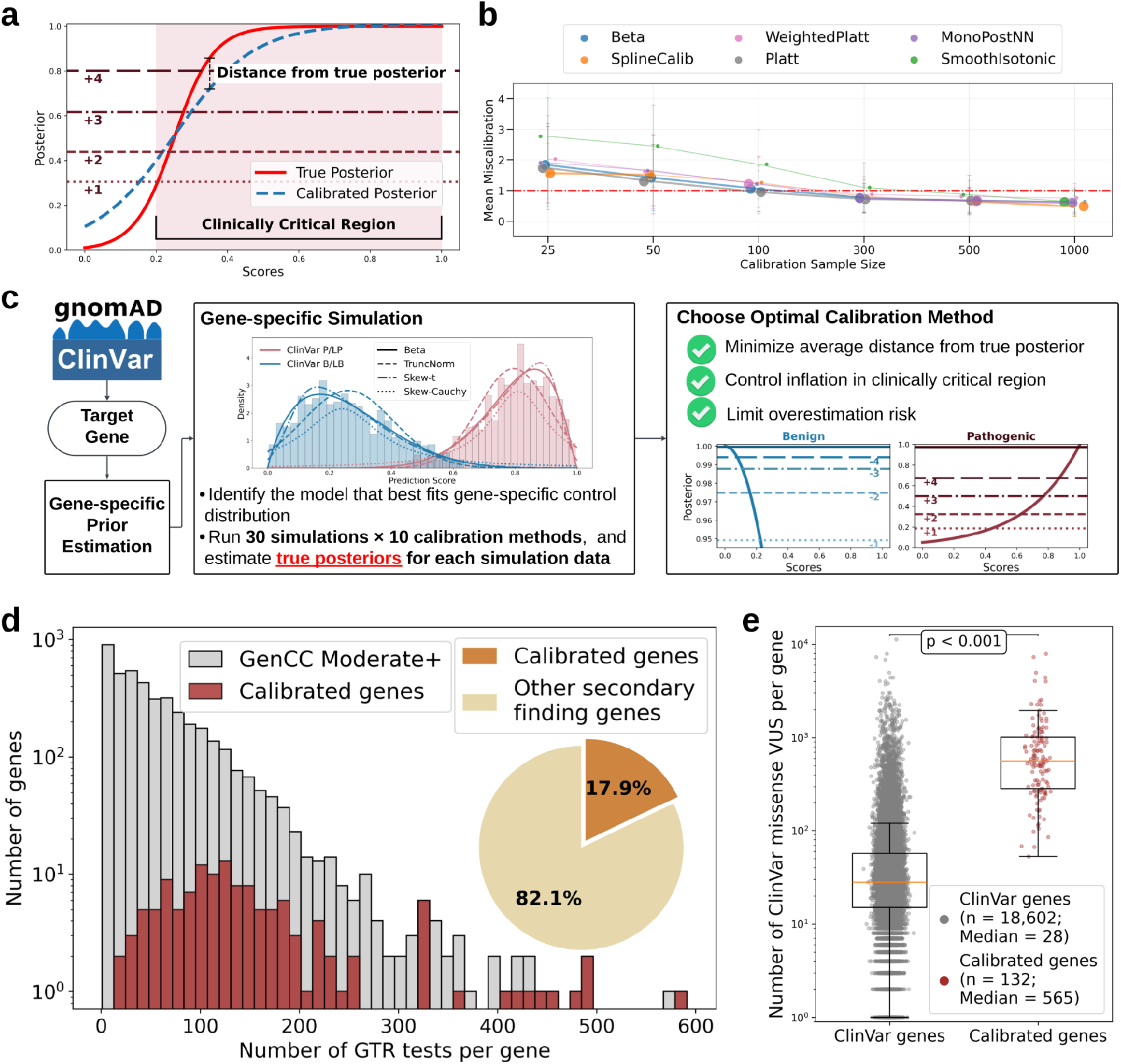
Development of a data-adaptive framework for gene-specific calibration. **(a)** Schematic illustration of the miscalibration metric. Miscalibration is quantified as the vertical deviation between the calibrated posterior (dashed blue) and the true posterior (solid red) within the score range where evidence points would be assigned. **(b)** Mean miscalibration as a function of the number of control variants across simulated datasets. Error bars denote variability across different pathogenic-to-benign ratios and prior probabilities. The top three performing calibration methods (of 11 evaluated) are shown, with enlarged scatter points indicating best method performance at each sample size. **(c)** Schematic of the data-adaptive framework for gene-specific calibration. For each gene, multiple calibration methods are evaluated and the optimal approach is selected based on empirical performance, prioritizing robustness, accuracy, and control of clinically relevant overestimation. The selected method is then used to derive calibrated score thresholds and corresponding evidence points. **(d)** Distribution of Genetic Testing Registry (GTR) test counts per gene, with calibrated genes in red and GenCC moderate+ genes in gray. The inset pie chart shows that calibrated genes account for 17.9% (15/84) of secondary finding genes. **(e)** Distribution of ClinVar missense VUS per gene for all ClinVar genes (gray; excluding calibrated genes, n = 18,602) compared with calibrated genes (red; n = 132). Calibrated genes harbor substantially more VUS per gene (median 565 versus 28; two-sided Mann–Whitney U test, P < 0.001).

In our data-adaptive framework, each gene is calibrated using 11 methods, available control variants^1^, and the DistCurve-derived prior (**Fig. 2c and Supplementary Table 1**). Genes were selected based on the number of available control variants and the balance between pathogenic and benign classes. Applying these criteria, we identified 157 genes suitable for calibration within our framework. We evaluated calibrations from each of the 11 methods for all 157 genes across 3 predictors using a three-step, miscalibration-aware selection procedure. Methods were first filtered to control inflation by excluding those with extreme overestimation in the tail regions of the posterior that would result in too much evidence assigned. Remaining methods were then ranked to minimize the average distance from the true posterior across the full score range using our miscalibration metric (**Fig. 2a**). Finally, residual overestimation risk in the tails was quantified, and the method with the lowest overall miscalibration was selected for each gene. The optimal method was then used to derive calibrated predictor score thresholds and corresponding evidence point assignments. By jointly accounting for gene-specific priors, data availability, and pathogenic-to-benign ratios, this framework enables principled and scalable selection of calibration strategies tailored to individual genes (**Extended Data Figs. 7, 8 and 9**).

Among the 157 candidate genes, we successfully calibrated at least one variant effect predictor for 132 genes. Predictor-specific calibration was achieved for 101 genes with REVEL, 98 genes with AlphaMissense, and 105 genes with MutPred2 (**Extended Data Fig. 10**). Across all calibrated predictors, every gene had variants that were assigned at least one point of benign and pathogenic evidence (±1 point). In addition, a substantial fraction of genes spanned the full range of evidence strengths. Specifically, variants reached the maximum benign evidence strength (−4 points) in 67.3%, 61.2%, and 65.7% of genes calibrated with REVEL, AlphaMissense, and MutPred2, respectively. Maximum pathogenic evidence strength (+4 points) was observed in 55.5%, 54.1%, and 41.0% of calibrated genes for the same predictors (**Extended Data Figs. 11, 12 and 13; Supplementary Table 2**). Both extremes (+4 and −4 points) were achieved in 41.6%, 38.8%, and 29.5% of genes for REVEL, AlphaMissense, and MutPred2, respectively, with REVEL identifying the largest number of such genes (n = 42).

Although these 132 genes represent a small fraction of disease-associated genes (3.1% of 4,297 GenCC moderate or higher confidence genes)^15^, they are enriched for genes with registered genetic tests (8.3%; 20,653 of 248,574 tests; **Fig. 2d**) and represent nearly 18% of secondary-finding genes^27,28^. These genes have also accumulated 20-fold more ClinVar missense VUS than typical disease genes (median 565 vs. 28 per gene; **Fig. 2e**), indicating that gene-specific predictor calibrations have high potential clinical utility to contribute to the resolution of the substantial VUS burden in these genes. Gene-specific calibration results for all 132 genes are available for exploration through PredictMD, an interactive web resource at https://igvf.mavedb.org/.

### Gene-specific calibration improves calibration accuracy and clinical utility of evidence for 132 genes critical to genomic medicine

Since there are no gold-standard computational evidence thresholds to compare against, we used alternative strategies to evaluate the correctness and utility of variant effect predictor score thresholds obtained from gene-specific calibration. First, we qualitatively compared the predictor score intervals defined by gene-specific calibration with genome-wide calibration. We illustrate this through the calibration of REVEL for *MSH2* (**Fig. 3a,b**). Under genome-wide calibration, several score intervals assigned pathogenic evidence failed to show enrichment for pathogenic control variants (**Fig. 3b**, top). In contrast, gene-specific calibration realigned score intervals with the underlying control variant distributions (**Fig. 3a**, bottom).

**Fig. 3:**
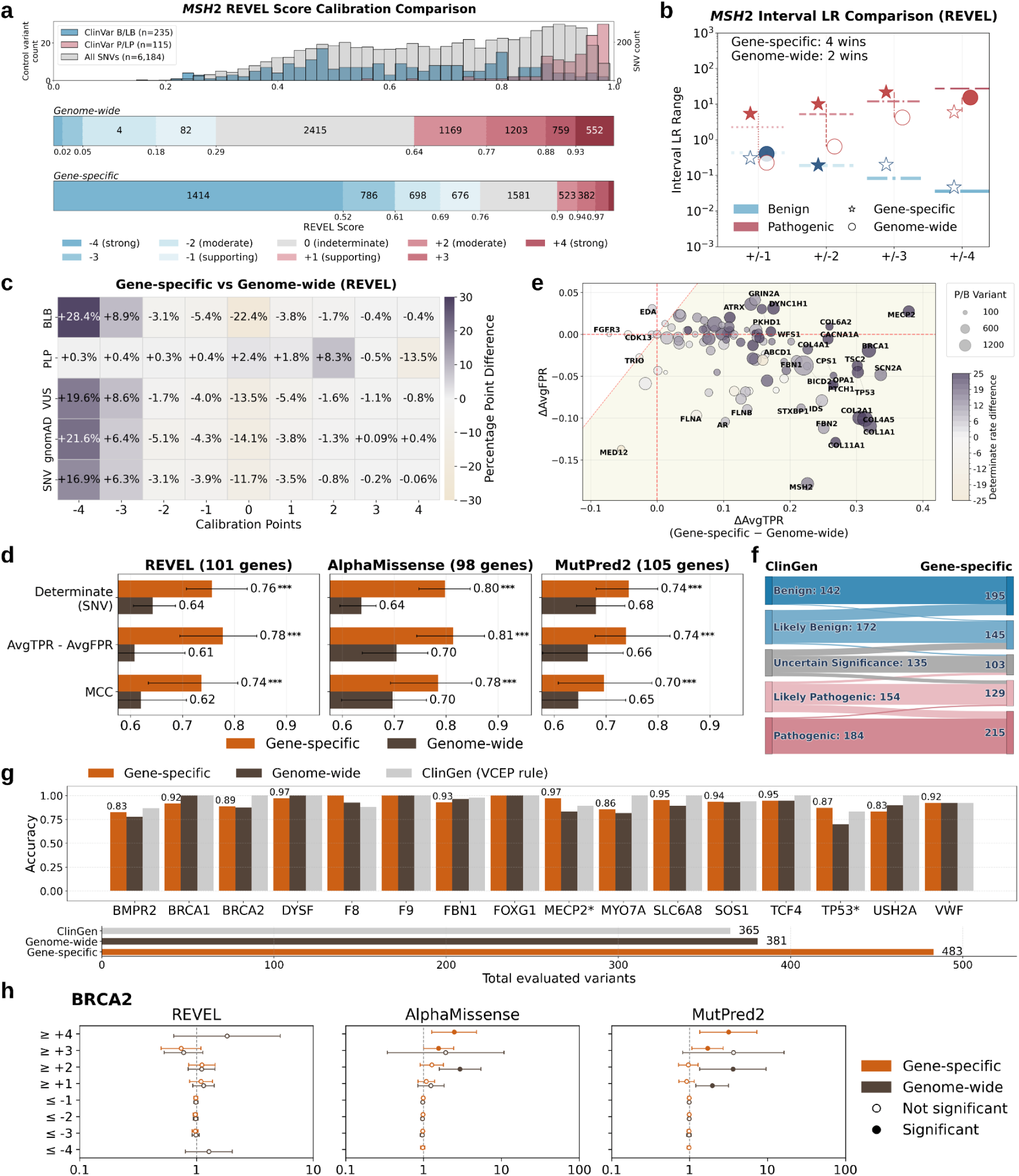
Gene-specific calibration improves accuracy and clinical utility. **(a)** *MSH2* showing improved alignment of pathogenic and benign variant distributions under gene-specific calibration. (Top) Histogram of REVEL scores for ClinVar-classified P/LP (red), B/LB (blue), and all possible SNVs (gray), with SNV counts shown on the right y-axis. (Middle) Stacked bar plot showing score intervals for each evidence strength under genome-wide calibration, with color intensity indicating different evidence strength as indicated. Variant counts in each bin. **(b)** Interval likelihood ratio (*LR*) comparison between gene-specific calibration (star) and genome-wide calibration (circle). Horizontal lines represent expected *LR* cutoffs for the indicated evidence points (x-axis). Each point is colored by evidence type and filled to indicate which method performed better for that evidence strength. A scorecard in the upper-left corner summarizes the number of “wins” for each method. **(c)** Heatmaps display the differences in the percentage of evidence point assignments stratified by variant classification or source. Grey indicates increased assignment rates by the gene-specific calibration method compared to genome-wide calibration. **(d)** Summary barplots of median performance metrics across calibrated genes with error bars indicating the 25th–75th percentiles. Evidence assignment rate (determinates), AvgTPR–AvgFPR, and MCC are compared for the gene-specific and with genome-wide calibration across predictors. **(e)** Gene-level changes in sensitivity and false positive rate following gene-specific calibration. The shaded area indicates increased performance. **(f)** Sankey diagram comparing classifications for 787 variants across 23 genes when computational evidence is reassigned using evidence from gene-specific calibration (right column), vs the original ClinGen classification (left column). **(g)** Evaluation of per-gene classification accuracy on the ClinGen non-circular variant set using REVEL scores under three approaches: ClinGen-provided computational evidence strengths, genome-wide calibration thresholds, and gene-specific calibration thresholds. Top bar plots show the fraction of variants correctly classified relative to ClinGen reference classifications after recomputing variant classifications using only non-computational evidence to maintain non-circularity. Variants assigned zero computational evidence points are excluded from accuracy calculations. Bottom bar plots show the total number of variants assigned non-zero computational evidence points by each method. **(h)** Odds ratios for disease occurrence in the All of Us biobank for variants meeting different evidence strength thresholds in *BRCA2* using gene-specific calibration compared with genome-wide calibration. The x-axis shows the odds ratio (vertical dashed line indicates OR = 1), and the y-axis shows total evidence points for variant sets. Circles represent estimated odds ratios with 95% confidence intervals (whiskers); filled circles indicate statistically significant associations.

To systematically compare performance of calibration approaches (choice of model and genome-wide vs. gene-specific approach) on a given gene, we used the interval-based *LR* (Methods) metric described previously^8,9^. The interval-based *LR* captures the *LR* of a group of variants lying in a given score interval as a single number and can be computed directly from the control variants of a gene without relying on a calibration model, thereby giving an unbiased measure of the gene-specific *LR* for the score interval. To assess the correctness of the score intervals and consequently, the points assigned by a calibration approach, we compare the interval-based *LR* with the corresponding *LR*-threshold, representing the least evidence that a variant in the score interval should have. An ideal interval-based *LR* is expected to be close to the corresponding *LR*-threshold, with a preference towards a conservative point assignment (**Methods**). Calibration approaches are compared by counting the number of wins across the 8 (4 pathogenic and 4 benign) score intervals for each gene. We compared gene-specific and genome-wide calibration approaches in this manner. For *MSH2*, the gene-specific approach “won” in 4 of those 8 intervals (**Fig. 3b**; filled stars). The shift in the score intervals assigned pathogenic and benign evidence by the gene-specific calibration greatly increased the accuracy of calibrated evidence for this gene over genome-wide calibration (93% of control variants assigned evidence in the correct direction vs. 53%, respectively).

Next, we evaluated the accuracy and clinical utility of our calibration framework across all 132 genes and predictors using multiple performance metrics. First we assessed whether the score intervals for each evidence strength from the gene-specific calibration was more or less consistent with the expected likelihood ratios than the score interval from the genome-wide calibration for each calibrated gene and predictor. For REVEL, gene-specific calibration achieved equal or greater interval-level performance for 75 of 101 genes (65 wins and 10 ties), significantly exceeding genome-wide aggregation (binomial test P = 3 × 10^−5^). Similar patterns were observed for MutPred2, with gene-specific calibration favored or tied in 71 of 105 genes (51 wins and 20 ties; P = 0.041), and for AlphaMissense, where gene-specific calibration was favored or tied in 64 of 98 genes (47 wins and 17 ties), although this trend did not reach statistical significance (P = 0.091) (**Supplementary Table 3**). This indicates that, for most genes, gene-specific calibrated score intervals produce likelihood ratios that more closely match the theoretical expectations for each evidence strength, whereas genome-wide intervals tend to over- or under-estimate evidence. Importantly, this assessment is more granular than gene-level summaries, as it evaluates whether likelihood ratios increase monotonically across pathogenic evidence intervals and decrease across benign intervals, reflecting more internally consistent evidence assignment within each gene.

Next we evaluated accuracy using control variants from ClinVar and clinical utility by assessing evidence assigned to ClinVar VUS, gnomAD variants and all possible missense SNVs. Gene-specific calibration substantially improved the concordance with benign control variants, yielding a 29 percentage-point higher proportion of variants assigned concordant benign evidence compared to the domain-aggregate approach. This improvement was driven by a reduction in indeterminate assignment (22 percentage points fewer variants assigned 0 points) and fewer false pathogenic assignments (6.4 percentage points lower) (**Fig. 3c**). Furthermore, among the variants correctly assigned benign evidence by both calibration methods, gene-specific calibration assigned stronger evidence for 67%. For pathogenic variants, gene-specific calibration applied stricter thresholds, reducing evidence strength for 42% of variants while introducing few pathogenic-to-benign misassignments (1.5 percentage points). AlphaMissense and MutPred2 showed similar improvements in accuracy with gene-specific calibration (**Extended Data Fig. 14**). Gene-specific calibration also increased clinical utility by assigning evidence to a greater fraction of ClinVar VUS (+14 percentage points; 9,064 variants), gnomAD variants (+14 percentage points; 32,827 variants), and possible missense SNVs (+12 percentage points; 143,594 variants) (**Fig. 3c, Extended Data Fig. 14**).

Across all three predictors, gene-specific calibration consistently outperformed genome-wide calibration. More variants were assigned at least ±1 point of evidence; for REVEL 76% for all missense SNVs for gene-specific calibration were assigned determinate evidence compared to 64% for the genome-wide calibration (**Fig. 3d**). Gene-specific calibration showed significantly higher discriminative performance by both average true and false positive rate difference (AvgTPR–AvgFPR), and Matthew’s correlation coefficient (MCC). These gains reflect both increased evidence coverage and improved accuracy of evidence assignment.

For each gene, gene-specific calibration generally increased sensitivity, with more variants assigned evidence in the correct direction relative to genome-wide calibration (right half of **Fig. 3e**). These gains were accompanied by lower or only modest increases in the average false positive rate (FPR), defined as the mean of benign- and pathogenic-side FPRs (**Fig. 3e**). 58.4% of genes (59/101) fall in the lower-right quadrant of **Fig. 3e**, representing the most favorable outcome with increased sensitivity and reduced false positive rates. Overall, gene-specific calibration yielded a net improvement, defined as sensitivity gains exceeding any increase in FPR, for 95% of genes (96/101), resulting in more correctly assigned evidence at the gene level (**Fig. 3e and Extended Data Figs. 15, 16, 17, and 18**).

Thus far, we have assessed calibration accuracy using concordance with control variants from ClinVar, but since most variants have been classified using predictive evidence it can render these comparisons circular. To avoid this circularity, we leveraged the ClinGen Evidence Repository, which records all the evidence expert panels use to classify variants, to assess our calibrations. We identified 787 variants in 23 genes that were in our gene-specific calibrated set. Here, we removed the original predictive evidence applied by the expert panel, replaced it with our newly calibrated evidence and recalculated the total points to generate an updated classification. The updated classifications incorporating gene-specific evidence did not cause any P/LP or B/LB variants to cross over to opposite classes (**Fig. 3f**) even though we applied stronger evidence for 18% of variants (143) (**Extended Data Figs. 19, 20 and 21**). Moreover, the newly calibrated evidence from REVEL demonstrated higher clinical utility, reducing the VUS by 24% from 135 to 103 (**Fig. 3f**) and moving variants originally in the likely benign/pathogenic (LB/LP) classes to the more certain benign and pathogenic classes, a trend also observed with AlphaMissense and MutPred2 (**Extended Data Figs. 20 and 21**).

We also used these ClinGen Evidence Repository variants to assess concordance of evidence assigned by gene-specific calibration with variants classified without predictive evidence for specific genes. Here, we identified 16 genes with at least 10 definitively classified variants in the repository, and used the non-computational evidence to recalculate an updated classification. Gene-specific evidence from REVEL showed 83–100% concordance with these non-circular P/LP and B/LB classifications. For 12/16 genes, gene-specific calibration matched or exceeded genome-wide calibration concordance, and for 6/16 genes matched or outperformed predictive evidence assignments provided by expert panels (**Fig. 3g and Extended Data Figs. 22 and 23**). These high concordance rates are particularly notable given that gene-specific calibration assigned evidence for substantially more variants (483) than the genome-wide calibration (381) or expert panels (365).

Finally, we reasoned that a variant’s calibrated evidence strength should correlate with the effect size for its association with disease, as estimated from a large cohort. To this end, using data from the All of Us Research Program’s biobank, we estimated odds ratios (ORs) for variants meeting different evidence-strength thresholds from five genes with well-established disease associations and sufficient data for gene-specific calibration. Correlations were most apparent for *BRCA2* (**Fig. 3h**). For AlphaMissense and MutPred2, ORs at ≥ +3 points exceeded 1 (1.56 and 1.71) and increased further at ≥ +4 points (2.47 and 3.16) for gene-specific calibration, whereas thresholds from genome-wide calibration resulted in significant ORs at lower thresholds (e.g., ≥ +2 points: 2.94 and 3.6) but weaker or inconsistent ones at higher thresholds (e.g., ≥ +3 points: 1.92 and 3.65). For REVEL, neither calibration strategy achieved significant ORs, suggesting the non-correlation to be a consequence of the predictor rather than calibration. For the remaining genes (*BRCA1, MSH2, TP53*, and *TSC2*), trends were heterogeneous (**Extended Data Fig. 24**) and potentially confounded by predictor-specific behavior, the limited number of variants at different thresholds, and challenges in defining high-quality phenotypes. In general, while biobank-based ORs provide a view of how calibrated evidence relates to clinical outcomes, they cannot by themselves classify variants as pathogenic or benign and should be interpreted with caution. Furthermore, for benign variants, ORs are expected to be 1 regardless of evidence strength, limiting the ability of biobank-based analyses to comprehensively assess calibration quality.

### A domain-based approach to aggregate variants is feasible for genes with too few control variants

Because gene-specific calibration is not feasible for most genes, we sought a more granular alternative to genome-wide aggregation. Analysis of predictor score distributions showed that domains within the same gene can exhibit distinct distributional shapes, whereas similar domains across different genes often share comparable distributions (**Fig. 1i**). We therefore hypothesized that domains with similar predictor score distributions would also share similar distributions of pathogenic and benign control variants. To test this hypothesis we developed a domain-aggregate approach that groups protein domains based on the similarity of the distribution of all predictor scores in each domain (**Fig. 4a**).

**Fig. 4:**
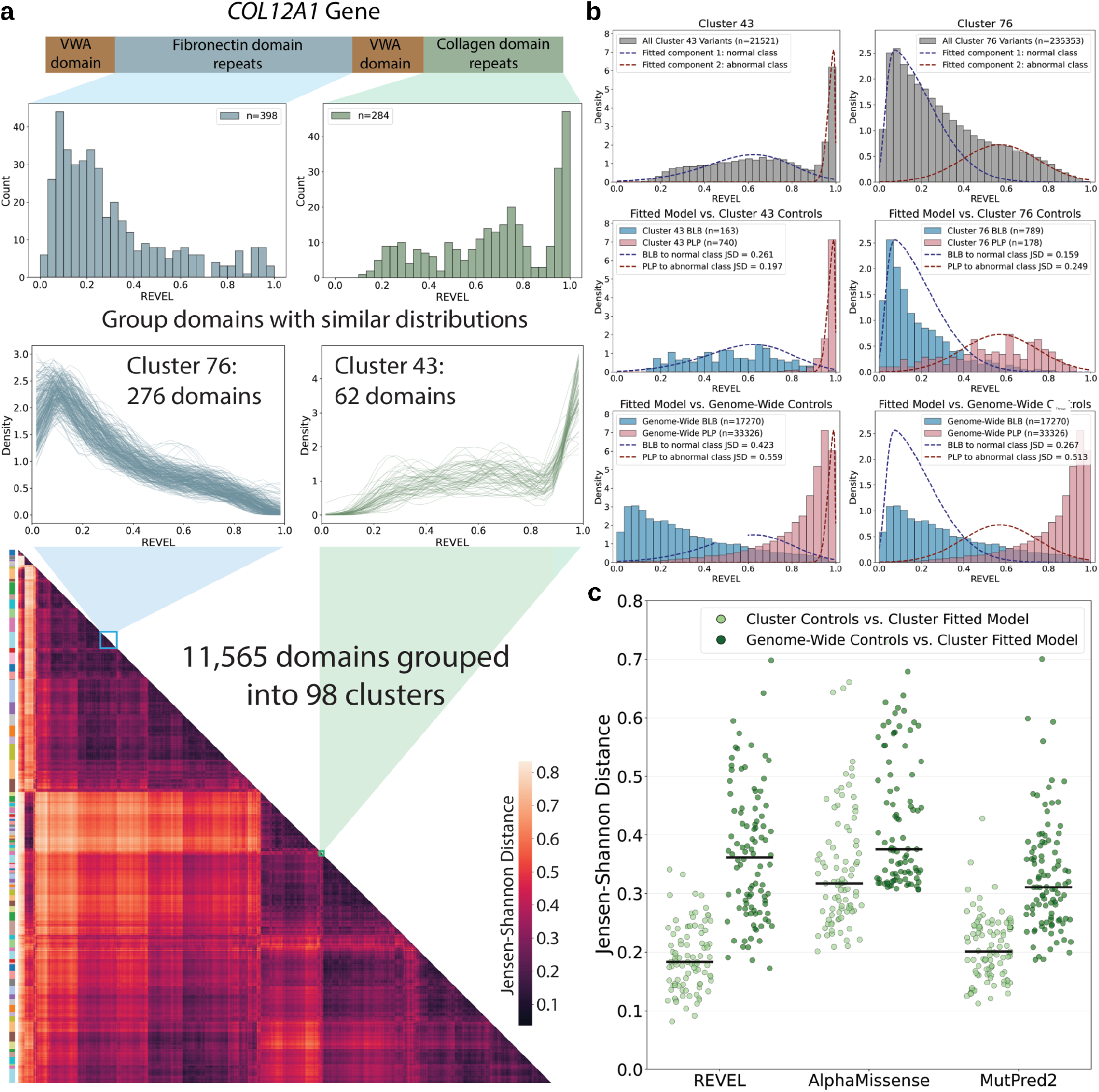
Domain aggregation yields cluster-specific control variant distributions that better match all-variant score distributions. **(a)** Example of domain aggregation for *COL12A1*, showing two clusters with distinct score distributions: fibronectin type III (FN3) domain repeats assigned to cluster 76 and collagen domain repeats assigned to cluster 43. Histograms show the distribution of predictor scores for all variants within an individual repeat of each domain. Density plots show the distribution for each domain grouped into that cluster where each line represents an individual domain. The heatmap below represents the full hierarchical clustering of domain distributions based on Jensen-Shannon distance. The 98 domain clusters are denoted as colored bars on the left of the heatmap. **(b)** For each cluster, a skew-normal mixture model was used, fitting two components to estimate the underlying normal and abnormal score distributions from the all-variant score distribution (top). The fitted normal and abnormal scoredistributions were compared to the cluster-specific control variant distribution (middle) and to the genome-wide control variant distribution (bottom) using the Jensen–Shannon distance. In both clusters 76 and 43, the cluster-specific control variant distributions more closely matched the modeled distributions than the genome-wide control variant distribution. **(c)** Extension of the analysis in (b) across all clusters and predictors. Swarm plots show the difference in the Jensen–Shannon distance measured for the all variant distributions vs cluster-specific control variant distributions and the genome-wide control variant distributions. Black lines indicate the median.

We identified 11,565 domains in the 2,753 genes with moderate or greater evidence for gene-disease association^15^ and at least one pathogenic or likely pathogenic missense variant in ClinVar^1,29,30^. Approximately half of all possible missense SNVs in these genes are in a domain (**Supplementary Table 4**). Clustering of domains based on the shape of the predictor score distribution for all missense SNVs yielded 98 clusters for REVEL, 92 for AlphaMissense, and 102 for MutPred2 (**Fig. 4a and Extended Data Fig. 25)**. Variants outside annotated domains were assigned to a single “non-domain” variant set.

We first evaluated the appropriateness of our clustering approach by quantifying how well the distribution of scores for all variants within each cluster matched the distribution of scores for labeled ClinVar variants within the cluster, compared to the labeled ClinVar variants from all genes. For example, fibronectin domain repeats in the *COL12A1* gene were assigned to cluster 76, which contained 276 domains with left-shifted score distributions. *COL12A1* collagen domain repeats were assigned to cluster 43, which contained 62 domains with right-shifted score distributions (**Fig. 4a**). From the all-variant distribution for each cluster, we fit a two-component mixture model and refer to component one as the normal class and component two as the abnormal class(**Fig. 4b, top**). We then computed the Jensen-Shannon distance between the modeled distributions and both the cluster-specific (**Fig. 4b**, middle) and genome-wide control variant distributions (**Fig. 4b**, bottom). In clusters 76 and 43, the cluster-specific control variant distributions more closely matched the modeled components representing the normal and abnormal classes than the genome-wide control variant distributions. We extended this analysis across all clusters from the three predictors, and the cluster-specific control variant distributions more closely matched the cluster-specific all-variant distribution than the genome-wide control variant distribution consistently (**Fig. 4c**). Thus, clustering domains based on their all-variant distribution yields aggregated sets of control variants whose distribution is similar.

### Aggregating by protein domains improves calibration accuracy across the clinical genome

We evaluated whether these domain-aggregate control variant sets would increase calibration accuracy compared to the previous genome-wide aggregation approach. We applied the adapted DistCurve algorithm to clusters, yielding priors for 91 of 98 clusters ranging from 1.1-15.8% for REVEL, 88 of 92 cluster for AlphaMissense (range: 0-10.9%) and 100 of 102 clusters MutPred2 (range: 0.28-15.7%) (**Supplementary Table 5**). Clusters where priors could not be estimated contained fewer than 10 P/LP control variants. For clusters with priors, we applied our data-adaptive framework (**Fig. 2c**), comparing 11 candidate calibration methods per cluster. Using the best-performing method for each cluster, we derived thresholds for 78 clusters for REVEL (2,647 genes, 95% clustered variants), 81 clusters for AlphaMissense (2,662 genes, 98% clustered variants), and 88 clusters for MutPred2 (2,637 genes, 97% clustered variants) (**Supplementary Table 6**). The domain-aggregate calibration results are available for exploration in the web resource, PredictMD (https://igvf.mavedb.org/).

To illustrate the comparison between domain-aggregate and genome-wide calibration, we highlight *COL12A1* where variants in the fibronectin domains were primarily in cluster 76 and those in the collagen domains were in cluster 43 (**Fig. 5a,b**). Predictor scores in cluster 76 are left-shifted compared to the genome-wide all-SNV distribution and, thus both pathogenic and benign control distributions are also left shifted (789 B/LB vs. 178 P/LP). Accordingly, calibration of this cluster yielded left-shifted thresholds for both pathogenic and benign evidence. While the indeterminate score interval was slightly smaller for cluster 76 relative to genome-wide calibration, more variants receive indeterminate evidence due to the left-shift of the all-variant distribution compared to genome-wide calibration. In contrast, predictor scores in cluster 43 are strongly right-shifted and both pathogenic and benign control distributions are also right-shifted (740 P/LP vs. 163 B/LB), with pathogenic variants mostly scoring above 0.8. Calibration of cluster 43 produced a smaller indeterminate region that better matched the score distribution relative to genome-wide calibration, substantially reducing the number of SNVs receiving no evidence (1,445 vs. 7,707 under genome-wide calibration). These examples highlight how domain-aggregate calibration yields evidence thresholds based on cluster-specific predictor score distributions, in contrast to uniform, genome-wide thresholding.

**Fig. 5:**
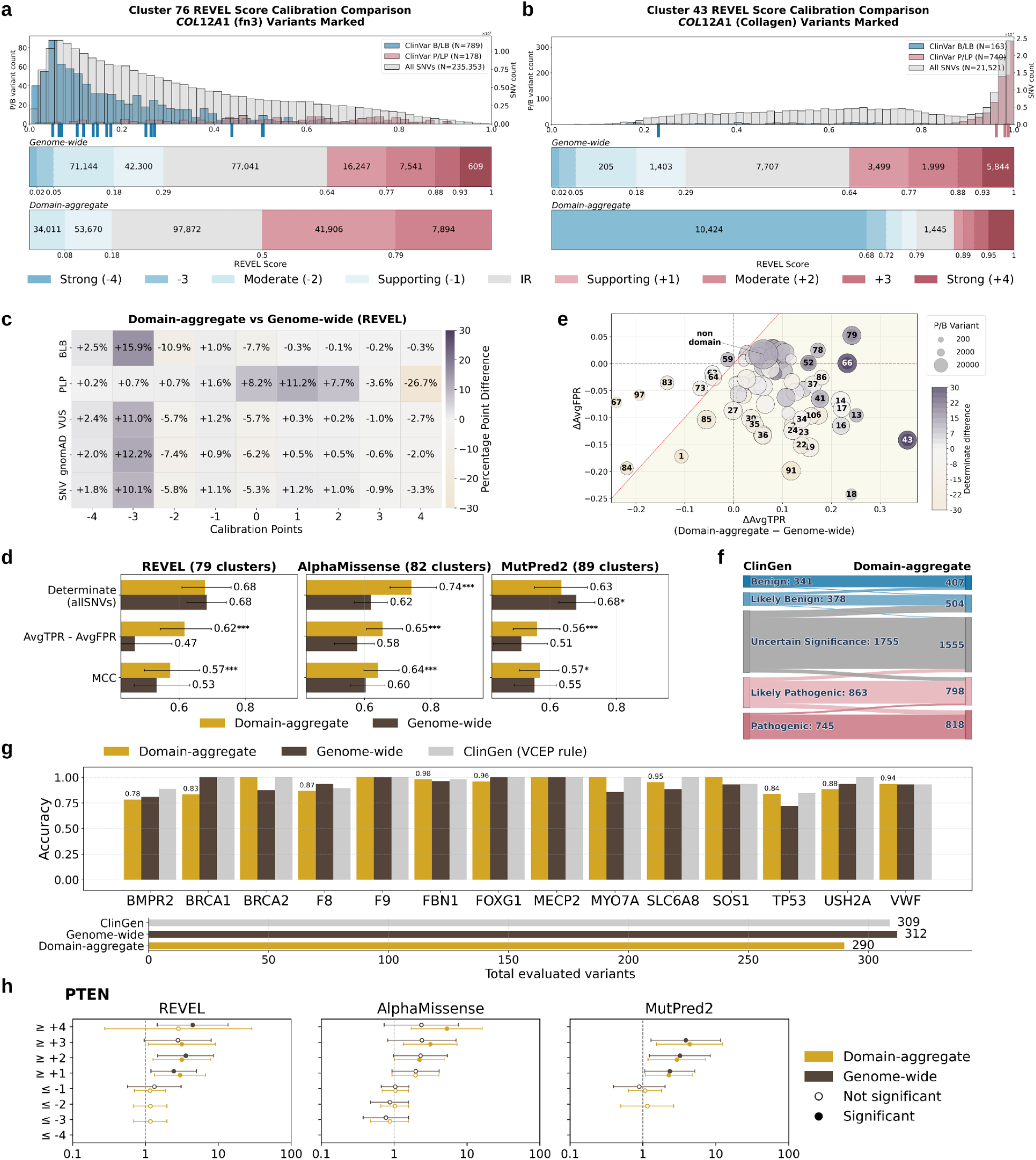
Domain-aggregate calibration performance. **(a)** Example of domain-aggregate calibration for cluster 76 which contains *COL12A1* FN3 domain. **(b)** Corresponding calibration for cluster 43 which contains *COL12A1* Collagen domain. In both panels, the top row shows the cluster-specific predictor score distribution. The middle row shows genome-wide calibration thresholds, and the bottom row shows domain-aggregation calibration thresholds. The number of variants in that interval are indicated. Red shades denote increasing pathogenic evidence strength and blue shades denote increasing benign evidence strength as indicated. Tick marks below the x-axis indicate the locations of *COL12A1* control variants. **(c)** A heatmap of comparing evidence assigned by the domain-aggregate vs. genome-wide calibration across 2,753 genes stratified evidence points assigned (x-axis) and source of variants (y-axis). Grey vs. yellow shading indicates an increased or decreased percentage of variants, respectively. **(d)** Summary barplots of median performance metrics across calibrated clusters with error bars indicating the 25th–75th percentiles. Evidence assignment rate (determinates), AvgTPR–AvgFPR, and MCC are compared for the domain-aggregate and genome-wide calibration across predictors. **(e)** Cluster-level changes in sensitivity and false positive rate following domain-aggregate calibration. The shaded area indicates increased performance. **(f)** Sankey diagram comparing classifications for 4082 variants across 106 genes when computational evidence is reassigned using evidence from domain-aggregate calibration (right column), vs the original ClinGen classification (left column). **(g)** Noncircular evaluation of the per-gene accuracy for three calibration methods: ClinGen-provided computational evidence points, genome-wide calibration thresholds, and domain-aggregate calibration thresholds as indicated. Top bar plots show the fraction of variants correctly classified relative to ClinGen reference classifications after recomputing variant classifications using only non-computational evidence points to ensure noncircularity. Accuracy reflects concordance between each calibration method’s assigned computational evidence and the recalculated classification. Bottom bar plots show the total number of variants assigned non-zero computational evidence points by each method. **(h)** Odds ratios for disease occurrence in the All of Us biobank for variants meeting different evidence strength thresholds in *PTEN* using domain-aggregate calibration compared with genome-wide calibration. The x-axis shows the odds ratio (vertical dashed line indicates OR = 1), and the y-axis shows total evidence points for variant sets. Circles represent estimated odds ratios with 95% confidence intervals (whiskers); filled circles indicate statistically significant associations.

We next evaluated domain-aggregate calibration by comparing evidence assignments for ClinVar control variants with those from genome-wide calibration across all domains (**Fig 5c**). For REVEL, domain aggregation increased the fraction of B/LB variants assigned benign evidence (−1 to −4 points) by 8.5 percentage points (3,781 variants), with larger gains at stronger evidence levels (−3 points: +16 percentage points; −4 points: +2.5 percentage points). Misassignment of pathogenic evidence decreased (−0.9 percentage points; 385 variants), as did indeterminate assignments (−7.7 percentage points; 3,396 variants). ClinVar VUS showed a similar shift, with more variants assigned benign evidence (+8.9 percentage points) and fewer assigned either zero (−5.7 percentage points) or pathogenic evidence (−3.2 percentage points). In contrast, among P/LP variants, the fraction assigned pathogenic evidence decreased (−11.4 percentage points; 4,434 variants), largely reflecting fewer strong evidence assignments (+4 points: −26.7 percentage points; 10,401 variants), accompanied by increases in indeterminate (+8.2 percentage points) and benign (+3.2 percentage points) assignments. Similar trends were observed for AlphaMissense and MutPred2 (**Extended Data Fig. 26**), consistent with gene-specific calibration.

To better understand how the evidence assigned by our domain-aggregate approach differed from evidence assigned by the genome-wide calibration, we analyzed overall performance across clusters for all three predictors (**Fig. 5d**). Evidence coverage varied by predictor, with AlphaMissense clusters having higher determinate evidence coverage than genome-wide calibration (median 0.74 vs. 0.62), REVEL clusters having comparable coverage (median 0.678 vs. 0.683), and MutPred2 clusters having slightly lower coverage (median 0.63 vs. 0.68). However, domain aggregation consistently improved discriminative performance relative to genome-wide aggregation, as measured by higher AvgTPR–AvgFPR, though effect sizes varied across clusters. At the cluster level, median AvgTPR–AvgFPR values were higher under domain aggregation, with paired Wilcoxon tests confirming statistically significant improvements (p <0.001). Similar improvements were observed for MCC.

Using the same TPR–FPR framework as the gene-specific calibration evaluation, we next evaluated cluster-wise calibration performance for domain aggregation by comparing changes relative to genome-wide aggregation (**Fig. 5e**; **Extended Data Figs. 27**). For REVEL, 42 of 79 clusters had simultaneous increases in sensitivity and decreases in false positive rate (**Fig. 5e**, lower right quadrant). Among the remaining clusters with increased FPR, most also showed increased TPR, and all exhibited higher determinate evidence coverage, reflecting net sensitivity gains that outweighed the accompanying increase in false positive assignments. Only eight clusters failed to show a net improvement (above the diagonal in **Fig. 5e**); these clusters were characterized by weaker predictor discrimination (cluster-level AUROC < 0.85), corresponding to domains where REVEL has limited separation between pathogenic and benign variants (**Extended Data Figs. 28, 29 and 30**). The aggregated non-domain cluster showed a modest overall improvement (ΔTPR = 0.06; ΔFPR = 0.017), accompanied by increased evidence coverage.

Using the same non-circular, ClinGen-based evaluation framework as the gene-specific calibration (**Fig. 3f**), we assessed the clinical impact of domain-aggregate calibration by substituting expert-applied predictive evidence with domain-aggregate calibration evidence. Domain-aggregate calibration provided evidence for 4,082 variants across 106 genes in the ClinGen Evidence Repository. Variants originally classified as B/LB or P/LP remained largely stable after application of domain-aggregate calibration evidence, reflecting preservation of strongly supported clinical assertions. VUS were reduced by 11.4% (1,755 to 1,555), and no direct benign–pathogenic reversals were observed (**Fig. 5f**). Most transitions occurred between adjacent ACMG/AMP classes, generally toward more definitive interpretations (e.g., likely pathogenic to pathogenic), with 12.3% of variants (504) receiving stronger calibrated evidence (**Extended Data Fig. 31**). Similar patterns were seen across REVEL, AlphaMissense, and MutPred2 (**Extended Data Figs. 32 and 33**). Together, these results indicate that domain-based calibration improves resolution at classification boundaries.

Next, we assessed concordance of evidence assigned by domain-aggregate calibration to variants in the ClinGen Evidence Repository that could be definitively classified without predictor evidence. These variants correspond to the non-circular P/LP and B/LB classifications described in the gene-specific calibration evaluation section, i.e., variants that did not originally include computational evidence at all. We compared per-gene evidence concordance for computational evidence assigned using domain-aggregate calibration, genome-wide calibration, and predictor evidence using ClinGen VCEP rules across 14 curated disease genes previously evaluated in gene-specific calibration (**Fig. 5g**). Overall, domain-aggregate and genome-wide calibration performed comparably. Domain aggregation improved accuracy relative to genome-wide aggregation for 7 of 14 genes and relative to ClinGen VCEP rules for 3 of 14 genes (**Extended Data Figs. 34 and 35**).

We performed the biobank analysis using data from the All of Us Research Program as before, expanding to 17 gene-disease pairs with previously established case-control cohorts^31^. *PTEN* emerged as an exemplar gene, with domain-aggregate calibration producing clear monotonic increases in OR for AlphaMissense and MutPred2 (**Fig. 5h**), and all thresholds reaching statistical significance (e.g., AlphaMissense ≥ +2: OR = 2.21, ≥ +3: OR = 3.12, ≥ +4: OR = 5.26). On the other hand, genome-wide calibration showed more variable and non-significant patterns at higher thresholds (e.g., AlphaMissense ≥ +2: OR = 2.30, ≥ +3: OR = 2.40, ≥ +4: OR = 2.36). For REVEL, domain-aggregate calibration showed patterns broadly comparable to genome-wide calibration on the pathogenic side (e.g., ≥ +1: OR = 2.98 vs. 2.45; ≥ +2: OR = 3.15 vs. 3.55). For the remaining genes, trends were inconsistent, owing to the aforementioned caveats on using biobank-based analyses to gauge calibration quality (**Extended Data Fig. 36**).

### Direct comparison of gene-specific and domain-aggregate calibration approaches provides insights into clinical use

Given the improvements of both gene-specific and domain-aggregate calibration over the genome-wide approach, we next compared the gene-specific and domain-aggregate approaches with each other, on genes with enough control variants for both calibration strategies (**Fig. 6a-d**).

**Fig. 6:**
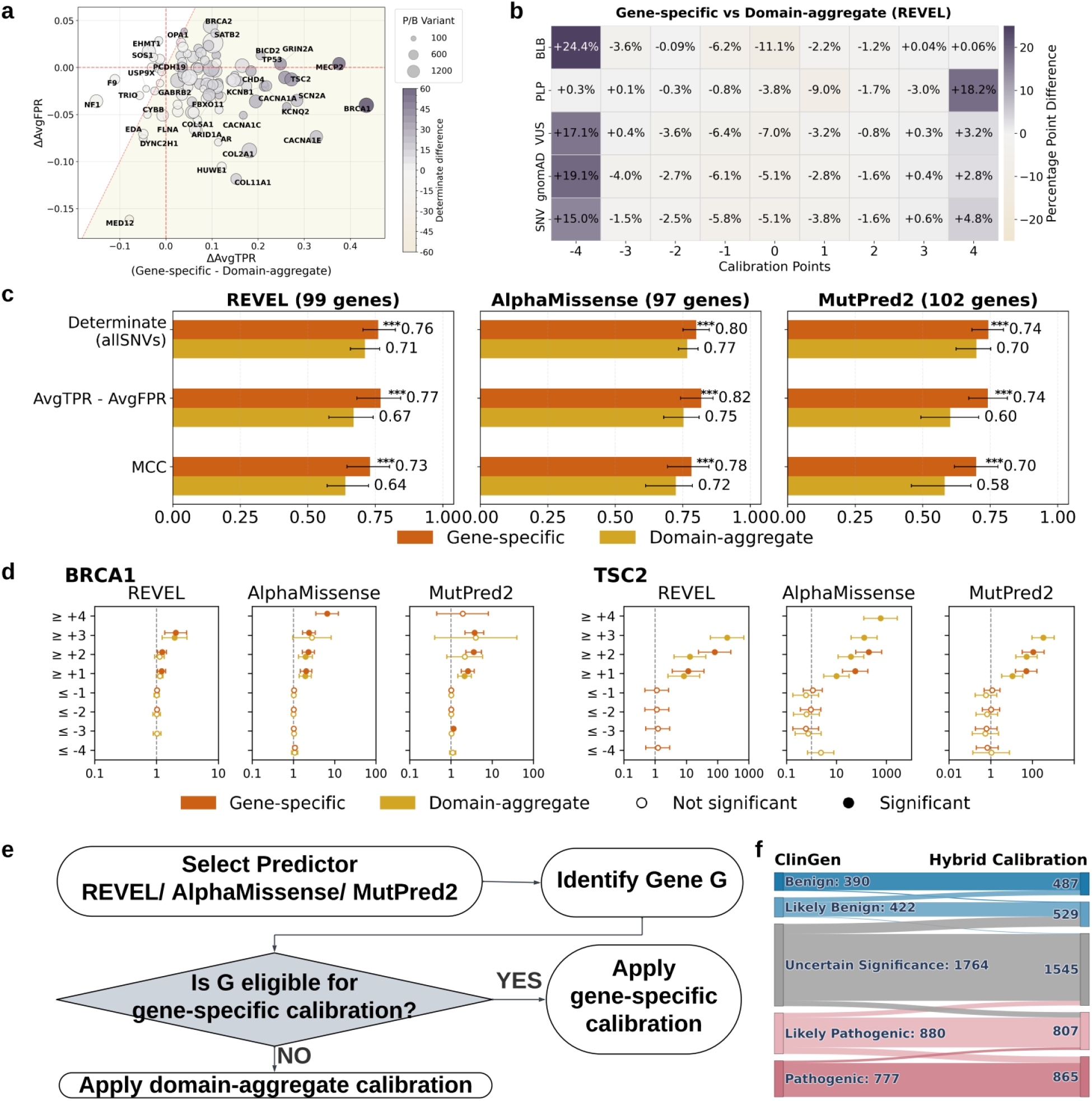
Comparative performance of gene-specific and domain-aggregate calibration. **(a)** Gene-level comparison of average ΔTPR (sensitivity) versus ΔFPR (false positive rate) between gene-specific and domain-aggregate calibration for REVEL. Each point represents a gene, with size proportional to the number of control variants and color intensity indicating percentage difference in evidence coverage. The shaded area below the diagonal dashed line highlights genes benefiting from gene-specific calibration (higher sensitivity with lower false positive rate). **(b)** A heatmap of comparing evidence assigned by the gene-specific vs. domain-aggregate calibration across 97 genes stratified evidence points assigned (x-axis) and source of variants (y-axis). Grey vs. yellow shading indicates an increased or decreased percentage of variants, respectively. **(c)** Summary barplots of median performance metrics across calibrated genes with error bars indicating the 25th–75th percentiles. Evidence assignment rate (determinates), AvgTPR–AvgFPR, and MCC are compared for the gene-specific and domain-aggregate calibration across predictors. **(d)** Odds ratios for disease occurrence in the All of Us biobank for variants meeting different evidence strength thresholds in example genes. *BRCA1* (left) and *TSC2* (right) use gene-specific calibration compared with domain-aggregate calibration. The x-axis shows the odds ratio (vertical dashed line indicates OR = 1), and the y-axis shows total evidence points for variant sets. Circles represent estimated odds ratios with 95% confidence intervals (whiskers); filled circles indicate statistically significant associations. **(e)** Hybrid calibration framework. For a target gene and predictor, genes eligible for gene-specific calibration are assigned gene-specific thresholds; otherwise, domain-aggregate thresholds are applied. **(f)** Sankey diagram comparing classifications for 4233 variants across 107 genes (23 using gene-specific calibration; 84 using domain-aggregate calibration) when computational evidence is reassigned using evidence from hybrid calibration (right column), vs the original ClinGen classification (left column).

Gene-level comparisons of sensitivity and false positive rate demonstrated that gene-specific calibration consistently outperformed domain-aggregate calibration, yielding larger gains in sensitivity for comparable or smaller increases in false positive rate (**Fig. 6a**; **Extended Data Fig. 37**). Among the 97 genes analyzed, gene-specific calibration simultaneously increased sensitivity and decreased false positive rates (the ideal calibration outcome) for 54 genes. Overall, gene-specific calibration showed superior performance for the majority of genes when compared with domain-aggregate calibration (86 of 97 genes for REVEL; shaded region in **Fig. 6a**). While similar trends were observed across the three variant effect predictors, the number of genes benefiting from gene-specific calibration across both metrics differed slightly: 75 of 97 for AlphaMissense and 89 of 102 for MutPred2.

When considering the assignment of evidence to different groups of variants, gene-specific calibration tended to correctly assign evidence on the extreme ends of the points scale (**Fig. 6b**; **Extended Data Fig. 38**). B/LB variants at −4 points increased by 24.4%, and P/LP variants at +4 points increased by 18.2%. This behavior extended to gnomAD, ClinVar VUS and all SNVs as well, suggesting that gene-specific calibration generally assigned higher evidence points in the correct direction than domain-specific calibration. Net misassignments of P/LP variants to negative points and B/LB variants to positive points were also lower for gene-specific calibration (-3.3 and -0.7 percentage points, respectively). Overall, approximately 5% fewer SNVs were assigned 0 points under gene-specific calibration.

Next, we compared standard evaluation metrics between these two approaches (**Fig. 6c**). Gene-specific calibration consistently achieved better performance across all three predictors, with Matthews correlation coefficients (MCC) ranging from 0.70 to 0.78 and improvements in the average true positive rate minus average false positive rate (AvgTPR - AvgFPR) of 0.74 to 0.82. These results indicate that, when sufficient control variants are available, gene-specific calibration provides a more accurate and discriminative framework than domain-level aggregation.

We next compared gene-specific and domain-aggregate calibration using biobank data from the All of Us Research Program, estimating odds ratios (ORs) for variants meeting increasing evidence-strength thresholds as before. Representative genes included *BRCA1* and *TSC2* across three predictors (**Fig. 6d**). For *BRCA1*, gene-specific calibration produced trends broadly similar to domain aggregation for REVEL and AlphaMissense but yielded higher ORs at stronger pathogenic thresholds and more statistically significant estimates (e.g., REVEL ≥ +3: OR = 2.05 vs. 1.96; AlphaMissense ≥ +4: OR = 6.50 for gene-specific vs. OR = 2.81 at ≥ +3 for domain-aggregate). For MutPred2, gene-specific calibration showed stronger significance at higher thresholds (e.g., ≥ +2: OR = 3.55 vs. 2.15; ≥ +3 OR = 3.72, reaching significance) and a clearer monotonic increase across thresholds (≥ +1: OR = 2.57, ≥ +2 OR = 3.55, ≥ +3: OR = 3.72).

In contrast, for *TSC2*, domain-aggregate calibration reached higher pathogenic thresholds and produced much larger ORs, reflecting stronger enrichment for predicted pathogenic variants. Gene-specific calibration assigned variants only up to ≥ +2 (e.g., AlphaMissense ≥ +1: OR = 56.21; ≥ +2: OR = 167.65), whereas domain-aggregate calibration extended to ≥ +4 with even higher ORs (≥ +1: OR = 10.01; ≥ +2: OR = 38.03; ≥ +3: OR = 129.45; ≥ +4: OR = 585.68).

However, gene-specific calibration produced more consistent behavior on the benign side, with ORs generally at or below 1, whereas domain-aggregate calibration showed occasional anomalies (e.g., AlphaMissense ≤ −4: OR = 2.36). There were three other genes for which both calibration approaches could be applied and biobank-based analyses were feasible (*BRCA2, MSH2* and *TP53)*. However, no consistent advantage for either calibration strategy was observed, with gene- and predictor-specific differences precluding any generalizable conclusions (**Extended Data Fig. 39**).

Based on the comparative performance of the three calibration strategies, we devised a hybrid framework that prioritized gene-specific calibration for genes with sufficient control variants to support reliable gene-level modeling and applied domain-aggregate calibration to the remaining genes (**Fig. 6e**). Using this hybrid framework as the calibration strategy, we re-evaluated classifications relative to the ClinGen baseline for 4233 variants (**Fig. 6f**) using evidence assigned by gene-specific calibrations where available (23 genes) and domain-based calibration for the remainder (84 genes) (**Supplementary Table 8**). Compared with the previous ClinGen reclassification analysis using domain-aggregate calibration alone (**Fig. 5f**), the hybrid framework calibrated more ClinGen variants with assigned evidence with increased VUS resolution rate from 11.4% to 12.4%. Among the 62.3% (1348/2163) of variants where both ClinGen and our hybrid calibration framework assigned evidence points in the same correct direction, 66.6% received higher evidence points from the hybrid calibration framework than the evidence used in the original ClinGen classification (**Extended Data Figs. 40, 41 and 42**). Scaling genome-wide, the hybrid framework increased evidence assignment by 8.53% for REVEL and 23.55% for AlphaMissense, while remaining comparable for MutPred2 (−0.29%), across over 12 million SNVs. This corresponded to improved or comparable accuracy (REVEL: 0.73 to 0.75; AlphaMissense: 0.71 to 0.78; MutPred2: 0.74 to 0.71), indicating that the hybrid strategy increases evidence yield, most notably for AlphaMissense, while generally maintaining or improving classification accuracy.

Overall, these results demonstrate that prioritizing gene-specific calibration when sufficient data are available, while using domain-aggregate calibration for genes with too few control variants, provides a practical and more accurate hybrid framework for assigning computational evidence compared with genome-wide calibration alone.

## Discussion

This study makes several key contributions. Our data-adaptive framework selects an optimal calibration method from a breadth of parametric and non-parametric methods that is more robust to the number and balance of available control variants than the local calibration algorithm used for genome-wide calibration. We not only used this for gene-specific calibration but also combined it with a new automated domain-based aggregation strategy that enables calibration even when only a few control variants are available for a gene. This aggregation strategy better accounts for differences in variant effect predictor behavior at more granular levels, including within-gene differences, while ensuring enough power to ensure reliable calibration. We also proposed rigorous metrics and evaluation strategies, aligned with clinical guidelines and applications, to assess the accuracy, utility and robustness of gene-specific and domain-aggregate calibration and compare them with genome-wide calibration. Together, these improve the accuracy of calibration, increase the number of variants assigned computational evidence, resolve hundreds of VUS in the ClinGen Evidence Repository, and reveal predictor performance differences at the gene and domain level.

Recent studies have proposed various alternatives to genome-wide calibration^10,32^. Among these, acmgscaler is the only fine-grained calibration approach that is designed for clinical variant classification. This approach uses kernel density estimation (KDE) to estimate probability densities for pathogenic and benign variants separately and then combines them to calculate likelihood ratios for gene-specific calibration on predictor scores. Our data-adaptive framework differs from acmgscaler in that it directly estimates posterior probabilities or density ratios, which is statistically preferable to separately estimating densities and then applying Bayes’ theorem, as the latter introduces unnecessary estimation error^33^, especially in the tails of predictor score distributions. Furthermore, our framework does not assume that a single calibration method will be optimal for different variant effect predictors and genes and, thus, is suited to a wider range of predictor score distributions and can easily be extended to include acmgscaler or the underlying KDE algorithm as a candidate method assessed during optimal method selection. Nonetheless, we compared our gene-specific calibration approach with acmgscaler in interval-based LR evaluations with gene-specific priors on the set of 132 genes with enough control variants. Our gene-specific calibration approach had significantly more ‘wins’ than acmgscaler (REVEL: 55 of 101 genes; AlphaMissense: 69 of 98; MutPred2: 63 of 105; all binomial P ≤ 0.006; **Supplementary Table 9**).

Gene-specific calibration resolves previously-highlighted discrepancies in calibration between genes^14^ and achieves higher accuracy than genome-wide calibration and, for genes with enough control variants, domain-aggregate calibration. Variant classification and curation efforts such as those by the Variant Curation Expert Panels (VCEPs) within ClinGen are typically organized around specific genes, or hereditary conditions^34–36^. The ClinGen Computational Working Group has recommended the use of genome-wide calibration to guide VCEPs in variant effect predictor selection and score thresholding, while endorsing gene- and domain-specific calibration if deemed more appropriate^8,19^. Our work enables such an assessment in a systematic and rigorous way, providing a valuable resource for VCEPs to harmonize current gene- or disease-specific practices in variant effect predictor use and laying the foundation for targeted gene-specific predictor recommendations for clinical variant classification in the future.

Domain clustering greatly expands coverage to thousands of additional genes while still accounting for meaningful differences across domains and improving over the genome-wide approach. Our clustering approach relies solely on the distributional shape of all predictor scores within a protein domain, without supervision from gene or domain labels. Remarkably, certain domains cluster consistently across multiple predictors, suggesting that stereotyped predictor behavior is intrinsic to specific domain families. This pattern is observed in domains where predictors perform well (e.g., collagen) and domains where predictors struggle (e.g., fibronectin), offering opportunities for future mechanistic investigations into the determinants of predictor accuracy and iterative refinement of methods. Furthermore, general patterns across predictors are evident: right-shifted clusters for one predictor tend to correspond to right-shifted clusters for others, and similarly for left shifts, even for AlphaMissense which is not directly trained on ClinVar control variants. These concordant shifts further highlight opportunities for investigation of protein properties and molecular mechanisms that drive behavior across predictors.

Despite these advantages, our approaches have limitations. We focused only on a set of high-performing variant effect predictors that have already been calibrated for clinical use, and calibration of additional, well-performing predictors^37–41^ may provide improved coverage or evidence strength levels for specific genes. Similarly, there may be alternative calibration models to the ones considered here that are better at reducing misestimation errors in data-sparse genes and further development and optimization of such methods is an avenue for future work., Additionally, a limitation of the clustering approach as developed here is that it did not include non-domain regions due to nebulous local behavior near domain boundaries. While smaller clusters were prioritized to optimize calibration with limited labeled variants, larger clusters may be appropriate in some cases. Developing methods to incorporate non-domain variants and optimize clustering could further expand calibration coverage and accuracy. Furthermore, while we identified a trade-off between calibration quality (gene-specific calibration) and coverage of the clinical genome (domain-aggregate calibration), it is unclear how this trade-off can be eliminated. There may be other factors influencing the decision to choose between gene-specific and domain-aggregate calibration, e.g., the number of domains in a protein, the enrichment of control variants in certain domains within a protein relative to others or non-domain regions, the degree of distributional skew and heterogeneity across domains within the same protein, among others. Novel control variant aggregation strategies that account for these factors are further expected to improve both computational evidence coverage and strength. Finally, the gold-standard assessment of calibrations is not straightforward. While good calibration approaches reduce VUS fractions and improve TPR-FPR trade-offs, these metrics do not account for the fact that the true pathogenic and benign classes depend on evidence other than predictor scores. Since calibration does not change the ranking of original predictor scores, performance on these metrics is not solely dependent on the calibration approach and is influenced by a predictor’s accuracy itself. Nonetheless, our analyses allowed us to systematically evaluate how well gene-specific and domain-aggregate calibrations improve classification consistency with ClinVar controls, reduce indeterminate assignments, and balance sensitivity and specificity relative to genome-wide aggregation. Furthermore, our interval *LR* -based assessment better reflects the goal of calibration – deriving the correct amount of evidence for a predictor score.

Our findings also underscore the gene- and region-specific variability in predictor performance. Different variant effect predictors provide varying maximal strengths of evidence across genes, consistent with prior recommendations to select a single predictor before clinical use^8,19^. Our work suggests that optimal predictor choice may vary not only between genes but also across regions within a gene, as domain clustering can assign different evidence strengths to identical raw predictor scores depending on their location. This has direct practical implications for the design of clinical bioinformatics pipelines and the interpretation of computational evidence in variant classification. In summary, our work extends the applicability of variant effect predictors to genes and domains with limited labeled variants, improves evidence alignment with curated clinical classifications, and provides a foundation for more nuanced, region-specific use of computational predictors in clinical genomics.

## Methods

### Datasets and preprocessing

#### ClinVar Data

We utilized labeled variants from the January 2025 release (version 2025.01) of ClinVar in the main tab-delimited file available through the ClinVar FTP site (https://ftp.ncbi.nlm.nih.gov/pub/clinvar/tab_delimited/). This dataset includes variants annotated as pathogenic (P), likely pathogenic (LP), benign (B), or likely benign (LB). We restricted our analyses to missense variants with a review status of at least one star (i.e., with assertion criteria provided by at least one submitter or expert panel). Genes were required to have at least one ClinVar-annotated P or LP missense variant, thereby establishing missense variation as a disease-relevant mechanism for each gene. In addition, global allele frequency (AF) below 0.01 was used as a filtering criteria for these variants, prioritizing exome AF when available, and falling back to genome AF otherwise. Finally, to minimize confounding from splice-altering effects, variants with SpliceAI-predicted scores ≥ 0.2 were excluded; variants without a SpliceAI prediction were retained. Furthermore, to avoid bias, we excluded any variants known to be part of the training datasets of the prediction tools under evaluation (REVEL and MutPred2), when such information was available. The final number of qualifying variants per gene and predictor is summarized in **Supplementary Table 4**.

#### gnomAD Data

To estimate the prior probabilities of pathogenicity, we used variant data from gnomAD v4.1^42^, including both exome and genome datasets. VCF files were downloaded from the official gnomAD release site. Filtering steps matched those used on the ClinVar dataset. These gnomAD variants were also used to assess evidence assignment by different calibration approaches.

#### Prediction Score Mapping

The initial gene set for the dynamic clustering of protein domains was obtained from the GenCC database^15^ and filtered to retain those with clinical validity ≥ moderate (n=3,955). For gene-specific calibration, we began with genes represented in the January 2025 release of ClinVar. We then assembled and harmonized pre-computed variant-level predictions for these genes from three major variant effect predictors: REVEL^43^, MutPred2^24^, and AlphaMissense^44^. These predictors were selected to represent complementary methodological classes and training paradigms, including one meta-predictor (REVEL) and two primary predictors trained on distinct supervision sources. MutPred2 was trained using disease-associated variants, and AlphaMissense was developed using population variation and protein sequence–based features. Collectively, these three predictors were among the most accurate methods not incorporating allele frequency as an explicit feature in an independent benchmark^6^, and all have been previously subjected to post-hoc calibration analyses by Pejaver and Bergquist^8,9^. Harmonization proved challenging due to incomplete variant information and incompatible gene transcripts across predictors. MutPred2 annotations, which lacked genomic coordinates, and REVEL annotations, which lacked amino acid positions, were mapped to AlphaMissense through protein sequence and genomic coordinate alignment.

#### Protein Domain Mapping

Variants were mapped to protein domains obtained from Pfam, a curated database that catalogs these conserved protein regions across gene families. Overlapping domains were merged if they covered ≥50% of each other, otherwise the overlapping region was split evenly from the middle, ensuring each variant received a single, consistent domain annotation at the amino acid level.

### Dynamic clustering of protein domains

We grouped domains across genes based on similarity in their variant effect predictor score distributions, thereby pooling sparse control variant annotations across distributionally similar regions. For each predictor an all-by-all matrix of Jensen–Shannon distance (JSD)^45^ as a measure of distribution shape similarity was generated. Domains were then clustered hierarchically using the JSD distance matrix for each predictor. (**Extended Data Fig. 24**). To ensure adequate control variants for calibration across most clusters while preserving resolution of distribution shapes, we capped the size of clusters by the number of ClinVar labeled variants. We imposed a maximum number of ClinVar variants per cluster that yielded a median of 500 control variants. To achieve this, we dynamically partitioned the hierarchical clustering tree based on ClinVar variant count. Starting from the root of the dendrogram, the dynamic splitting algorithm iteratively traversed branches and split any branch containing more than a specified maximum number of ClinVar variants into two sub-branches, following the hierarchical structure. This process continued until all resulting clusters contained the threshold number of ClinVar variants or fewer. We performed a grid search across threshold values to identify the maximum ClinVar variant count that yielded a median cluster size closest to 500 variants. This ensures sufficient ClinVar variants for calibration for most clusters while maintaining granular grouping of distribution shapes. Each resulting cluster represents a set of protein domains drawn from multiple genes that share similar predictor score distributions. Variants falling outside annotated Pfam domains were not subjected to this clustering procedure and were simply aggregated into a separate non-domain “cluster”.

### Prior probability of pathogenicity estimation

The prior probability of pathogenicity (prior) plays a central role in determining the ACMG/AMP evidence strength under the Bayesian framework^7,17^ It enables the estimation of interpretable posterior probabilities and determines the likelihood ratio (*LR*) values corresponding to the different strengths of evidence. Until recently, a universal prior probability of 10% was applied based on the assumption that 1 out of 10 variants detected during clinical genetic testing turn out to be pathogenic. More recently, a genome-wide prior probability of 4.41% was estimated in a principled, data-driven manner as the proportion of pathogenic variants among all rare missense variants observed in disease-associated genes in gnomAD^8,9^, using a semi-supervised machine learning algorithm, DistCurve^16^. ClinVar P/LP variants and gnomAD variants, used as the set of labeled positives (*P*) and unlabeled data (*U*), were fed as input to the DistCurve algorithm to estimate the proportion of pathogenic variants in *U*. Though the estimated prior of 0.0441 is appropriate as a single universal prior probability of pathogenicity at a genome-wide scale, genes may harbor a much lower or higher proportion of pathogenic variants leading to incorrect posterior probabilities and *LR*-thresholds, and more importantly, inaccurate assignment of evidence strength. In order to improve on the limitations of a single genome-wide prior, we adopted a framework to derive more accurate prior probabilities for a variant depending on the gene or domain it belongs to by adapting DistCurve to estimate a prior probability unique to each gene with enough ClinVar labeled variants. For other variants from data-deficient genes, we first clustered domains to pool labeled variants and estimate the prior probabilities for each cluster using DistCurve.

#### DistCurve Algorithm

The DistCurve algorithm takes a labeled positive set of P/LP variants (*P*) and an unlabeled set of gnomAD variants (*U*) as input to estimate the proportion (prevalence rate) of positives in the unlabeled set (**Extended Data Fig. 1**). It relies on the variant scores from a predictor trained to separate the labeled positives from the unlabeled variants to compute pairwise distance between two variants as the absolute distance between their scores. Labeled positive variants from *P* are randomly picked with replacement iteratively; in each iteration the closest unlabeled variant is removed from *U* after recording the distance. DistCurve’s prior estimation approach relies on the fact that positives are preferentially removed under this approach and the recorded distances tend to be smaller until all unlabeled positives are removed after which the recorded distances tend to be larger. Precisely, the prior probability requires estimating the number of unlabeled positives, which is given by the iteration number after which the magnitude of recorded distances increases significantly. Visually it corresponds to the elbow of the distance curve with the recorded distances on the y-axis and the corresponding iteration on the x-axis. Computationally, the prior is inferred from a machine learning model pretrained on a diverse set of simulated data. The difference between the genome-wide prior, gene-specific prior and domain-clustering based prior estimates arises primarily from variants used to construct the labeled positives and the unlabeled sets.

Using scores from existing predictors for the distance computations in DistCurve is problematic due to overfitting, since some of the variants may have been included in their training sets. Filtering such variants is not feasible for most tools, whose training dataset might not be publicly available. Furthermore, such filtering would reduce the number of available P/LP variants, whose counts are typically small to begin with for a single gene. To overcome this limitation we trained a new gene/cluster-specific predictor using an out-of-bag (OOB) approach to separate the labeled positives from the unlabeled variants from the target gene or cluster. The OOB approach allows training and predicting on the same set of variants without overfitting by training an ensemble of predictors, each on a different bootstrapped random sample (random sampling with replacement) of the variants. Each variant received a score only from those members of the ensemble whose training bootstrap sample did not contain the variant. The final OOB score for a variant was an average of all the scores it received from each ensemble member. Since the raw features from REVEL and AlphaMissense were not available, we use the MutPred2 features^24^ for training our OOB predictor ensemble. Precomputed MutPred2 features are available for about 215 million missense variants across all human genes^46^.

#### Adapting DistCurve to this study

To further mitigate issues stemming from small numbers of P/LP variants, common variants, and size imbalance between ClinVar P/LP and gnomAD variant sets in score computation we applied the following key enhancements:

- **Unique Bootstrapping Sampling Approach**: Since ClinVar P/LP and gnomAD variant sets may contain the same variant, the standard bootstrapping technique used in the OOB approach, can lead to cases where a variant might appear in both the training and out-of-bag (test) sample, violating the core assumption that prevents overfitting. To address this issue we derived a *unique bootstrapping sampling approach* that ensures that any duplicate variant either appears in the training sample twice, as P/LP and also as gnomAD, or in the test sample twice but not in both.
- **Dimensionality Reduction via PCA**: The original MutPred2 feature space contains 1345 dimensions, posing overfitting risks due to the small variant set sizes. To mitigate this, we applied Principal Component Analysis (PCA) after z-score normalization to the training data in each bootstrap iteration, selecting the top *k* components explaining 95% of the variance. The same transformation (mean, scaling, projection) is then applied to the OOB variants before inputting them to the trained predictor. This retains the most informative components, improving model generalizability.
- **Random Under-Sampling (RUS)**: To address the substantial class imbalance between P/LP variants and gnomAD unlabeled variants, we applied RUS during each bootstrap iteration. The same number of variants were picked randomly with replacement from the ClinVar P/LP and gnomAD sets by undersampling from the larger set (here gnomAD). This balances the class distribution in the training set, reduces model bias toward the majority class, and enhances the sensitivity of pathogenicity prediction.

The OOB scores for the ClinVar P/LP and gnomAD variant sets were used by DistCurve for its distance computations, and ultimately, the estimation of priors. To account for the uncertainty in prior estimation on small datasets, we obtained 5000 estimates of the prior, each based on a different bootstrap sample. The bootstrapping for prior estimation was done separately from that for generating the OOB scores. The final prior probability was defined as the median of the 5000 bootstrap estimates, and uncertainty was summarized using the interquartile range (25th and 75th percentiles).

#### Gene/domain-specific prior estimation using adapted DistCurve

We estimated gene- and domain cluster-specific prior probabilities using our adapted DistCurve approach only for genes with at least 10 P/LP variants, resulting in prior estimates for 808 genes. To evaluate the reliability of the estimated priors, we computed the PU-AUC, defined as the area under the receiver operating characteristic curve for distinguishing ClinVar P/LP variants from unlabeled variants. We use PU-AUC as a proxy for the so-called *irreducibility assumption* made by DistCurve. Calibration methods were retained only if the PU-AUC exceeded 0.75, ensuring sufficient separability between positive and unlabeled variants. After applying this threshold, 648 of 808 genes were retained for this study (**Supplementary Table 1**). Similarly, cluster-level prior estimation retained 91 of 98 REVEL clusters, 88 of 92 AlphaMissense clusters, and 100 of 102 MutPred2 clusters (**Supplementary Table 5**).

### Calibration techniques considered

We evaluated ten post-hoc calibration methods spanning parametric, semi-parametric, and non-parametric approaches. These methods were selected to represent the principal functional forms used in probability calibration, including logistic transformations, distribution-based mixture algorithms, spline-based smoothers, and neural-network–based monotonic mappings. This diversity allowed us to assess calibration performance across a range of assumptions about calibration dataset requirements, score distributions and functional relationships between predictor scores and posterior probabilities.

#### Parametric calibration methods

Parametric methods assume a predefined functional form relating the raw prediction score to the calibrated posterior probability. Platt scaling^25^ models this relationship using a logistic function, providing a simple and widely used calibration approach. Weighted Platt scaling extends this framework by incorporating class weights to account for imbalance between pathogenic and benign variants. Beta calibration^26^ generalizes Platt scaling by introducing a three-parameter beta distribution-based transformation, enabling greater flexibility in modeling asymmetric score–probability relationships. We also evaluated distribution-based parametric mixture methods that explicitly model the score distributions of pathogenic and benign variants, separately. The skew-normal mixture model fits a two-component mixture of skew-normal distributions, allowing asymmetric score distributions^47–49^. The beta mixture model (different from the aforementioned beta calibration) uses mixtures of beta distributions to accommodate bounded and potentially bimodal score distributions. The truncated normal mixture model fits mixtures of truncated normal distributions to model bounded scores while capturing multimodal structure. Unlike other approaches where the posterior probability is directly estimated, in mixture-based approaches, it is estimated from the prior probability (α) and the fitted pathogenic and benign density functions as:

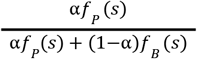

Where *f*_*P*_ (*s*) and *f*_*B*_ (*s*) are the fitted pathogenic and benign density functions for score *s*.

#### Non-parametric and semi-parametric calibration methods

Non-parametric and semi-parametric methods relax assumptions about functional form by relying on the calibration dataset itself, allowing more flexible mappings between scores and probabilities. Isotonic regression^50^ fits a monotonic stepwise function that preserves score ranking without imposing a parametric form. Smoothed isotonic regression^51^ refines this approach using piecewise cubic Hermite interpolating polynomials to produce smoother calibration curves while maintaining monotonicity. SplineCalib^52^ applies cubic smoothing splines to learn a smooth, non-linear mapping between scores and probabilities, providing greater flexibility than parametric approaches while retaining smoothness constraints. MonoPostNN, a monotonic neural-network–based calibration method developed for this study, learns a flexible non-parametric mapping from scores to posterior probabilities while enforcing monotonicity constraints. An open-source implementation is available at https://github.com/shajain/PosteriorCalibration.

Last, we compared these methods to the genome-wide calibration proposed in Pejaver et al.^8^ involving estimating local posterior probabilities for each predictor score. We refer to it here as the local posterior calibration method and systematically evaluated its performance through a sensitivity analysis on simulated data over its two main hyperparameters. Our goals were: (1) to quantify how different parameter settings affect calibration accuracy, and (2) to assess whether local calibration remains reliable under realistic, gene-level data constraints (e.g., small sample size and class imbalance). The two key parameters investigated were:

1. Window size: the proportion of labeled pathogenic and benign variants included in each sliding window used to estimate the local posterior. We tested window sizes of 10%, 20%, and 30% of the total labeled dataset.
2. gnomAD fraction: the minimum proportion of unlabeled gnomAD variants required within each window to stabilize the local posterior estimate. We evaluated values of 0%, 3%, and 6% (relative to the total gnomAD set).

All nine parameter combinations were evaluated using the ACMG-based misestimation metrics described below, with particular emphasis on *P*_*Fraction*_ and *B*_*Fraction*_, as metrics that reflect over-confidence in point estimation in clinically relevant pathogenic- and benign-supporting regions. This analysis enabled us to characterize the robustness of the local calibration method and to identify hyperparameter settings that minimize clinically meaningful miscalibration while maintaining overall accuracy.

### Large scale simulation experiments

To evaluate the robustness and accuracy of calibration methods under realistic conditions, we constructed synthetic variant effect predictor score sets that simulated real predictor score distributions for a gene or a domain cluster. Unlike the real scores, the true posterior probabilities of the simulated scores can be derived theoretically, which enables a robust evaluation of the estimated posterior probabilities by comparing against their true values. For each example distribution, we generated 924 simulated datasets, spanning combinations of 6 calibration set sample sizes, 11 observed proportion of pathogenic variants in each calibration set, and 14 true pathogenic prior probabilities, to systematically reflect a wide range of realistic data scenarios. This simulation framework was designed to determine whether alternative calibration approaches could outperform local calibration and to characterize the relative strengths and limitations of each method across varying data availability and prior misspecification conditions.

- **Generation of Synthetic Datasets**: We modeled variant effect predictor scores using four parametric distribution families commonly observed in real data: *Beta, truncated Normal, truncated Skew-t*, and *truncated Skew-Cauchy*. Together, these families capture a wide range of behaviors (bounded support, symmetry, skewness, and heavy tails) typical of real prediction score distributions. For each real score set across genes and predictors, pathogenic and benign scores were independently fitted to the four candidate families. Each unique pair of pathogenic and benign fitted distribution was considered. Random samples from each distribution pair were then used as synthetic pathogenic and benign score sets, providing realistic yet controlled scenarios for benchmarking calibration methods.
- **Calibration Set**: For each fitted pathogenic/benign score distribution, we generated calibration sets of varying sizes to assess sample efficiency:

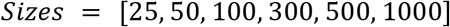

To examine robustness to class imbalance, we varied the observed proportion of pathogenic variants in the calibration set across:

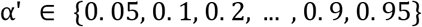
- **Test Set and True Posterior**: For each scenario, we generated an independent test set of 1000 variants. The true pathogenic prior probability α was varied to reflect different gene-specific priors:

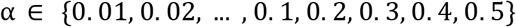

The true posterior probability for the test set score *s* was computed from the prior probability (α) and the fitted pathogenic and benign densities (*f*_*P*_ (*s*) and *f*_*B*_ (*s*), respectively) as

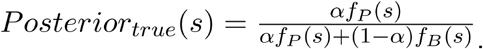
- **Prior Adjustment**: Most post-hoc calibration methods (e.g., Beta, Platt, SplineCalib, Isotonic, SmoothIsotonic, BetaMixture, TruncNorm) implicitly assume that the class prior is the same in the calibration and test sets. To account for prior shifts (α′ ≠ α), we applied a post-hoc correction to the calibrated posteriors at test time. Let ρ(*s*) denote the uncorrected calibrated posterior from the calibration set and *α*’ the pathogenic fraction in the calibration set. We first compute the likelihood ratio, accounting for the incorrect prior, as

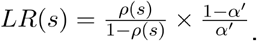

Next we obtain the prior-corrected posterior as

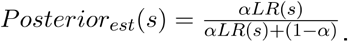
- **ACMG Points Misestimation Error:** To assess calibration quality in clinically meaningful terms, we quantified the discrepancy between calibrated and true posterior probabilities using a novel metric directly tied to ACMG/AMP evidence strength. Specifically, instead of measuring the difference between the estimated and true posterior probabilities, we measure the difference between the estimated and true ACMG evidence points Δ*EP*(*s*) at score *s*. This metric reflects the number of evidence points that are misestimated due to calibration error, providing a clinically relevant measure of miscalibration severity. The number of ACMG evidence points at score *s* with a likelihood ratio *LR*(*s*) is given by

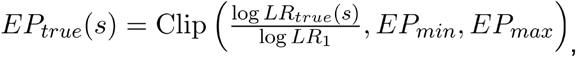

where *LR*_1_ is the likelihood ratio threshold corresponding to +1 ACMG evidence point (Supporting-level pathogenic evidence); the *Clip* function truncates the evidence points between their minimum and maximum possible values of *EP*_*min*_ = −8 and *EP*_*max*_ = +8, respectively. Note that this formula gives ACMG evidence points on a continuous scale and can be discretized to obtain the integer valued points. We clip the points between -8 and +8, instead of the -4 and +4, the maximum allowable strength level for computational evidence. This choice enables capturing errors due to overconfident point assignments effectively; for example, when the formula gives +5 as true evidence and +7 as estimated evidence the resulting error is Δ*EP*(*s*) = +2 whereas clipping with -4 and +4 would result in zero error.

The true ACMG evidence points *EP*_*true*_(*s*) is obtained by plugging in the true likelihood ratio *LR*_*true*_(*s*) obtained as the odds ratio of the true posterior probability *Posterior*_*true*_(*s*) and the prior probability α (**Fig. 1a**). Similarly the estimated ACMG evidence points *EP*_*est*_(*s*) is obtained by plugging in the estimated likelihood ratio *LR*_*est*_(*s*) obtained as the odds ratio of the estimated/calibrated posterior probability *Posterior*_*est*_ (*s*) and the prior probability α (**Fig. 1a)**. Our Calibration metrics are expressed in terms of the difference between the estimated and true ACMG evidence points:

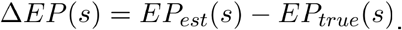

When calibration at score *s* is accurate, Δ*EP*(*s*) is close to zero. A positive value of Δ*EP*(*s*) for a score *s* in the Pathogenic region (where the true ACMG points for *s, EP*_*true*_(*s*) is one or more) indicates overconfidence in assigning pathogenic evidence. Similarly a negative value of Δ*EP*(*s*) for a score *s* in the Benign region (where the true ACMG points for *s, EP*_*true*_(*s*) is -1 or less) indicates overconfidence in assigning benign evidence.

- **Calibration Metrics and Ranking Criteria**: We consider the following three metrics that summarizes the misestimation of points over the entire test score set and use them to compare and rank methods.
  1. **Mean absolute points misestimation error (MAPE)**: The average of |Δ*EP*(*s*)| over all scores in the test set.
  2. **Proportion of over-confident pathogenic assignments** (*P*_*Fraction*_): The proportion of scores assigned at least one additional ACMG point above the true value, i.e., Δ*EP*(*s*) > 1 among all test scores lying in the Pathogenic region, where *EP*_*true*_ (*s*) > 1.
  3. **Proportion of over-confident benign assignments** (*B*_*Fraction*_): The proportion of scores assigned at least one additional benign (negative) ACMG point below the true value, i.e., Δ*EP*(*s*) < −1 among all test scores having *EP*_*true*_ (*s*) < −1.

These three metrics emphasize clinically meaningful miscalibration on the ACMG point scale and explicitly consider the overconfidence in point assignments, facilitating selection of conservative calibration approaches.

- **Unified Evaluation Protocol**: All methods were compared under a common evaluation framework. For each simulated data scenario, we generated 30 independent datasets and applied every calibration method to quantify variability and robustness. The best-performing configuration of the local calibration method—defined by its optimal combination of window size and gnomAD fraction—was included in all comparisons. The evaluation was designed to identify whether any alternative method consistently outperforms local calibration, particularly under realistic gene-level constraints such as limited labeled data and strong class imbalance. A broader goal was to inform the design of dynamic workflows that could leverage the strengths of multiple methods.
- **Analysis of Failure Cases**: Finally, we identified and examined scenarios in which individual calibration methods performed poorly, with particular attention to extreme class imbalance and severe data sparsity. These failure cases were used to diagnose method-specific limitations and to develop practical recommendations for method choice under constrained conditions.

### Data-adaptive calibration framework for genes and domain clusters

Genes eligible for gene-specific calibration were identified using ClinVar (January 2025 release) based on the number and class distribution of pathogenic/likely pathogenic (*N*_*P*/*LP*_) and benign/likely benign (*N*_*B*/*LB*_) single-nucleotide variants. For each gene, we defined ***sum*** as the total number of labeled variants (*N*_*P*/*LP*_ + *N*_*B*/*LB*_) and ***pfrac*** as the fraction of pathogenic variants, calculated as *N*_*P*/*LP*_/(*N*_*P*/*LP*_ + *N*_*B*/*LB*_). Eligibility criteria were designed based on observations from our large-scale simulation experiments, balancing statistical stability with gene coverage by imposing tiered requirements on sample size and class distribution.

Genes with 50–99 labeled variants were considered suitable only if the pathogenic fraction was moderately balanced (0.4 ≤ *pfrac* ≤ 0.7), ensuring adequate representation of both P/LP and B/LB variants under limited sample sizes (condition 1). For genes with 100–299 labeled variants, broader class imbalance was permitted (0.1 ≤ *pfrac* ≤ 0.9), reflecting improved estimation stability with increased sample size (condition 2). Genes with ≥300 labeled variants were eligible under more relaxed class ratio constraints (0.05 ≤ *pfrac* ≤ 0.99), provided that at least 10 B/LB variants were present to enable stable estimation of score distributions in the benign range (condition 3).

Across all filtered genes, 157 met at least one of the predefined eligibility criteria. Of these, 36 genes satisfied the criteria for condition 1, 94 genes met condition 2, and 27 genes met condition 3. Each eligible gene then underwent gene-specific calibration, implemented through a workflow that adaptively selects the optimal calibration methods for each gene using simulated score sets, preserving distribution characteristics and data constraints of the gene. Our strategy is divided into four steps.

1. **Gene-specific prior estimation**: For each gene, we first estimated the prior probability of pathogenicity using an adapted *DistCurve* algorithm^16,53^ leveraging ClinVar control pathogenic variants together with unlabelled variants from gnomAD. This procedure yields a gene-specific prior probability of pathogenicity that was subsequently used to generate simulated datasets and to define the posterior probability ranges corresponding to ACMG/AMP evidence points.
2. **Simulated score set generation**: For each gene, we independently fitted the pathogenic and benign variant score sets using four parametric distribution families: *Beta, Truncated Normal, Truncated Skew-t*, and *Truncated Skew-Cauchy*. The best-fitting distribution was picked using the maximum log-likelihood criteria. Randomly sampling from the fitted distributions, we generated 30 synthetic score sets per gene. The number of pathogenic and benign scores in each simulation were kept identical to that from the gene. The true posterior probabilities for the simulated set were computed from the estimated gene prior and fitted distribution densities as described above in the ‘Large-scale simulation experiments’ section.
3. **Application of calibration methods:** For each gene, we applied 9 local calibration configurations (varying window size and gnomAD smoothing fraction) as in Pejaver et al.^8^ and 10 alternative post hoc calibration methods^53^ on each synthetic score set. Using 1,000 bootstrap samples, an out-of-bag approach was applied to obtain calibrated posterior probability estimates. In one bootstrap iteration, the calibration method was trained on the bootstrapped score sample and applied to out-of-bag scores (those not in the bootstrap sample). Across all bootstrap iterations, each score in the score set was out-of-bag several times and, thus, received many posterior probability estimates. Following Pejaver et al.^8^, the 5th and 95th percentile of all posterior probability estimates a score receives were used as conservative estimates in the pathogenic- and benign-supporting regions described in the ‘Large-scale simulation experiments’ section, respectively. Using the conservative posterior probability estimates mitigates the risk of overestimating evidence strength both in the pathogenic and benign regions. The scores in the indeterminate region were assigned posterior probabilities from the calibration method trained on the entire synthetic score set. In this manner, posterior probability estimates for each of the 30 synthetic score sets were obtained.
4. **Multi-stage method selection**: Using the simulated score sets, a single calibration method was selected for each gene through a three-stage filtering procedure:
  a. An over-confident estimation risk filter was applied to remove calibration methods exhibiting clinically unsafe behavior. This was defined as more than 25% of the 30 synthetic score sets having the 75th percentile or the maximum ACMG point estimation error greater than 2 or 3 evidence points, respectively, in the pathogenic- or benign-supporting regions. This corresponded to using Δ*EP*(*s*) and -Δ*EP*(*s*) as the errors in the pathogenic and benign regions, respectively.
  b. The top three methods with the lowest Mean Absolute Point Estimation (MAPE) error across the entire simulated score set were retained. The ranks were averaged over the 30 synthetic sets to get the final ranking.
  c. Out of the top 3 methods, the one with the lowest combined proportion of over-confident point assignment in pathogenic- and benign-supporting regions *P*_*Fraction*_+ *B*_*Fraction*_ is picked as the optimal calibration method for the gene. The ranks were averaged over the 30 synthetic sets to get the final ranking.
5. **Computing the optimal posterior probability for the gene:** The selected optimal calibration method was then applied to the target gene score set to produce final calibrated posterior probabilities. The same bootstrap and out-of-bag approach, as used for the synthetic data, was applied. The conservative 5th and 95th percentile out-of-bag posterior probability estimates were used to derive the pathogenic and benign evidence score thresholds.

#### Domain-aggregate calibration

This framework can also be applied to domain-aggregate calibration. In this context, we first estimated cluster-specific prior probabilities. We then simulated a score set for each cluster and applied the same multi-stage method selection process to the simulated data in order to identify the optimal calibration method. The selected method was subsequently used to compute the final posterior probabilities for the target cluster.

### Calibration performance evaluation

To assess the practical utility of our data-adaptive calibration framework, we conducted a series of real-world validation experiments using data from ClinVar, the Clinical Genome Resource (ClinGen), and ClinGen Variant Curation Expert Panel (VCEP) annotations.

#### Interval-based likelihood ratio evaluation (evidence strength accuracy)

To evaluate calibration performance at the level of ACMG/AMP evidence strengths, we applied the Bayesian likelihood-ratio framework proposed by Tavtigian *et al*.^*17*^, which defines theoretical *LR* thresholds for each evidence level (8 total): ±1 point (supporting), ±2 points (moderate), ±3 points, ±4 points (strong), given a prior probability of pathogenicity (**Fig. 1f**). These *LR* thresholds are used to determine the number of points a given piece of evidence qualifies for. Let *LR*_*x*_ denote the *LR* threshold for *x* points. If the *LR* of a variant based on some evidence (e.g., predictor score) is between thresholds *LR*_2_ and *LR*_3_ than it qualifies for +2 points towards pathogenicity. Similarly, if the *LR* is between *LR*_−2_ and *LR*_−3_ it qualifies for -2 points towards benignity.

Unlike the *LR* corresponding to an individual variant’s predictor score, which is estimated through a calibration model, the interval *LR*, summarizing the *LR* of a group of variants lying in a given score interval, can be calculated without a model, as the ratio of the proportion of pathogenic variants within the interval among all pathogenic variants to the proportion of benign variants within the interval among all benign variants. Consequently it serves as an unbiased measure of *LR*, for a fair evaluation of the score intervals and points assigned by a calibration method by providing theoretical *LR* thresholds to compare against. The interval *LR* is expected to be close to the corresponding *LR* threshold. For example, the interval *LR* for the score interval corresponding to +2 and +3 points is expected to be close to *LR*_2_. However, in order to reflect our preference towards a conservative point assignment, we consider interval *LR* value above (below) the *LR* threshold as preferable over a value below (above) the threshold for pathogenicity (benignity) points. An interval *LR* below (above) the threshold implies that the variants in the score interval have less evidence towards pathogenicity (benignity) relative to the points assigned by the calibration method.

For each gene (gene-specific calibration) or each cluster (domain-aggregate calibration), we estimated the prior probability of pathogenicity from ClinVar P/LP variants and gnomAD variants, and used this prior to compute the *LR* threshold for each evidence category using the Tavtigian framework^7^.

We then calculated interval *LRs* for the calibrated score intervals generated by the two calibration approaches evaluated in this study: (1) gene-specific or domain-aggregate calibration (our methods), and (2) the previously published genome-wide aggregation^8,9^.

For each evidence level, we determined which method “won” using the following rules: (1) If both methods had an interval LR either above (pathogenic) or below (benign) the *LR* threshold, the method whose *LR* was closest to the threshold was counted as the winner. (2) If the two methods had interval *LRs* on the opposite side of the threshold, the method with higher (lower) interval LR was considered as the winner for pathogenic (benign) evidence, giving higher preference to a conservative method. For each gene or cluster, we counted the number of evidence levels (out of eight possible) in which a method won. To assess whether gene-specific and domain-aggregate calibration systematically outperformed the genome-wide aggregation approach across genes, we applied a one-sided binomial test. The null hypothesis assumed that both methods were equally likely to “win” for any given gene or cluster (p = 0.5), where the *K* parameter (number of successes) was the number of genes or clusters where gene-specific or domain-aggregate calibration outperformed the genome-wide aggregation method, and the *N* parameter (number of trials) was the total number of evaluable genes, excluding genes or clusters for which the two methods were tied.

#### Classification consistency with ClinVar (outcome-level validation)

Using the final calibrated thresholds selected by the data-adaptive calibration framework, we compared variant classifications derived from gene-specific calibration or domain-aggregate calibration against those produced using published genome-wide aggregated thresholds^8,9^. Three complementary analyses were performed:

1. **Evidence assignment distribution (heatmap):** For each gene (gene-specific calibration) or cluster (domain-aggregate calibration) and for each predictor, we quantified the percentage of variants assigned to each evidence point category: ±1 (supporting), ±2 (moderate), ±3, and ±4 (strong). This was done separately for: ClinVar P/LP, B/LB, and VUS variants, gnomAD variants, and all possible missense SNVs. This was applied to three types of pairwise comparisons:
  - Gene-specific calibration vs genome-wide aggregation (**Fig. 3c**)
  - Domain-aggregate calibration vs genome-wide aggregation (**Fig. 5c**)
  - Gene-specific calibration vs domain-aggregate calibration (**Fig. 6b**) These heatmaps allowed us to assess whether gene-specific or domain-aggregate calibration: (1) reduced the fraction of variants remaining in the indeterminate region (point = 0), and (2) prevented misassignment to overly strong or inappropriate evidence levels.
2. **TPR–FPR difference (scatter plot):** For ClinVar P/LP and B/LB control variants, we evaluated difference in classification performance between two calibration approaches by calculating the true positive rate (TPR) and false positive rate (FPR), separately for the pathogenic and benign classes When treating pathogenic variants (P/LP) as positive, TPR was calculated as the proportion of P/LP variants assigned pathogenic evidence (≥ +1 evidence point), and FPR as the proportion of B/LB variants incorrectly assigned pathogenic evidence. Conversely, when treating benign variants (B/LB) as positive, TPR and FPR were computed using ≤ −1 evidence point. For each method, the average TPR and FPR were obtained by averaging the pathogenic and benign sides:

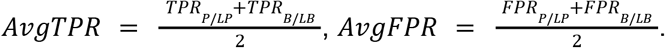

The Δ*AvgTPR* and Δ*AvgFPR* were then computed as the difference between the two calibration approaches being compared:

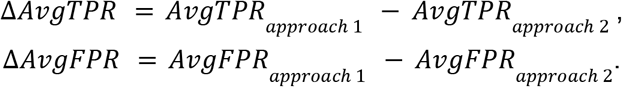

This was applied to three types of pairwise comparisons:
  - Gene-specific calibration vs genome-wide aggregation (**Fig. 3e**)
  - Domain-aggregate calibration vs genome-wide aggregation (**Fig. 5e**)
  - Gene-specific calibration vs domain-aggregate calibration (**Fig. 6a**) Positive Δ*AvgTPR* and/or negative Δ*AvgFPR* indicates improved performance of the first method relative to the second, and results were visualized as scatter plots for each comparison.
3. **Summarized metrics (metric barplots):** We computed several summary metrics for each calibration method (gene-specific/domain-aggregate/genome-wide calibration):
  - **Determinate rate (SNV)**: proportion of all possible SNVs assigned a nonzero evidence point by gene-specific, domain-aggregate, or genome-wide calibration.
  - **Average TPR–FPR difference**: TPR and FPR were calculated separately for pathogenic and benign sides, averaged within each side, and then subtracted (AvgTPR − AvgFPR). This is equivalent to the average Youden’s J statistic, a well-established metric in diagnostic testing^54^.
  - **Matthews correlation coefficient (MCC)**: computed separately for the pathogenic (*MCC*_*P*/*LP*_) and benign (*MCC*_*B*/*LB*_) sides, and then took the mean as the final *MCC*.

#### ClinGen annotation–based reclassification analysis

For genes with curated variant classifications from ClinGen Variant Curation Expert Panels (VCEPs), we evaluated how gene-specific and domain-aggregate calibration would alter computational evidence assignment and downstream ACMG/AMP clinical interpretation.

Under the ACMG/AMP framework as implemented by ClinGen, the computational evidence codes PP3 (pathogenic-supporting) and BP4 (benign-supporting) contribute additive evidence points toward the final classification. When no strength modifier is specified, PP3 and BP4 correspond to ±1 evidence point (+1 for PP3, −1 for BP4). When strength modifiers are applied, Supporting corresponds to ±1 point, Moderate to ±2 points, and Strong to ±4 points. Thus, recalibration of computational thresholds directly changes the numerical contribution of PP3/BP4 to the total evidence score and can shift the final pathogenicity classification.

Variant-level prediction scores from REVEL, AlphaMissense, and MutPred2 were retrieved for all curated variants. Each variant was reclassified under two calibration schemes: (1) Published genome-wide aggregation calibration thresholds^8,9^, and (2) Gene-specific thresholds selected by our data-adaptive calibration framework.

For each calibration scheme, we reassigned the computational evidence-defined evidence points while keeping all other ClinGen evidence codes (“Met Codes”) unchanged. We then recomputed the overall ACMG/AMP classification by summing evidence points according to ClinGen’s point-based system.

We quantified the extent and direction of reclassification relative to the original ClinGen classification. Transitions (e.g., VUS to B/LB or VUS to P/LP) were summarized and visualized using Sankey plots to illustrate shifts in clinical interpretation attributable to recalibrated computational evidence.

To isolate the contribution of computational evidence independent of prior ClinGen curation decisions, we additionally removed the original PP3/BP4 code from the ClinGen evidence list and recalculated clinical significance using only non-computational evidence. This produced a computational-null baseline classification. We then compared this baseline to classifications defined only by: (1) ClinGen-provided PP3/BP4 codes, (2) genome-wide aggregation calibration thresholds, and (3) gene-specific calibration thresholds.

For each method, we evaluated per-gene classification accuracy among variants assigned to non-indeterminate evidence levels and quantified the number of variants for which PP3/BP4 evidence could be assigned. This framework allowed us to determine whether gene-specific calibration is consistent with the non-computational evidence defined classification, increases the number of variants that are not in the indeterminate region, and reduces incorrect computational evidence relative to the aggregate calibration approach.

#### Biobank validation of variant classes

To further validate our calibration results, we assessed the relationship between computationally predicted variant classes and clinical phenotypes in ∼400,000 participants in the All of Us biobank. The validation was performed within the All of Us Workbench using the All of Us Controlled Tier Dataset v8 ^21,55^.

We used the case-control cohorts established in Tejura et al. (2026)^31^ to validate calibrations for 17 gene-disease pairs (all of these genes have score thresholds from genome-wide and domain-aggregate calibration and 5 of these genes have score thresholds from gene-specific calibration for all three variant effect predictors). Case cohorts were defined by including participants with phenotype-specific conditions, measurements, or survey responses; control cohorts excluded participants with the phenotype or closely related traits (**Supplementary Table 10**). For each predictor, carriers were defined as participants with one or more variants within a range of calibrated evidence points. Participants with variants in the gene with SpliceAI scores >0.2 were excluded.

For each variant class (range of evidence points), odds ratios of disease phenotype, given carrier status, were estimated using logistic regression. The model included carrier status, sex assigned at birth (male, female, other), age at data release, and 16 genetic ancestry principal components. We report point estimates, 95% confidence intervals, and Wald test p-values for the carrier status coefficient.

## Data Availability

All supplementary data files and large supporting files for analysis and figure generation are hosted at https://zenodo.org/uploads/18668684.

All calibration visualizations are available at https://igvf.mavedb.org/.

## Code availability

Data-adaptive calibration framework is available at the following repository: https://github.com/yileevechen/VEP_calibration

## Acknowledgements

This work was primarily supported by the following National Institute of Health (NIH) National Human Genome Research Institute (NHGRI) Impact of Genomic Variation on Function (IGVF) consortium center awards: the Center for Actionable Variant Analysis (CAVA) to D.M.F. and L.M.S. (UM1HG011969), the Radivojac Center to P.R. (U01HG012022), and the Craven Center to M.C. (U01HG012039). Additional financial support was provided by an NHGRI Advancing Genome Medicine Research consortium award (R01HG013025, D.M.F. and L.M.S.), by NHGRI grant (R01HG013350, H.S., T.B., and V.P.), the generous support of the Brotman and Baty families through the Brotman Baty Institute for Precision Medicine, and a Medical Genetics Postdoctoral Fellowship (T32GM007454, S.F.). We gratefully acknowledge All of Us participants for their contributions and thank the NIH All of Us Research Program for making available the participant data. This work was also supported in part through the Minerva computational and data resources and staff expertise provided by Scientific Computing and Data at the Icahn School of Medicine at Mount Sinai and supported by the Clinical and Translational Science Awards (CTSA) grant UL1TR004419 from the National Center for Advancing Translational Sciences. Research reported in this publication was also supported by the Office of Research Infrastructure of the NIH under award number S10OD030463 and S10OD038231. This research was also supported [in part] by the Intramural Research Program of the NIH. The contributions of the NIH author(s) are considered Works of the United States Government. The findings and conclusions presented in this paper are those of the author(s) and do not necessarily reflect the views of the NIH or the U.S. Department of Health and Human Services.

## Author Contributions

This project was conceptualized by Y.C., S.F., S.D.M., P.R., L.M.S., D.M.F., and V.P. Methodology was developed by Y.C., S.F., S.J., M.B., Y.S., and V.P. Software was developed by Y.C., S.F., J.S., H.S., and M.B. Validation was performed by Y.C., S.F., S.J., M.B., and Y.S., with supervision from M.W.C. and V.P. Formal analysis was conducted by Y.C., S.F., S.J., M.B., and Y.S. Investigation was carried out by Y.C., S.F., S.J., M.B., and J.S. Data curation and resource preparation were performed by Y.C., S.F., M.B., Y.S., and T.B. Computational support for calibration simulations was provided by H.S. Variant score mapping and MutPred2 computations were performed by T.B. Visualization was performed by Y.C., S.F., and R.S. The original draft of the manuscript was prepared by Y.C. Writing review and editing were performed by Y.C., S.F., S.J., M.B., P.R., L.M.S., D.M.F., and V.P., with input from all authors. Predictor calibration was led by S.F. and Y.C., with contributions from M.B., S.J., and H.S. Development of computational platforms, including MaveMD and the CVFG website, was led by J.S. Supervision was provided by S.D.M., M.W.C., P.R., L.M.S., D.M.F., and V.P. Project administration was carried out by J.S., L.M.S., and D.M.F. Funding acquisition was led by S.D.M., M.W.C., P.R., L.M.S., D.M.F., and V.P. All authors reviewed and approved the final manuscript.

**Extended Data Fig. 1:**
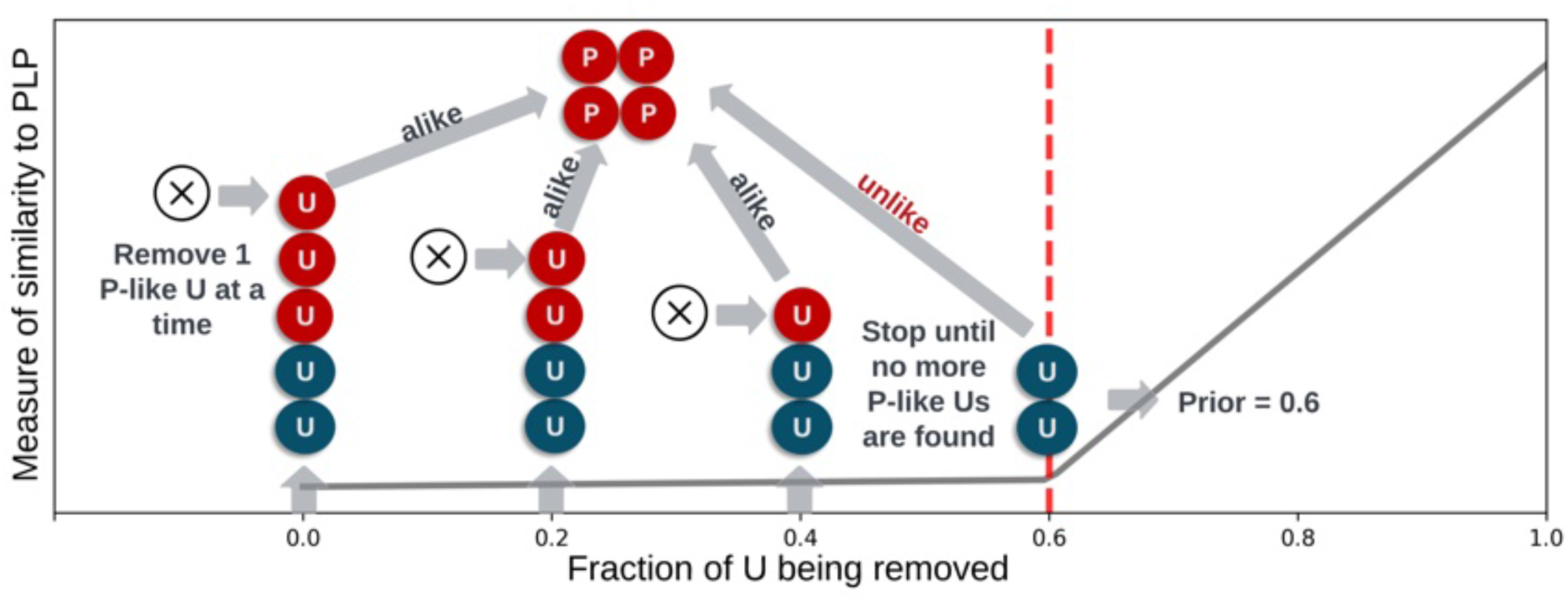
DistCurve algorithm schematic. Schematic of the DistCurve algorithm for estimating gene-specific pathogenicity priors. The algorithm iteratively identifies ClinVar pathogenic-like (P-like) variants from gnomAD unlabeled variants (U), removing them at each step. Distance between remaining P (ClinVar pathogenic, red circles) and U (gnomAD, blue/red circles) sets is computed iteratively. Red U circles represent P-like samples identified during the process. The algorithm terminates when no more P-like samples can be identified, indicated by the vertical dashed red line, corresponding to the turning point in the distance curve which estimates the prior.

**Extended Data Fig. 2:**
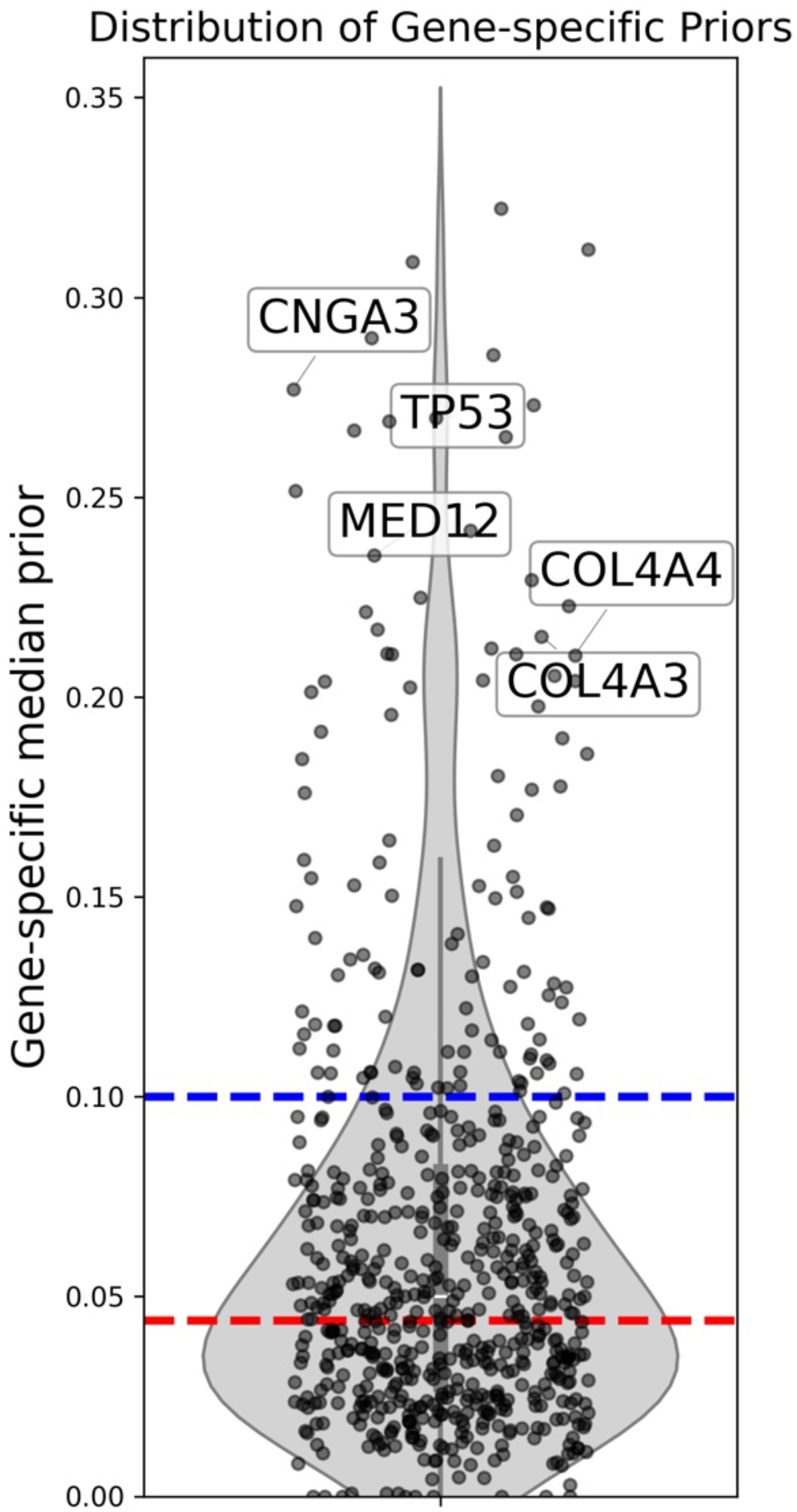
DistCurve-estimated gene-specific pathogenicity priors. Distribution of gene-specific prior probabilities of pathogenicity estimated using the DistCurve algorithm across analyzed genes. Each point represents the estimated prior for a single gene, demonstrating substantial heterogeneity relative to commonly used fixed genome-wide priors. The red dashed line indicates the 4.41% prior reported by Pejaver *et al*., whereas the blue dashed line indicates the 10% prior proposed by Tavtigian *et al*. The five genes with the highest estimated priors are labeled.

**Extended Data Fig. 3:**
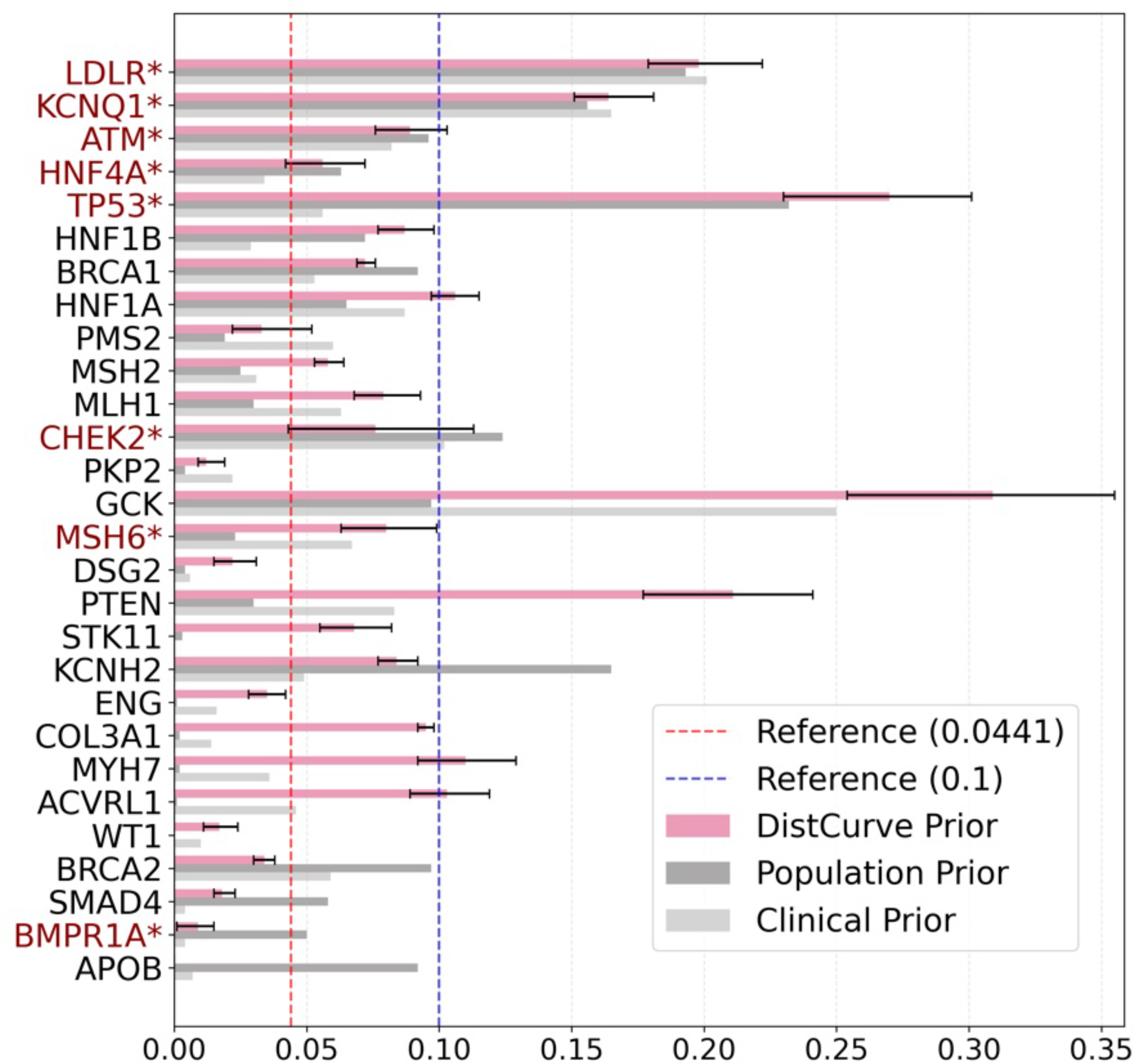
Comparison of gene-specific prior probabilities across data sources. Comparison of DistCurve-derived gene-specific prior probabilities of pathogenicity (pink) with population-based priors derived from UK Biobank (Population prior; dark gray) and clinically derived priors from ClinVar (Clinical prior; light gray). Each set of bars represents one gene. Asterisks indicate genes for which the DistCurve estimate falls within the interquartile range (Q1–Q3) of either the population or clinical prior distribution. Horizontal dashed lines denote previously reported reference priors: 4.41% (red dashed line; Pejaver *et al*.) and 10% (blue dashed line; Tavtigian *et al*.).

**Extended Data Fig. 4:**
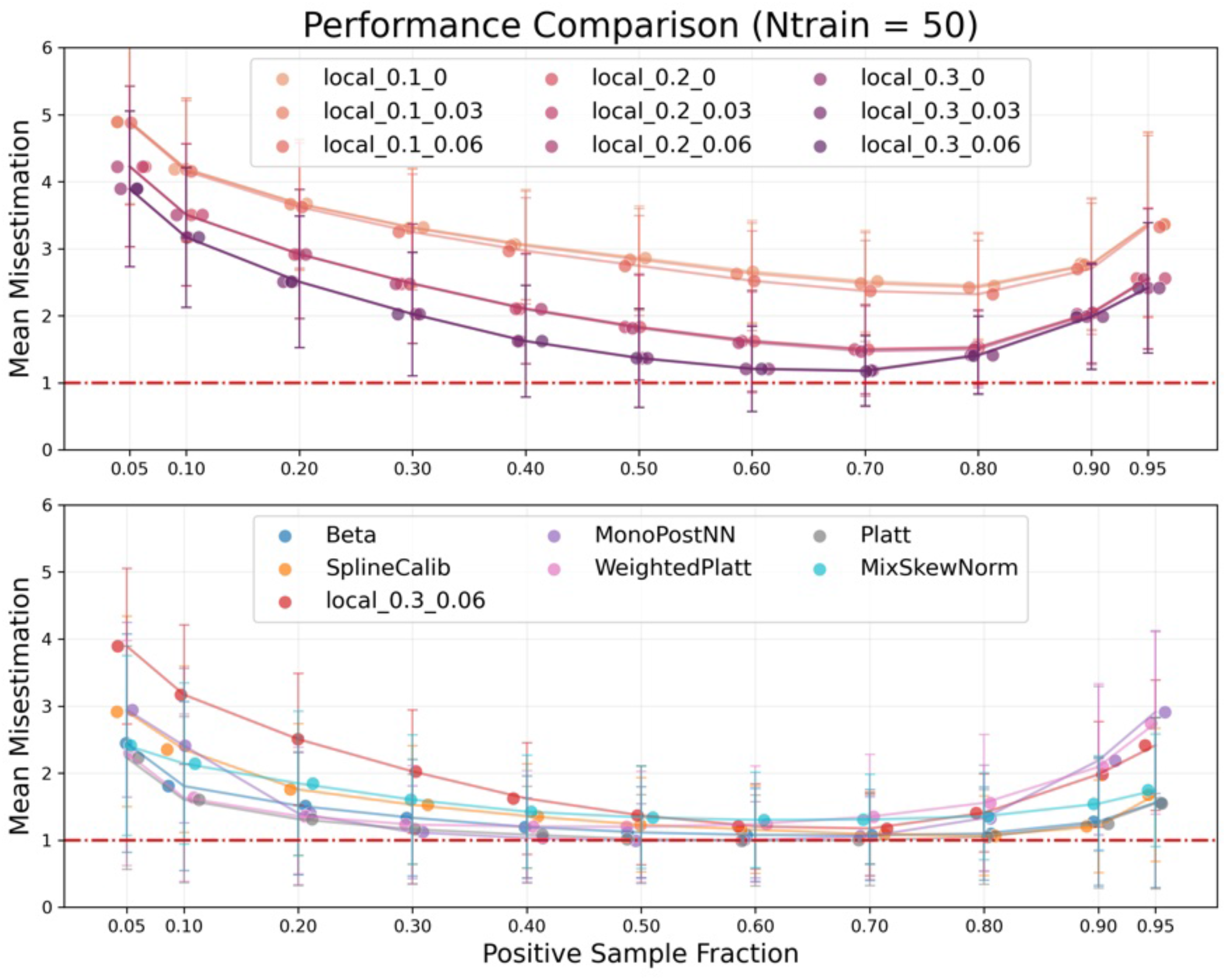
Calibration performance comparison with 50 variants. Performance comparison of calibration methods with calibration set size = 50 variants. Results are evaluated on test sets of 1,000 variants under varying class balance conditions. Error bars denote standard deviations across simulated true prior probabilities. Only methods achieving average misestimation below 1 in at least one scenario are shown.

**Extended Data Fig. 5:**
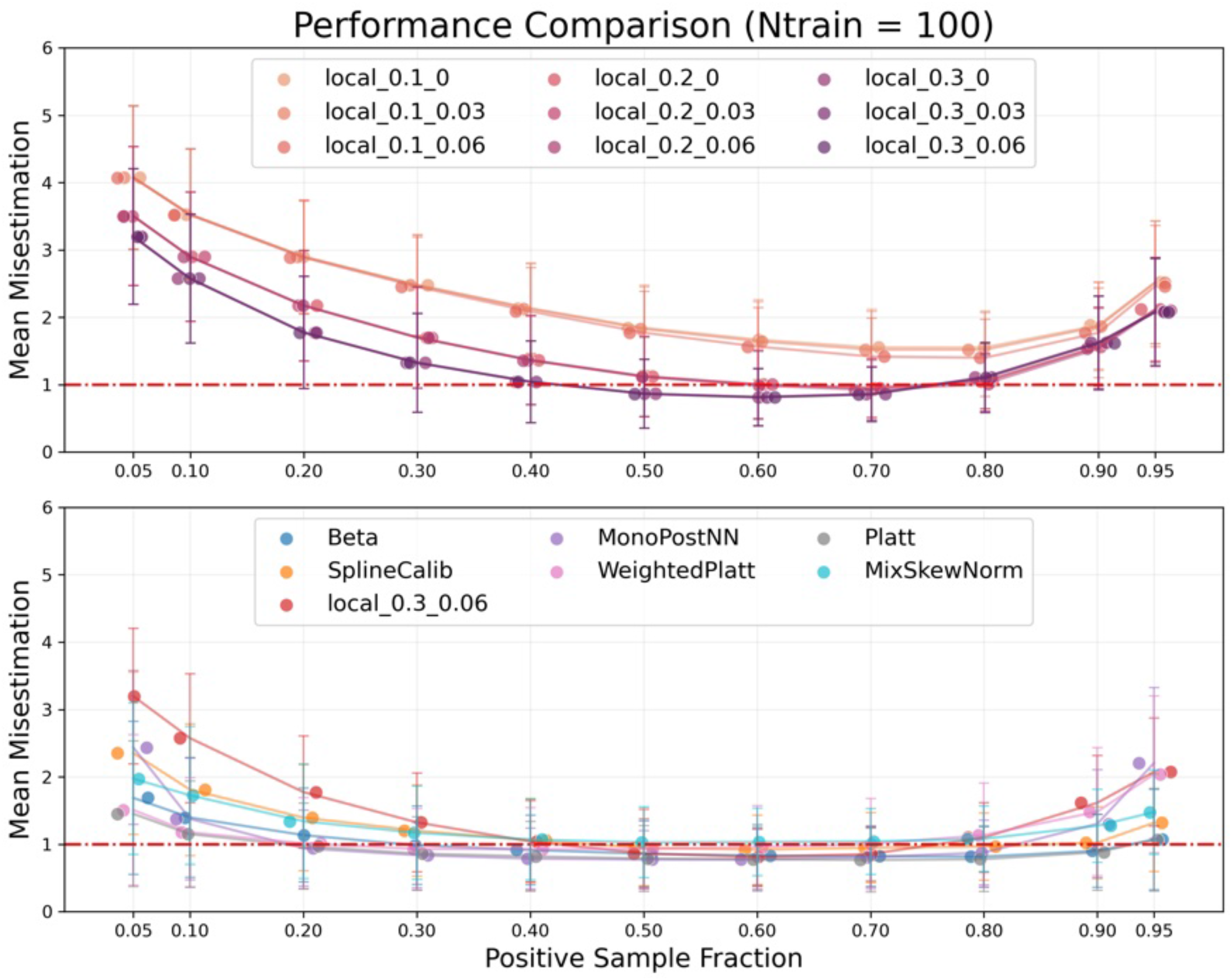
Calibration performance comparison with 100 variants. Performance comparison of calibration methods with calibration set size = 100 variants. Results are evaluated on test sets of 1,000 variants under varying class balance conditions. Error bars denote standard deviations across simulated true prior probabilities. Only methods achieving average misestimation below 1 in at least one scenario are shown.

**Extended Data Fig. 6:**
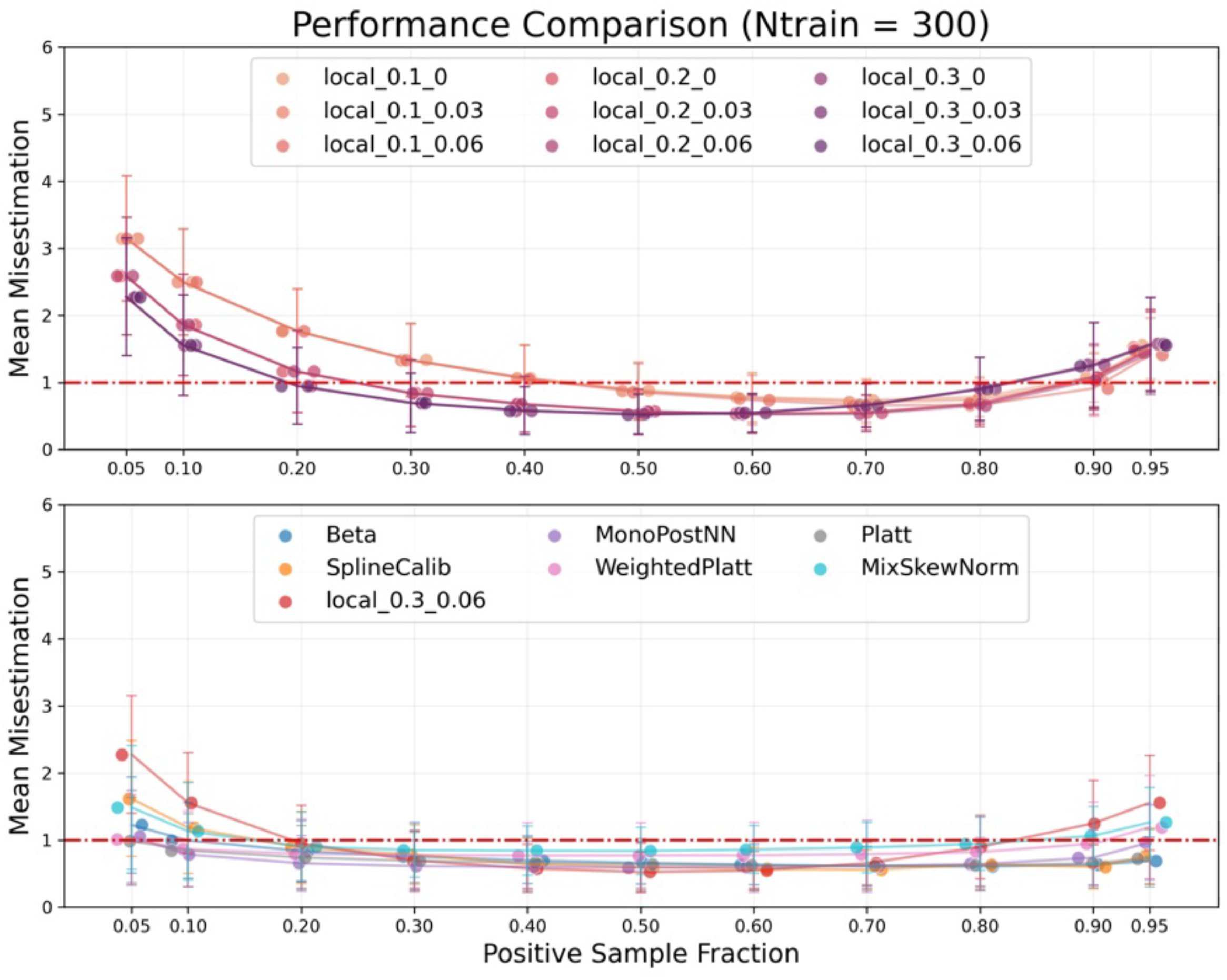
Calibration performance comparison with 300 variants. Performance comparison of calibration methods with calibration set size = 300 variants. Results are evaluated on test sets of 1,000 variants under varying class balance conditions. Error bars denote standard deviations across simulated true prior probabilities. Only methods achieving average misestimation below 1 in at least one scenario are shown.

**Extended Data Fig. 7:**
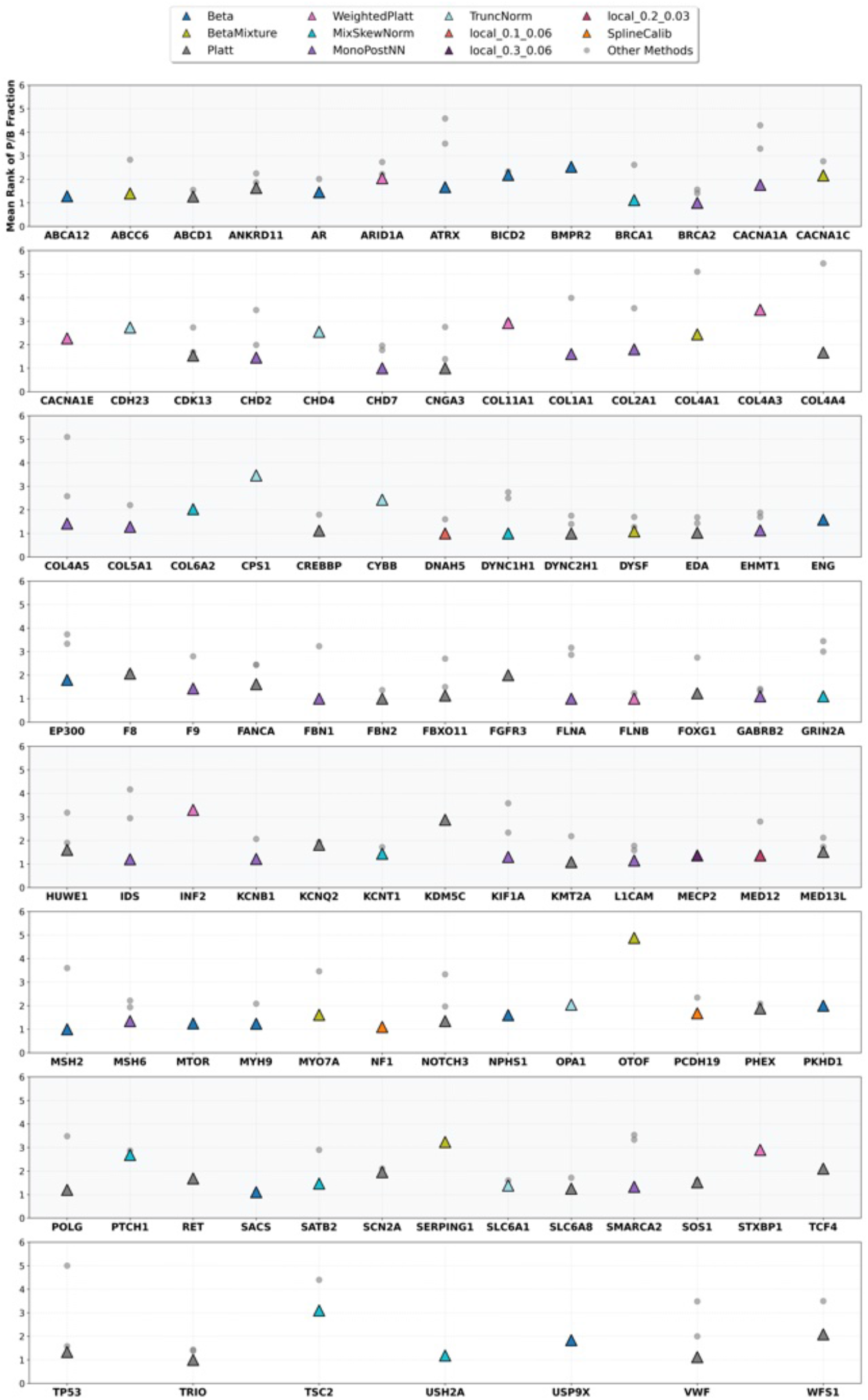
Mean rank by *P/B*_*Fraction*_ across calibration methods per gene (REVEL) Mean rank of the *P*_*Fraction*_ and *B*_*Fraction*_ metrics across calibration methods filtered by the first 2 steps of the Multi-stage method selection (Methods). Using the simulated score sets, a single calibration method was selected for each gene through a three-stage filtering procedure across 30 simulation datasets; only valid methods are retained in the figure. This is using REVEL score. For each gene, the final selected optimal calibration method is indicated by a colored triangle (with color corresponding to the method), while all other methods are shown as grey circles. Points represent 101 genes evaluated for REVEL.

**Extended Data Fig. 8:**
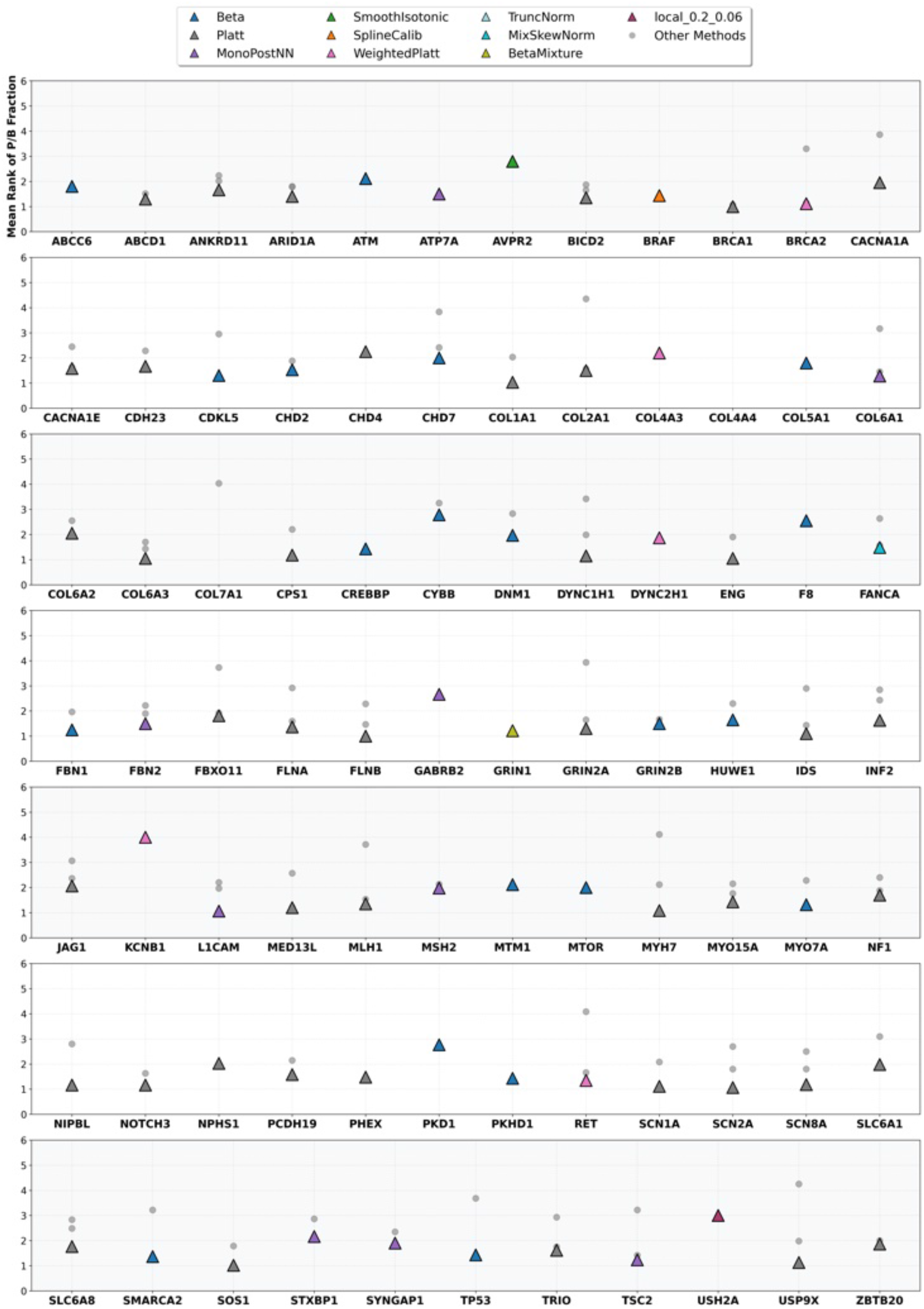
Mean rank by *P/B*_*Fraction*_ across calibration methods per gene (AlphaMissense) Mean rank of the *P*_*Fraction*_ and *B*_*Fraction*_ metrics across calibration methods filtered by the first 2 steps of the Multi-stage method selection (Methods). Using the simulated score sets, a single calibration method was selected for each gene through a three-stage filtering procedure across 30 simulation datasets; only valid methods are retained in the figure. This is using AlphaMissense score. For each gene, the final selected optimal calibration method is indicated by a colored triangle (with color corresponding to the method), while all other methods are shown as grey circles. Points represent 98 genes evaluated for AlphaMissense.

**Extended Data Fig. 9:**
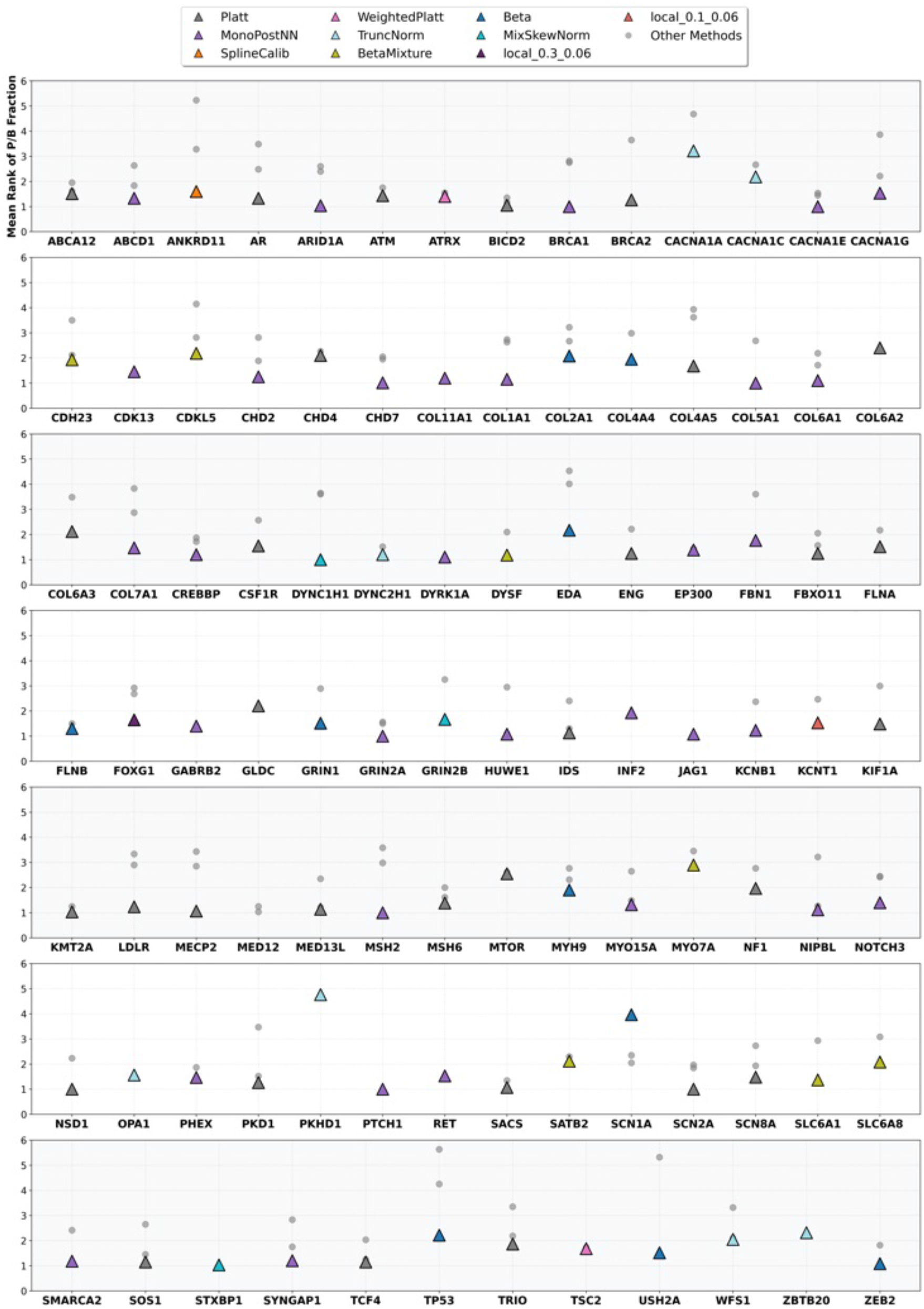
Mean rank by *P/B*_*Fraction*_ across calibration methods per gene (MutPred2) Mean rank of the *P*_*Fraction*_ and *B*_*Fraction*_ metrics across calibration methods filtered by the first 2 steps of the Multi-stage method selection (Methods). Using the simulated score sets, a single calibration method was selected for each gene through a three-stage filtering procedure across 30 simulation datasets; only valid methods are retained in the figure. This is using MutPred2 score. For each gene, the final selected optimal calibration method is indicated by a colored triangle (with color corresponding to the method), while all other methods are shown as grey circles. Points represent 105 genes evaluated for MutPred2.

**Extended Data Fig. 10:**
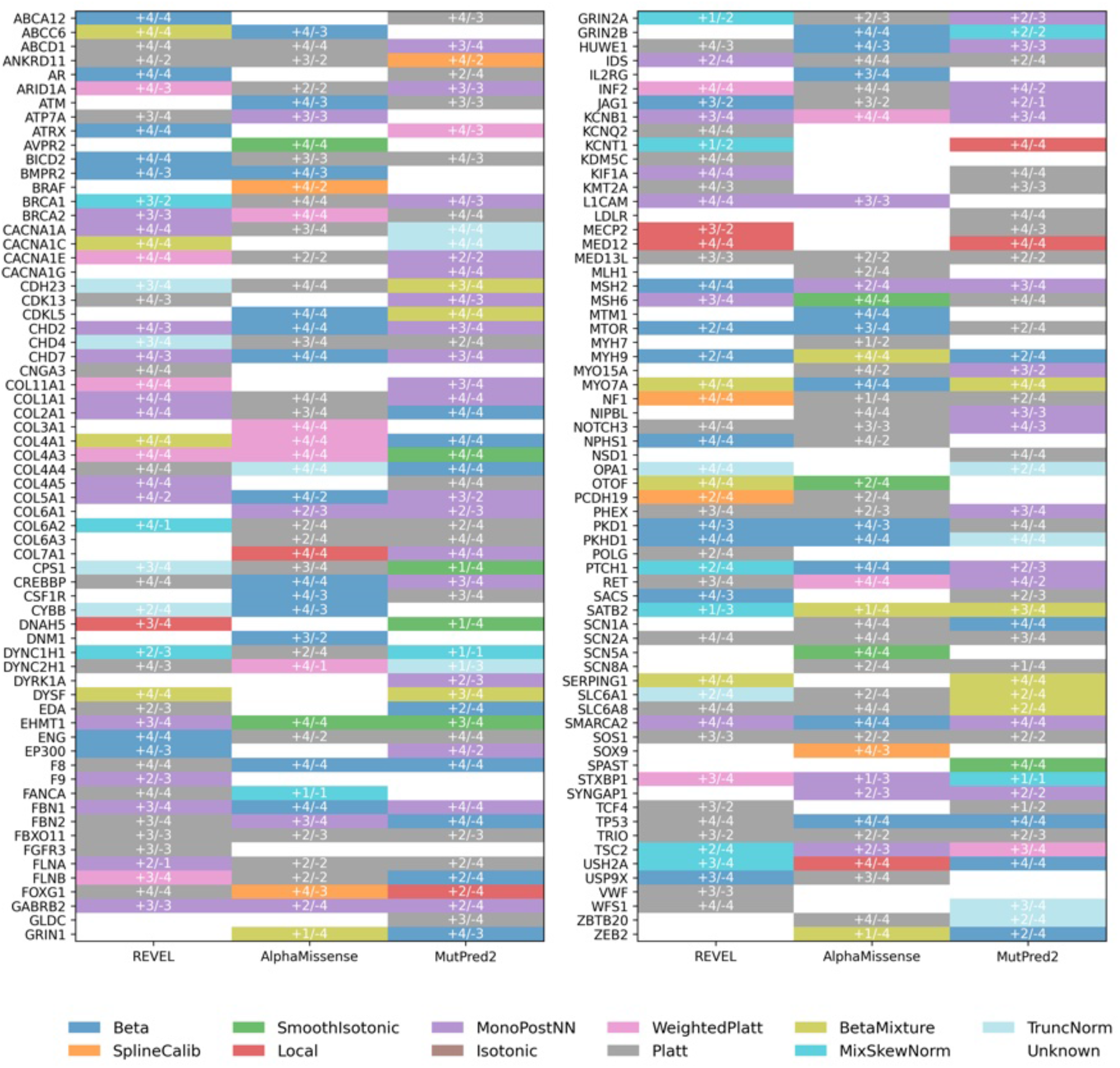
Best calibration method selected per gene across REVEL, AlphaMissense, and MutPred2. Best calibration method selected for each predictor across genes. Heatmap showing the optimal calibration method selected for each gene and predictor (REVEL, AlphaMissense and MutPred2). Rows represent genes (n = 132), sorted by gene name, and columns represent predictors. Colors denote the selected calibration method, and overlaid text indicates the corresponding benign (negative)/pathogenic (positive) evidence point assigned after calibration. Genes without a selected method are shown in white (Unknown). Across genes, method selection varied by predictor. For REVEL, Platt (32 genes), MonoPostNN (19 genes) and Beta (16 genes) were most frequently selected. For AlphaMissense, Platt was most common (42 genes), followed by Beta (24 genes), with 34 genes labeled Unknown. For MutPred2, MonoPostNN (32 genes) and Platt (31 genes) were most frequently selected, with 27 genes labeled Unknown.

**Extended Data Fig. 11:**
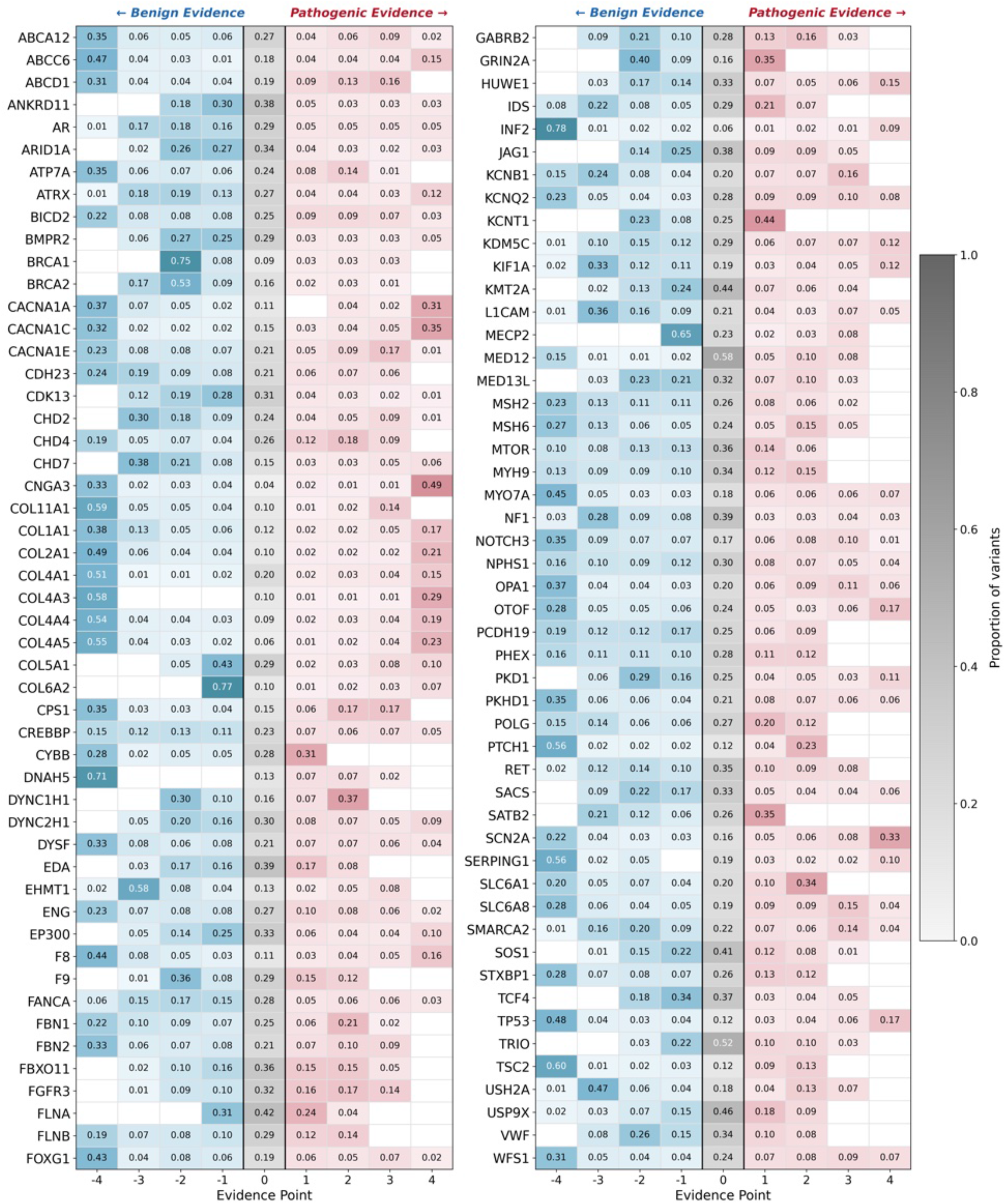
Evidence point assignment heatmap (REVEL) Gene-specific evidence point assignment proportions using REVEL scores. Each row represents a gene; each column represents an evidence level. Color intensity indicates variant proportion at each level: blue shades (-1 to -4) represent benign evidence; red shades (+1 to +4) represent pathogenic evidence; white (0) indicates indeterminate evidence. Blank cells denote proportions rounded to 0.00.

**Extended Data Fig. 12:**
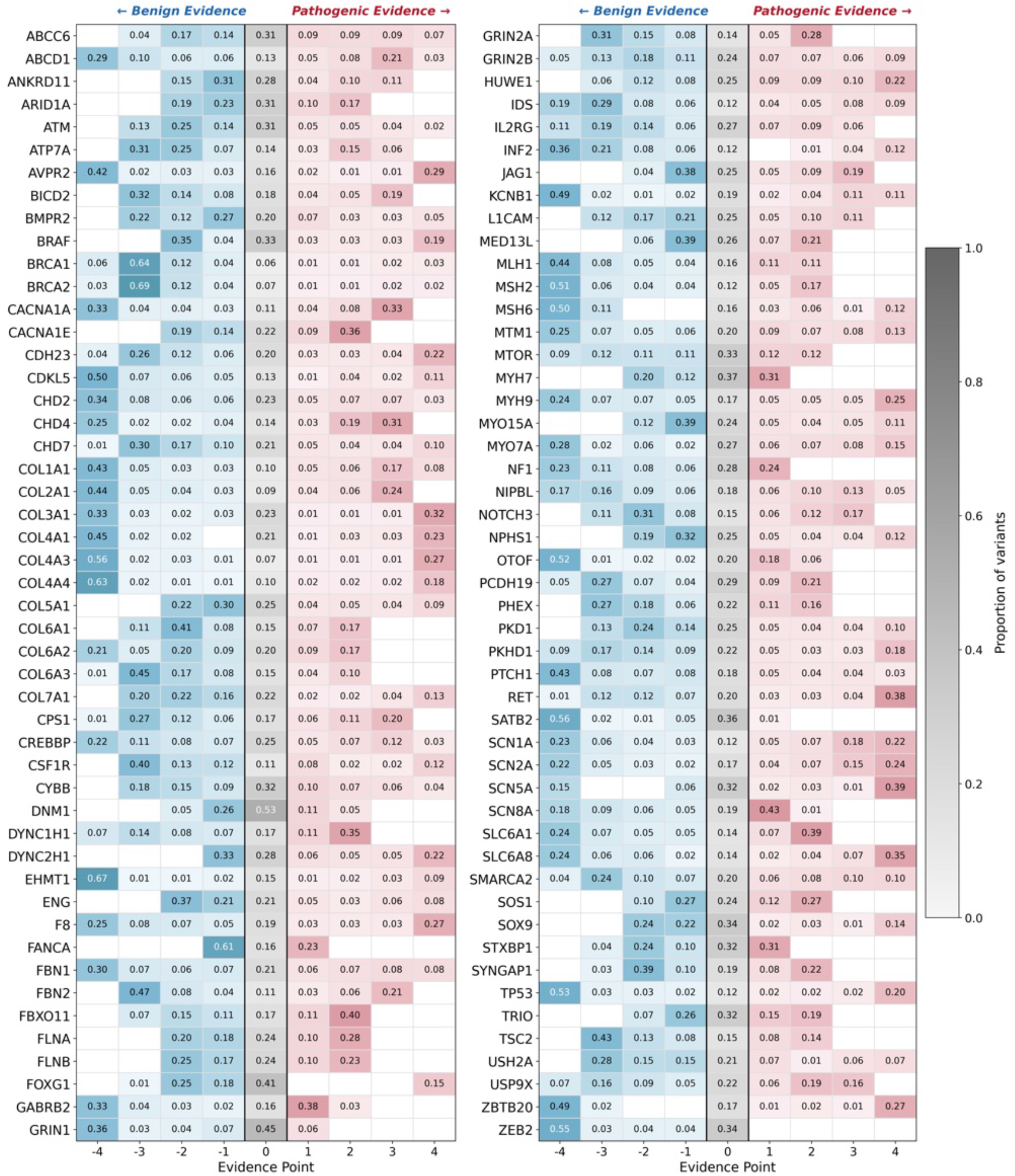
Evidence point assignment heatmap (AlphaMissense) Gene-specific evidence point assignment proportions using AlphaMissense scores. Each row represents a gene; each column represents an evidence level. Color intensity indicates variant proportion at each level: blue shades (-1 to -4) represent benign evidence; red shades (+1 to +4) represent pathogenic evidence; white (0) indicates indeterminate evidence. Blank cells denote proportions rounded to 0.00.

**Extended Data Fig. 13:**
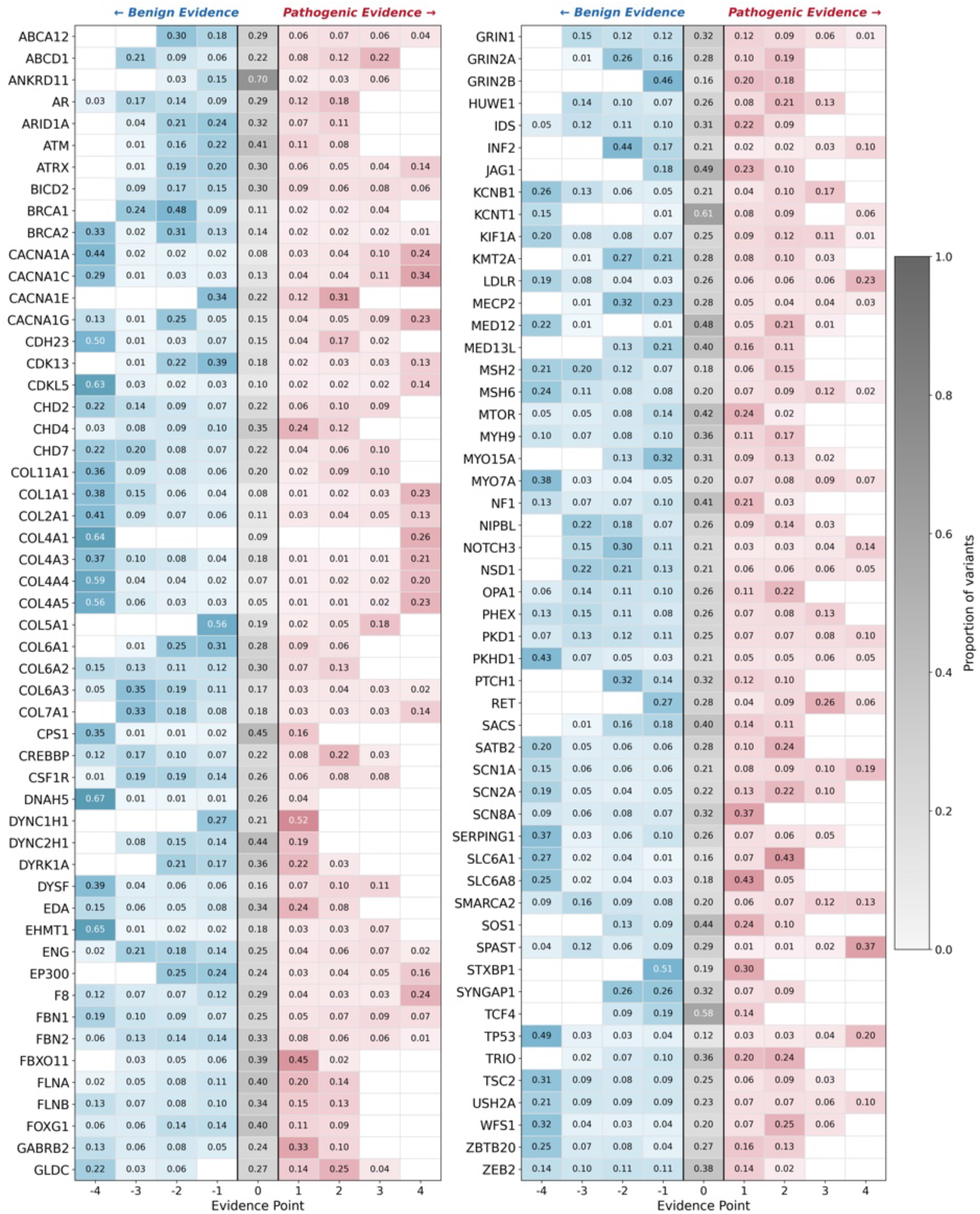
Evidence point assignment heatmap (MutPred2) Gene-specific evidence point assignment proportions using MutPred2 scores. Each row represents a gene; each column represents an evidence level. Color intensity indicates variant proportion at each level: blue shades (-1 to -4) represent benign evidence; red shades (+1 to +4) represent pathogenic evidence; white (0) indicates indeterminate evidence. Blank cells denote proportions rounded to 0.00.

**Extended Data Fig. 14:**
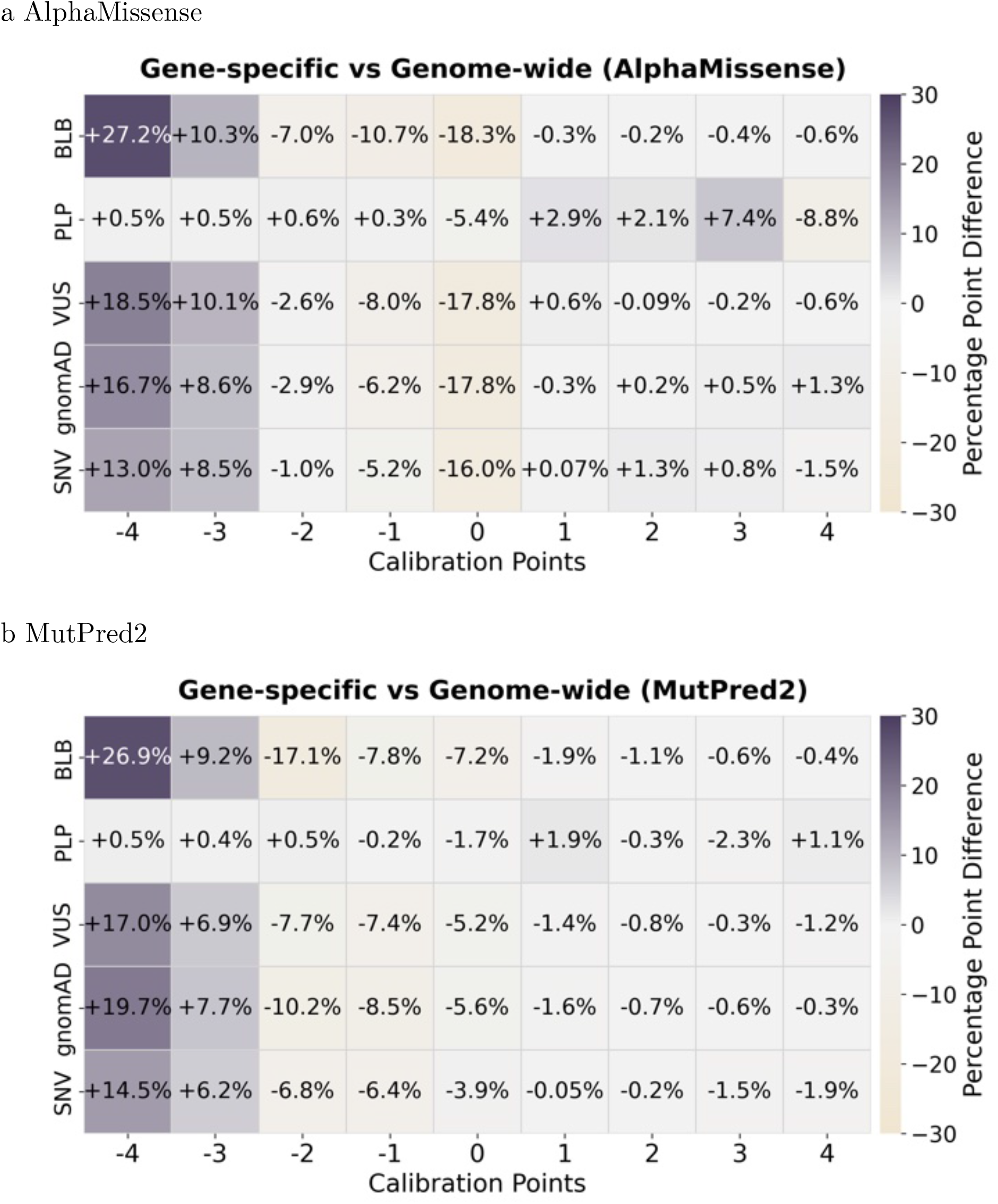
Evidence point assignment difference (gene-specific vs. genome-wide): AlphaMissense and MutPred2. Comparison of evidence point assignments between gene-specific and genome-wide calibration methods. (a) Results using AlphaMissense scores. (b) Results using MutPred2 scores. Heatmaps display the differences in the percentage of evidence point assignments stratified by variant classification or source. Grey indicates increased assignment rates by the gene-specific calibration method compared to genome-wide calibration.

**Extended Data Fig. 15:**
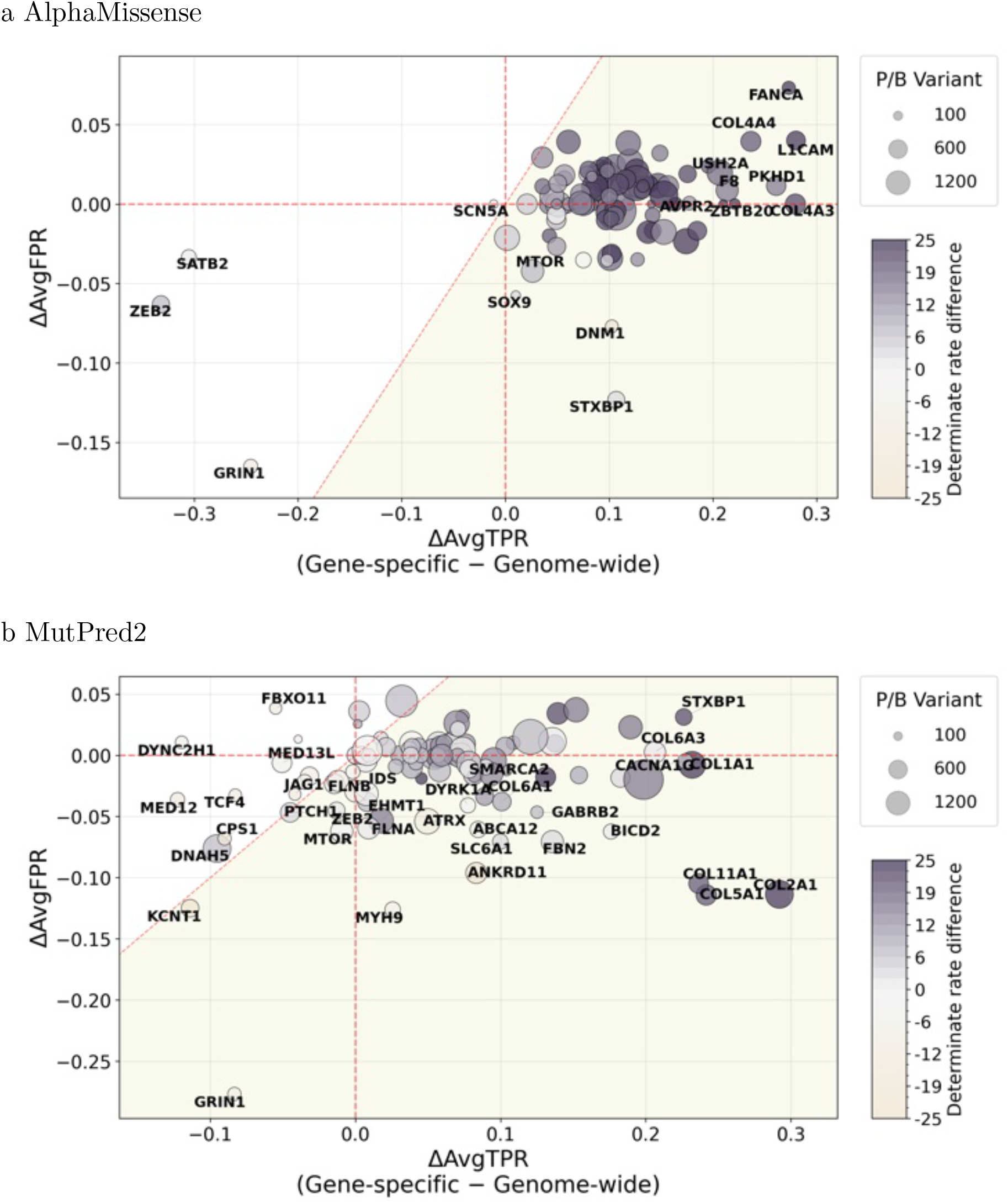
Calibration method comparison: AlphaMissense and MutPred2 (per-gene) Gene-level changes in sensitivity and false positive rate following gene-specific calibration. (a) Results using AlphaMissense scores. (b) Results using MutPred2 scores. Each point represents a gene. The x-axis shows the average change in true positive rate (ΔAvgTPR) between gene-specific and genome-wide calibration (gene-specific minus aggregated), and the y-axis shows the corresponding change in false positive rate (ΔAvgFPR). Point color encodes the change in evidence coverage (fraction of variants receiving at least -/+1 point of evidence), with deeper grey indicating higher coverage under the gene-specific method. Point size is proportional to the number of pathogenic and benign variants (*n*_PB_; P/LP and B/LB). The shaded area indicates increased performance (below the dashed red diagonal *x* = *y*).

**Extended Data Fig. 16:**
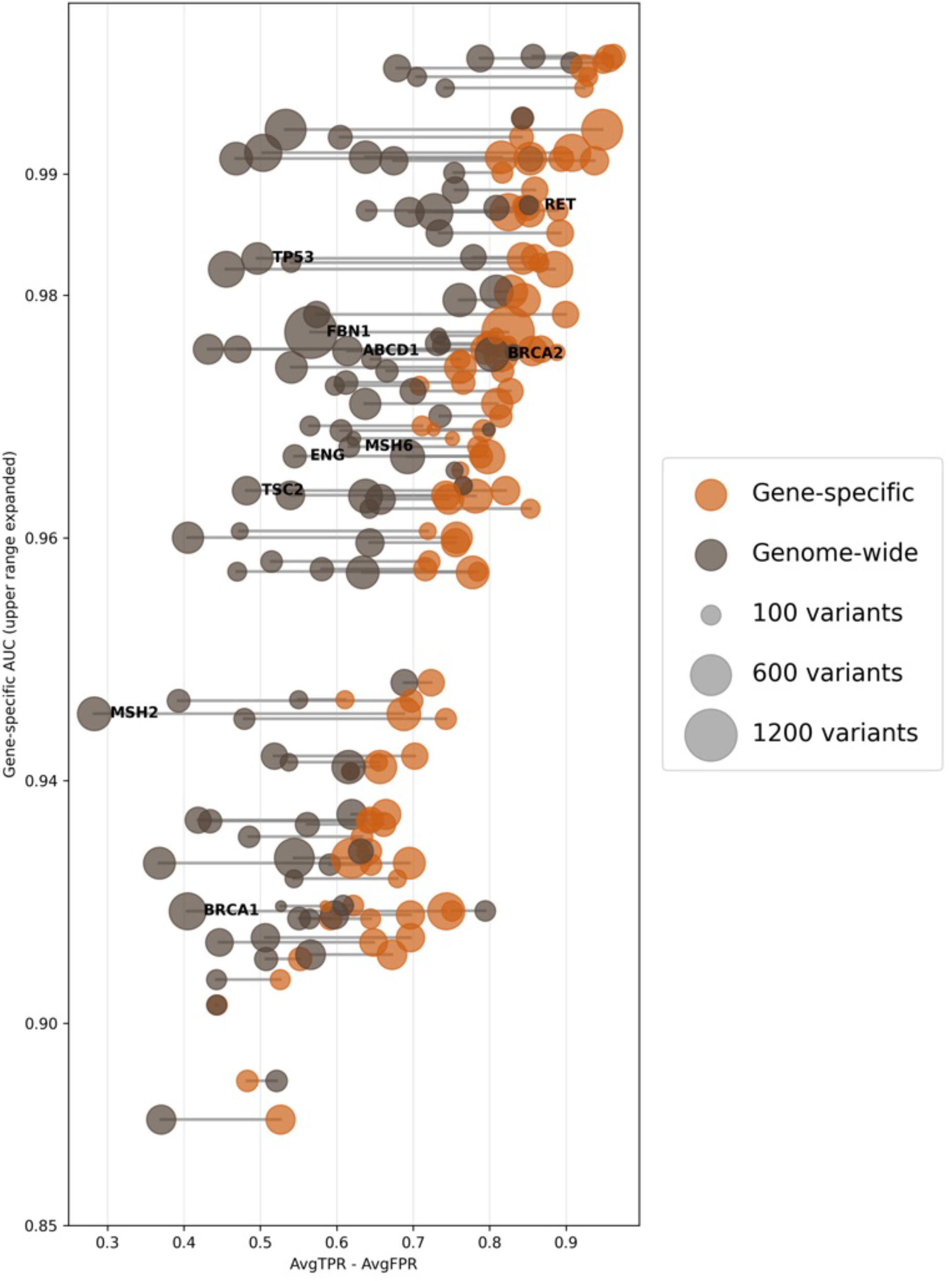
Gene-specific calibration performance and REVEL AUC. Gene-specific performance (AvgTPR − AvgFPR) versus gene-specific REVEL AUC shown as a dumbbell plot. Each gene is represented by a pair of connected points (gene-specific calibration vs. genome-wide calibration). The x-axis shows the gene-specific performance metric (AvgTPR − AvgFPR), and the y-axis shows the gene-specific REVEL AUC. Genes annotated with labels correspond to ACMG secondary finding genes. Higher values of AvgTPR − AvgFPR indicate better discrimination performance. The y-axis is displayed on a non-linear scale, with the upper range expanded to improve visualization of di!erences across genes.

**Extended Data Fig. 17:**
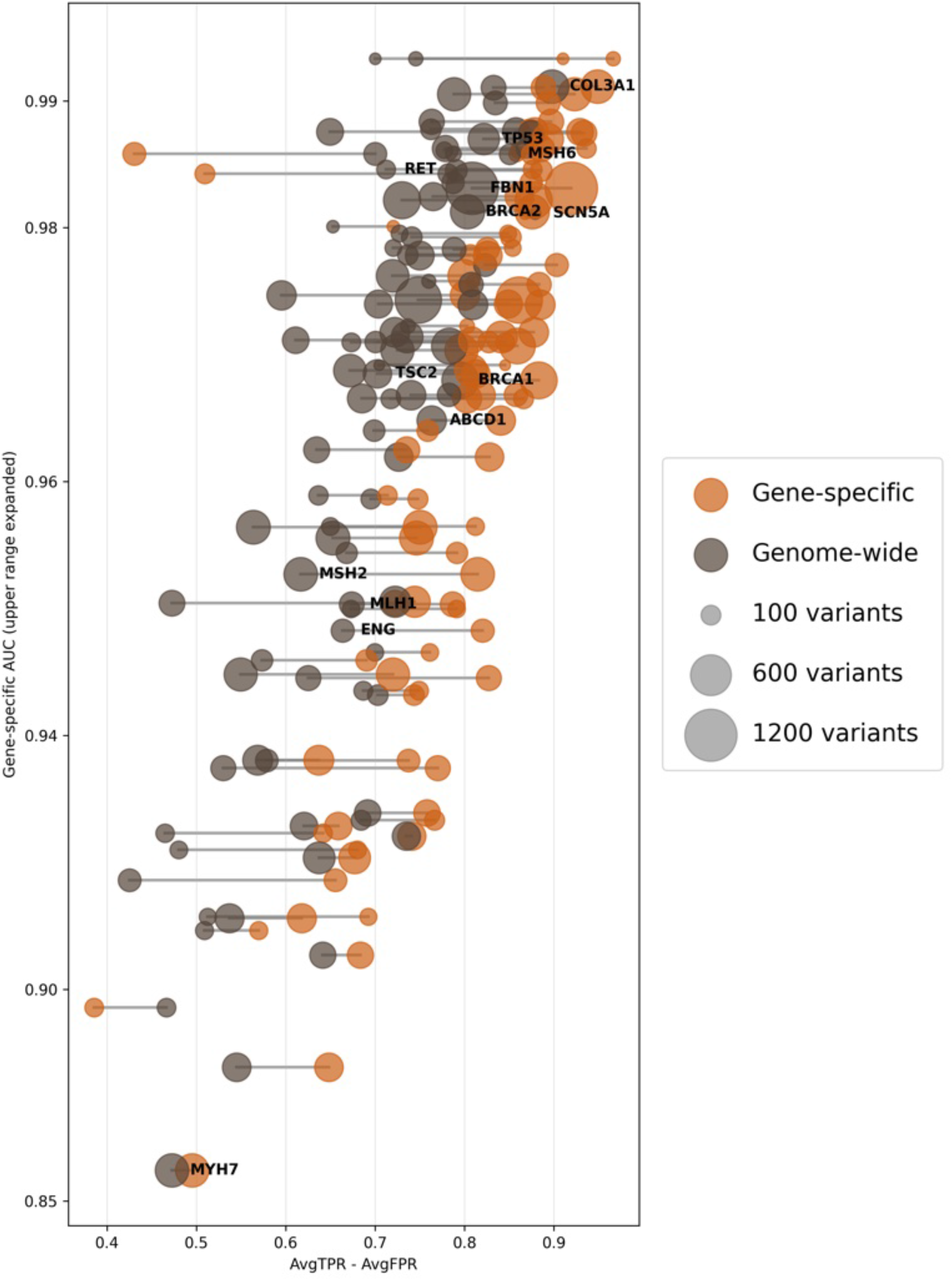
Gene-specific calibration performance and AlphaMissense AUC. Gene-specific performance (AvgTPR − AvgFPR) versus gene-specific AlphaMissense AUC shown as a dumbbell plot. Each gene is represented by a pair of connected points (gene-specific calibration vs. genome-wide calibration). The x-axis shows the gene-specific performance metric (AvgTPR − AvgFPR), and the y-axis shows the gene-specific AlphaMissense AUC. Genes annotated with labels correspond to ACMG secondary finding genes. Higher values of AvgTPR − AvgFPR indicate better discrimination performance. The y-axis is displayed on a non-linear scale, with the upper range expanded to improve visualization of di!erences across genes.

**Extended Data Fig. 18:**
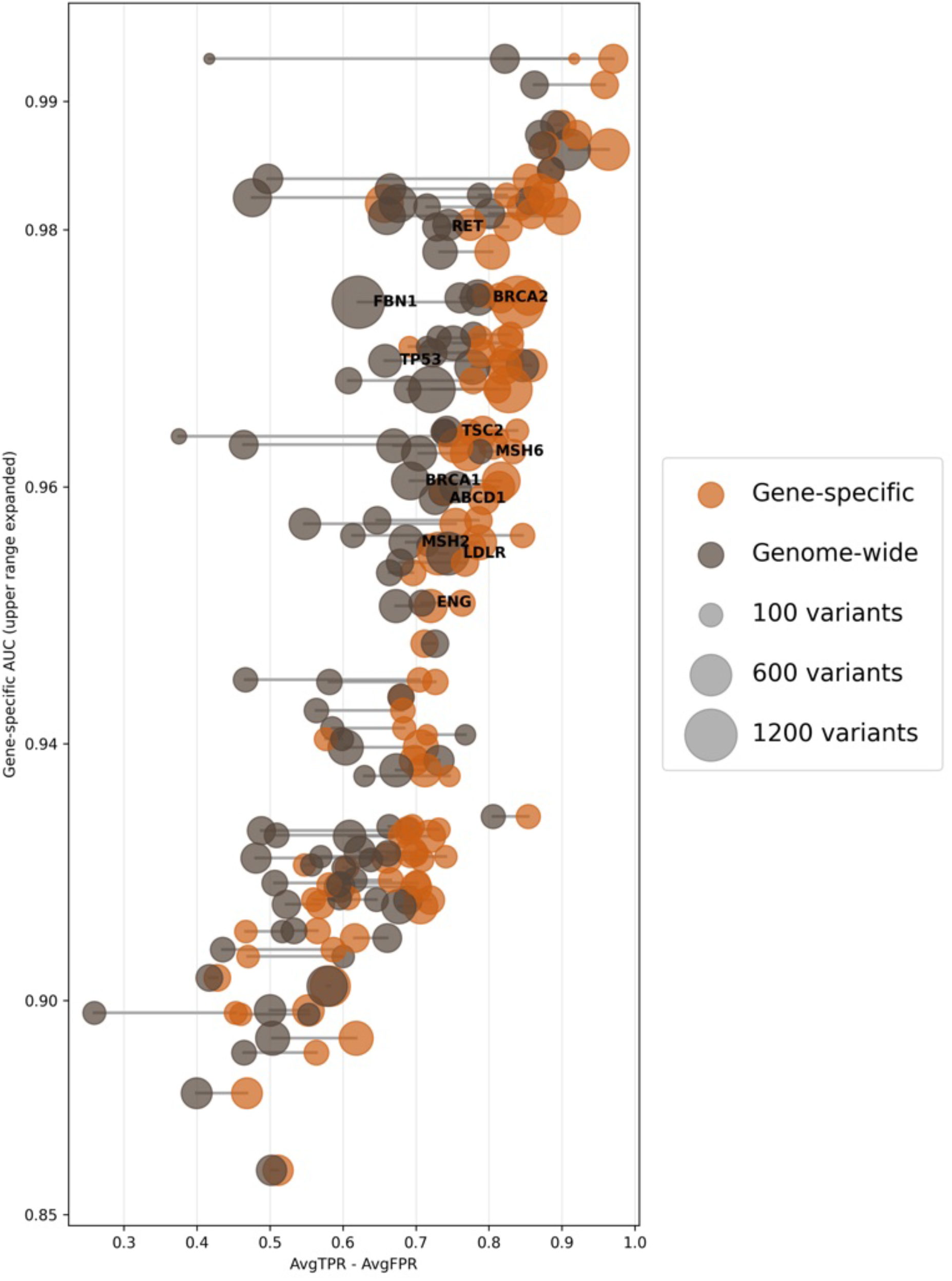
Gene-specific calibration performance and MutPred2 AUC. Gene-specific performance (AvgTPR − AvgFPR) versus gene-specific MutPred2 AUC shown as a dumbbell plot. Each gene is represented by a pair of connected points (gene-specific calibration vs. genome-wide calibration). The x-axis shows the gene-specific performance metric (AvgTPR − AvgFPR), and the y-axis shows the gene-specific MutPred2 AUC. Genes annotated with labels correspond to ACMG secondary finding genes. Higher values of AvgTPR − AvgFPR indicate better discrimination performance. The y-axis is displayed on a non-linear scale, with the upper range expanded to improve visualization of di!erences across genes.

**Extended Data Fig. 19:**
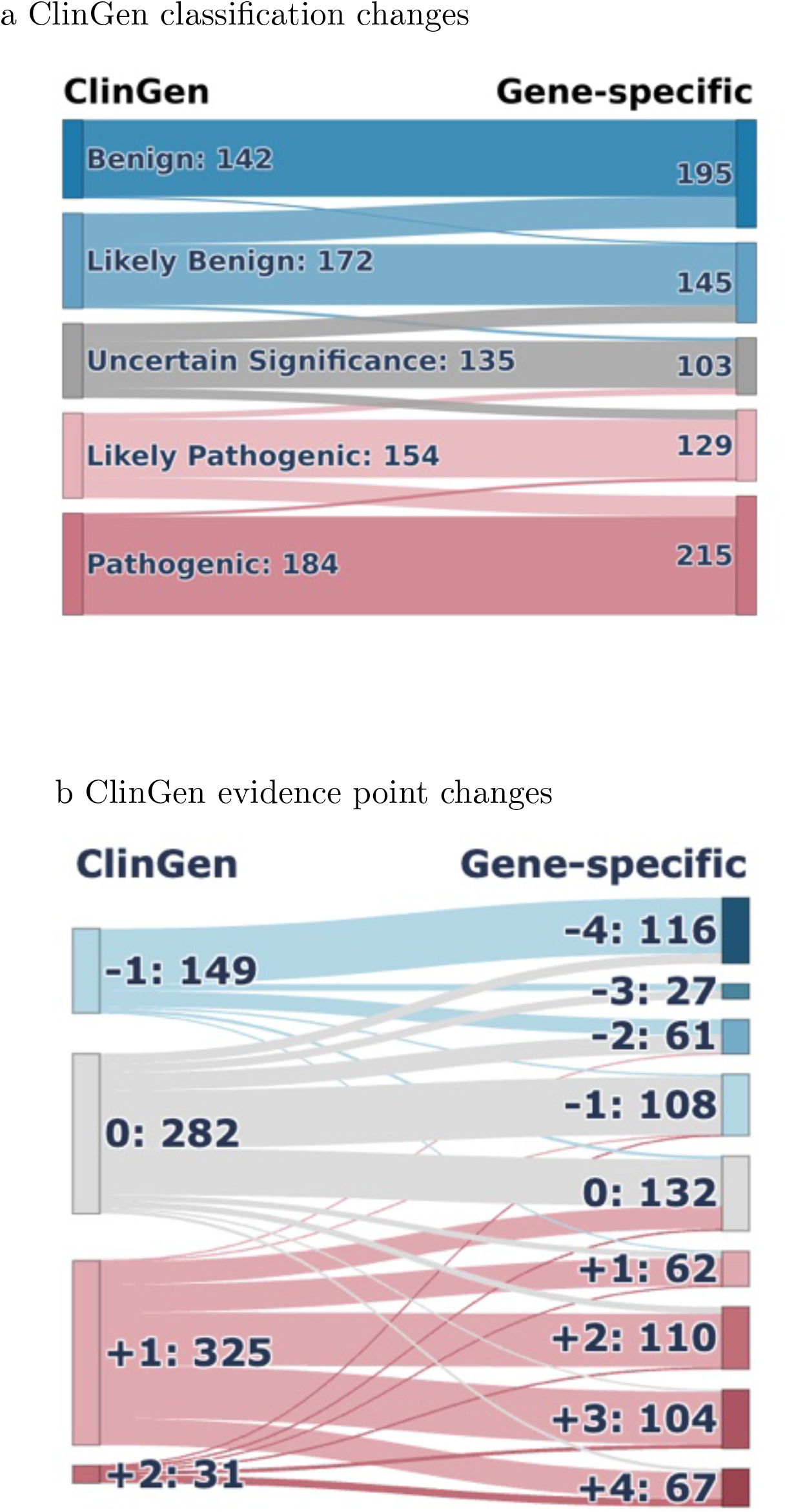
ClinGen Sankey by gene-specific calibration (REVEL) Sankey diagrams illustrating the impact of gene-specific calibration using REVEL scores. (a) Transitions in clinical classifications from original ClinGen assignments to recalibrated classifications. (b) Reassignment of computational evidence points under ClinGen guidelines compared with gene-specific calibration.

**Extended Data Fig. 20:**
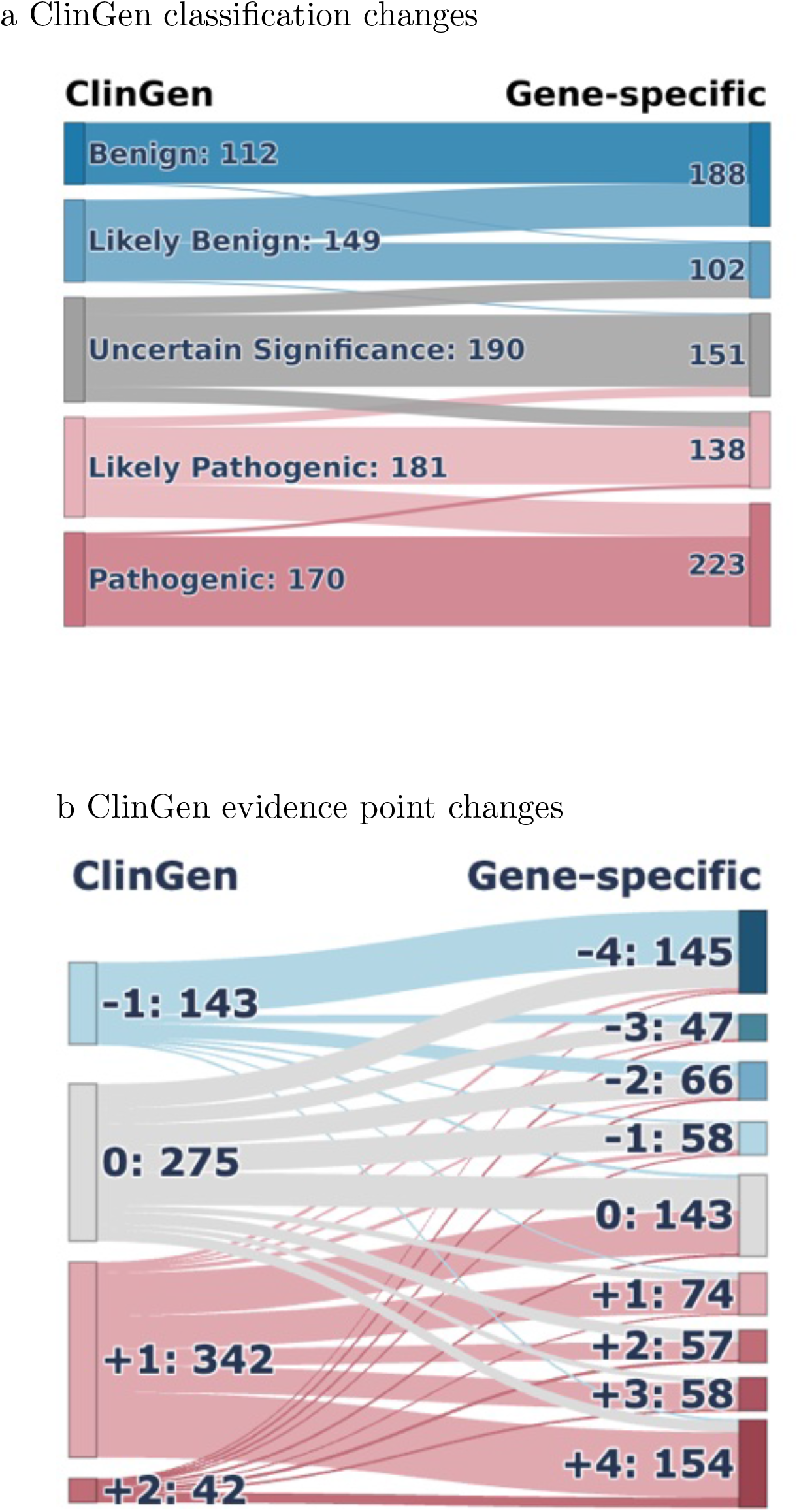
ClinGen Sankey by gene-specific calibration (AlphaMissense) Sankey diagrams illustrating the impact of gene-specific calibration using AlphaMissense scores. (a) Transitions in clinical classifications from original ClinGen assignments to recalibrated classifications. (b) Reassignment of computational evidence points under ClinGen guidelines compared with gene-specific calibration.

**Extended Data Fig. 21:**
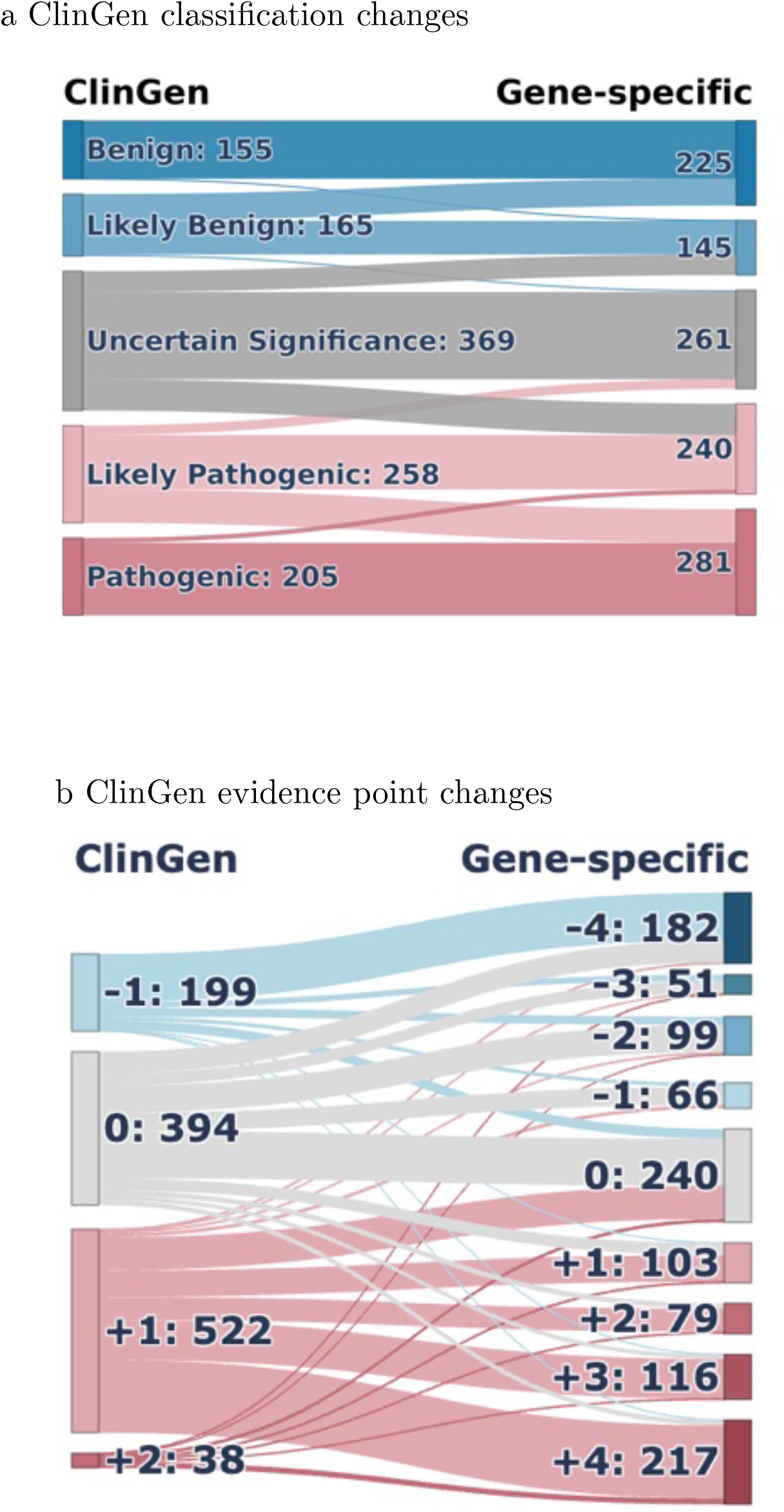
ClinGen Sankey by gene-specific calibration (MutPred2) Sankey diagrams illustrating the impact of gene-specific calibration using MutPred2 scores. (a) Transitions in clinical classifications from original ClinGen assignments to recalibrated classifications. (b) Reassignment of computational evidence points under ClinGen guidelines compared with gene-specific calibration.

**Extended Data Fig. 22:**
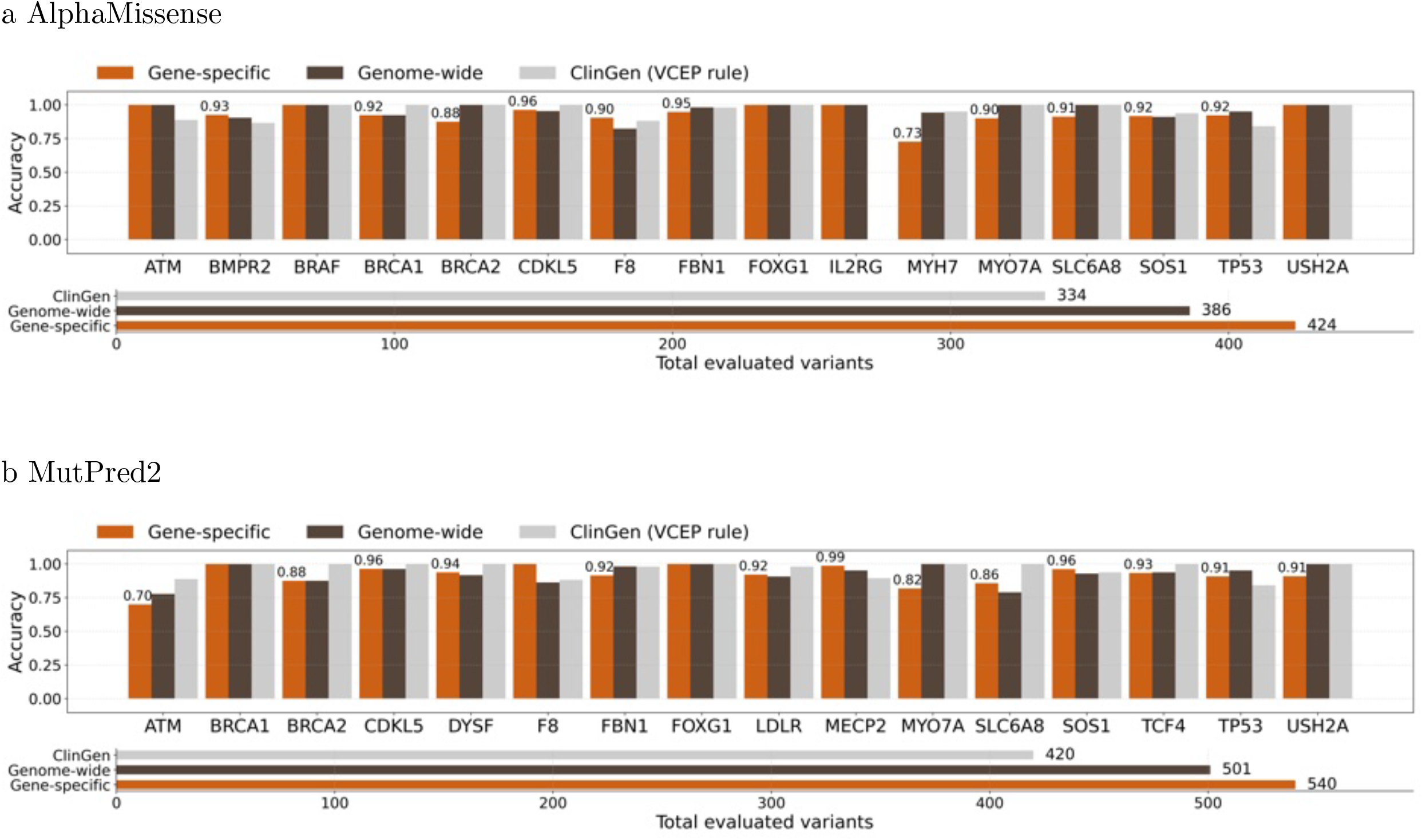
ClinGen non-circular set evaluation for gene-specific calibration (excluding zero-point assignments) Evaluation of per-gene classification accuracy on the ClinGen non-circular variant set using (a) AlphaMissense and (b) MutPred2 scores under three approaches: ClinGen-provided computational evidence strengths, genome-wide calibration thresholds, and gene-specific calibration thresholds. Top bar plots show the fraction of variants correctly classified relative to ClinGen reference classifications after recomputing variant classifications using only non-computational evidence to maintain non-circularity. Variants assigned zero computational evidence points are excluded from accuracy calculations. Bottom bar plots show the total number of variants assigned non-zero computational evidence points by each method.

**Extended Data Fig. 23:**
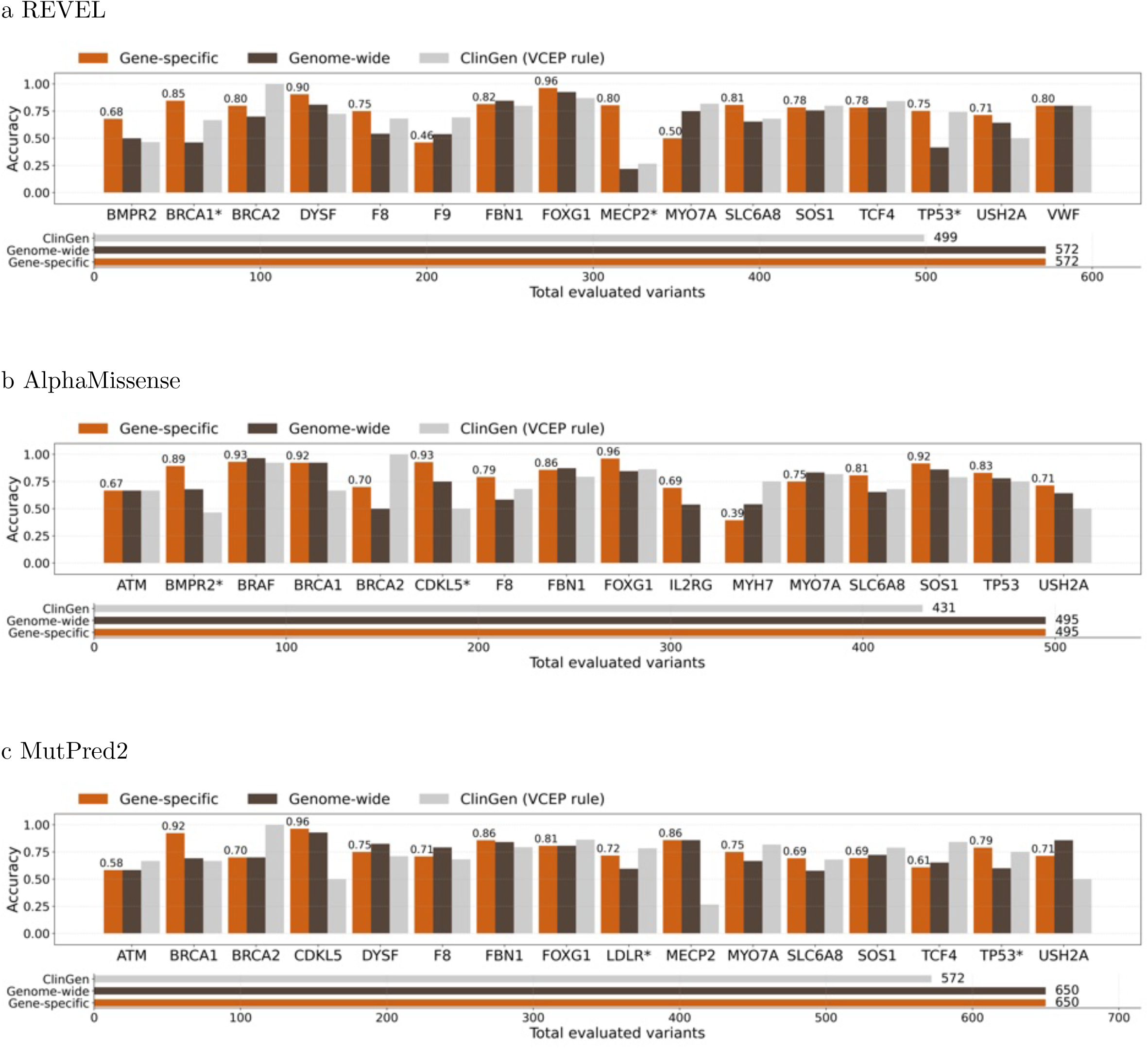
ClinGen non-circular set evaluation for gene-specific calibration (including zero-point assignments) Evaluation of per-gene classification accuracy on the ClinGen non-circular variant set using (a) REVEL, (b) AlphaMissense, and (c) MutPred2 scores under three approaches: ClinGen-provided computational evidence strengths, genome-wide calibration thresholds, and gene-specific calibration thresholds. Top bar plots show the fraction of variants correctly classified relative to ClinGen reference classifications after recomputing variant classifications using only non-computational evidence to maintain non-circularity. Variants assigned zero computational evidence points are included in the accuracy calculations. Bottom bar plots show the total number of variants assigned computational evidence points by each method.

**Extended Data Fig. 24:**
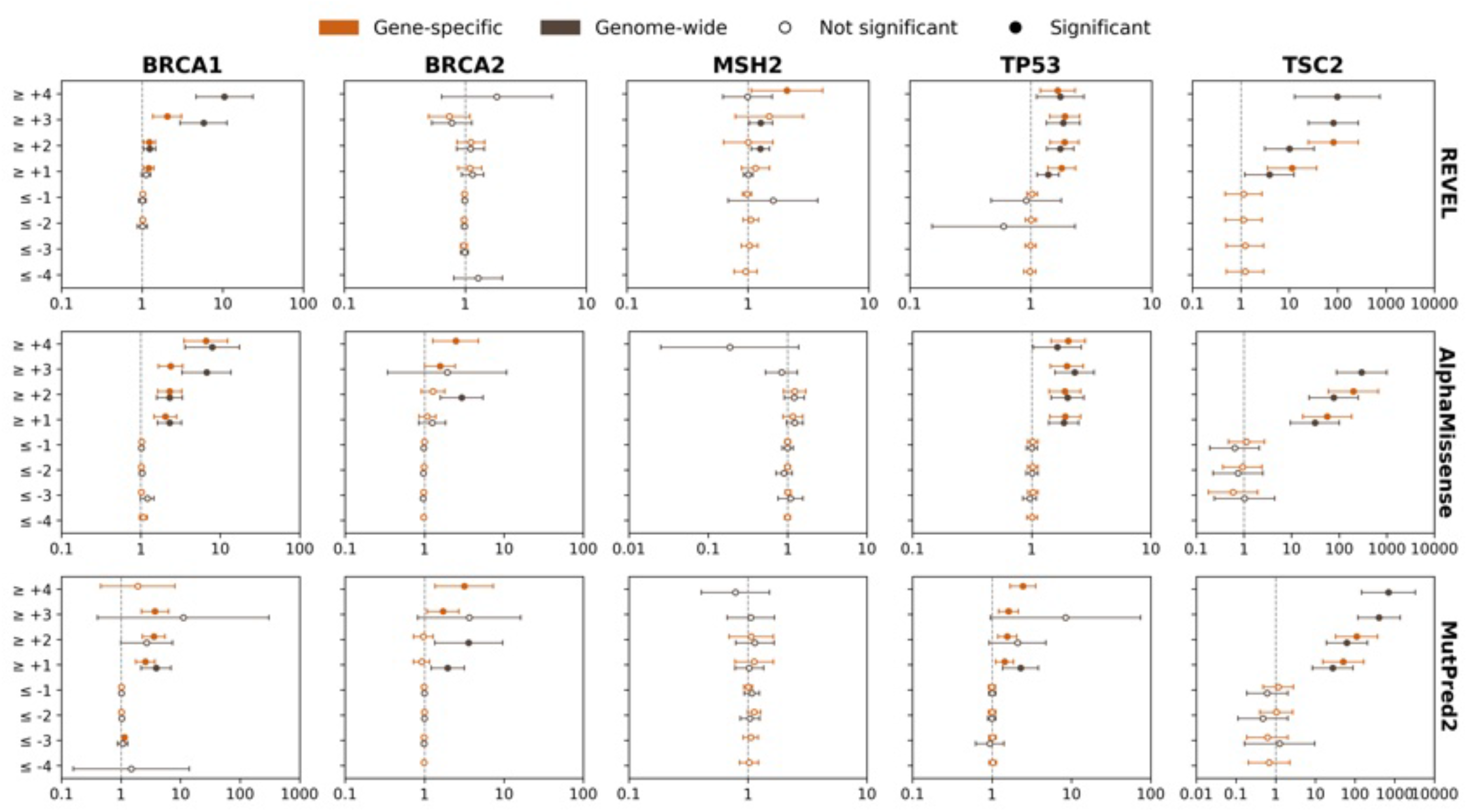
Odds ratios for disease occurrence in the All of Us biobank. Odds ratios for disease occurrence in the All of Us biobank for variants meeting different evidence strength thresholds in example genes using gene-specific calibration compared with genome-wide calibration. The x-axis shows the odds ratio (vertical dashed line indicates OR = 1), and the y-axis shows total evidence points for variant sets. Circles represent estimated odds ratios with 95% confidence intervals (whiskers); filled circles indicate statistically significant associations.

**Extended Data Fig. 25:**
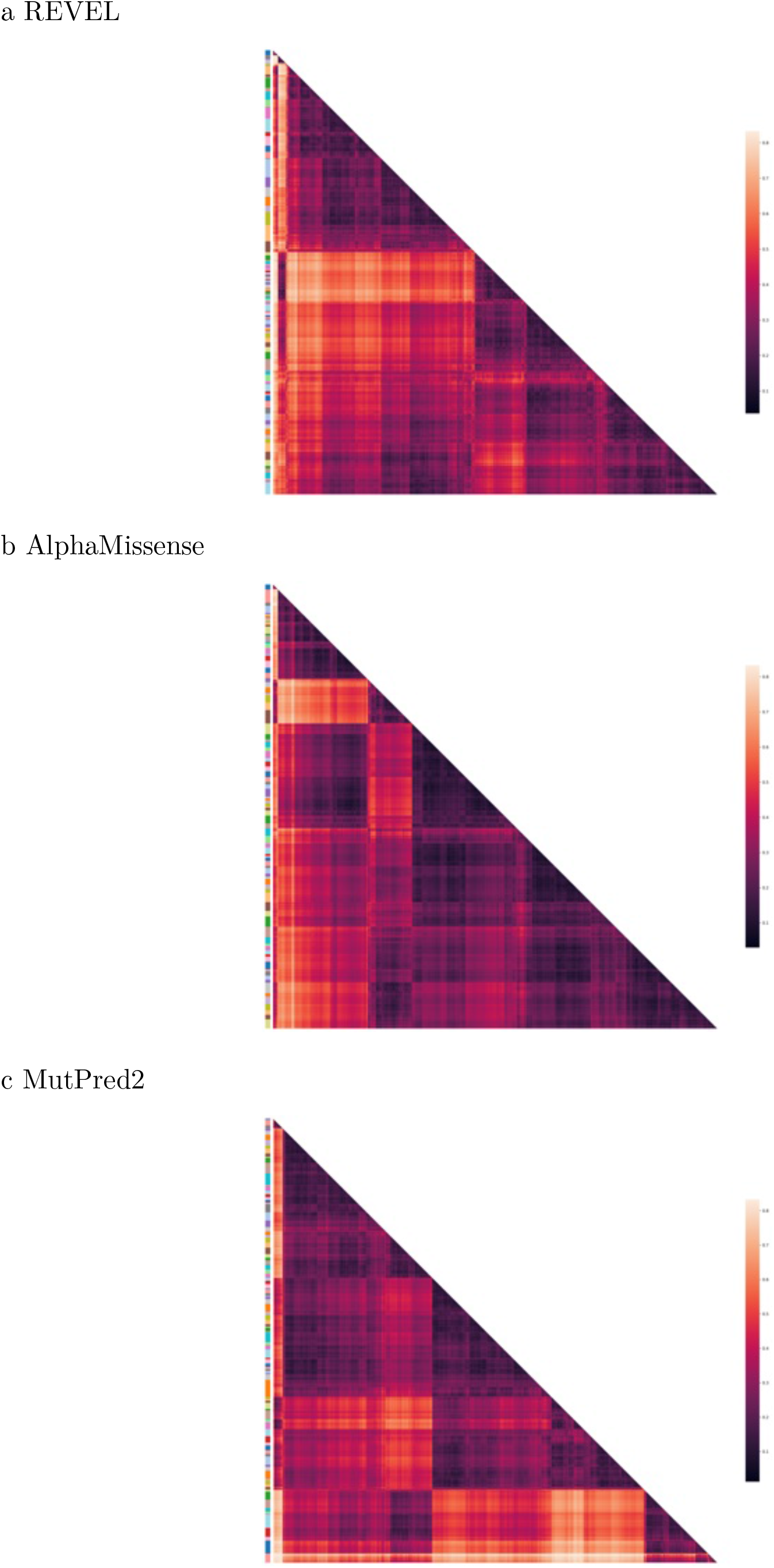
Domain-based clustering heatmap. Hierarchical clustering of domain score distributions for three variant effect predictors. Heatmaps show the full hierarchical clustering of domains based on Jensen–Shannon distance between score distributions, with sidebar colors indicating cluster assignments. (a) REVEL (98 clusters). (b) AlphaMissense (92 clusters). (c) MutPred2 (102 clusters).

**Extended Data Fig. 26:**
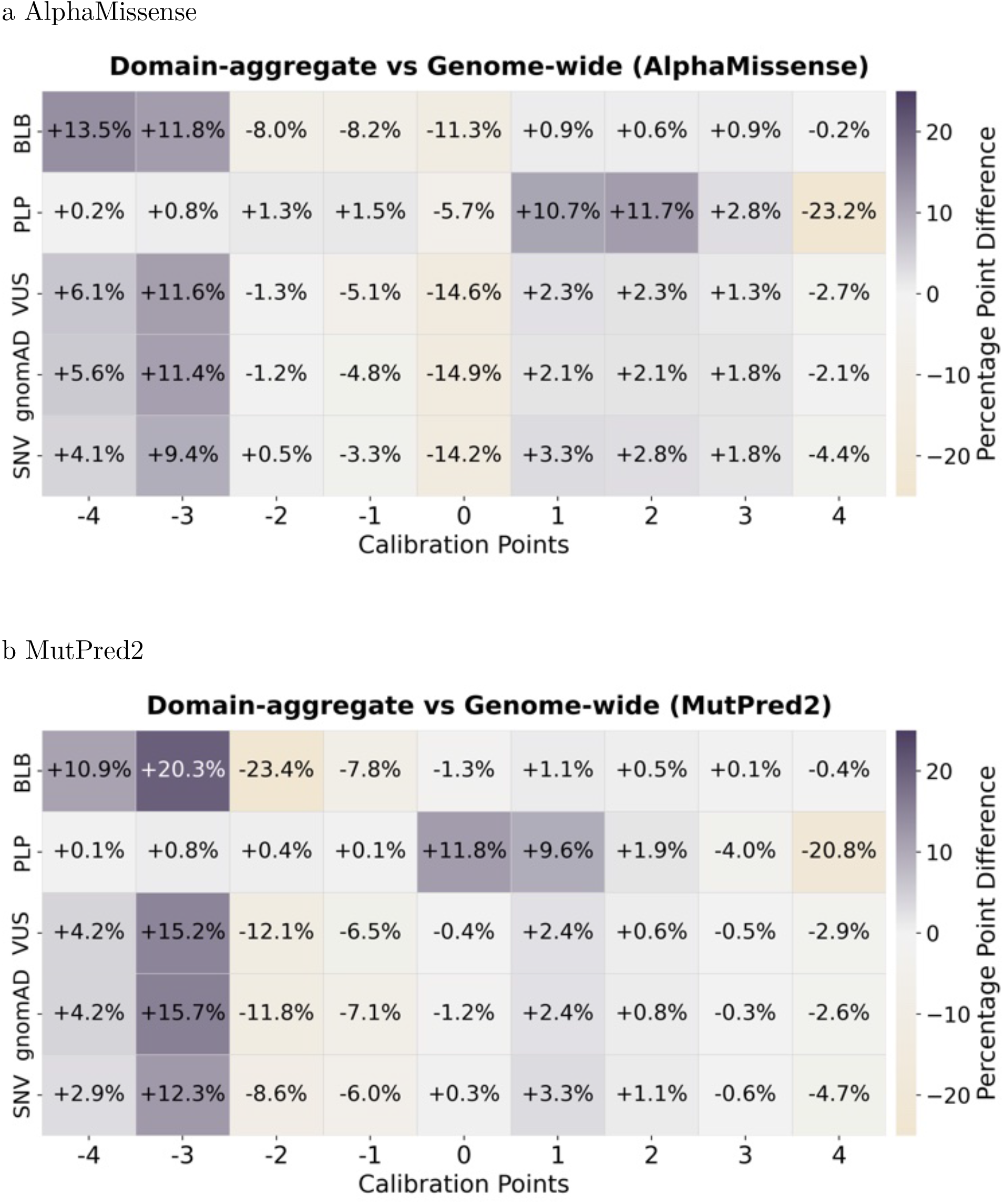
Evidence point assignment difference (domain-aggregate vs. genome-wide): AlphaMissense and MutPred2. Comparison of evidence point assignments between domain-aggregate and genome-wide calibration methods. (a) Results using AlphaMissense scores. (b) Results using MutPred2 scores. Heatmaps display the differences in the percentage of evidence point assignments stratified by variant classification or source. Grey indicates increased assignment rates by the domain-aggregate calibration method compared to genome-wide calibration.

**Extended Data Fig. 27:**
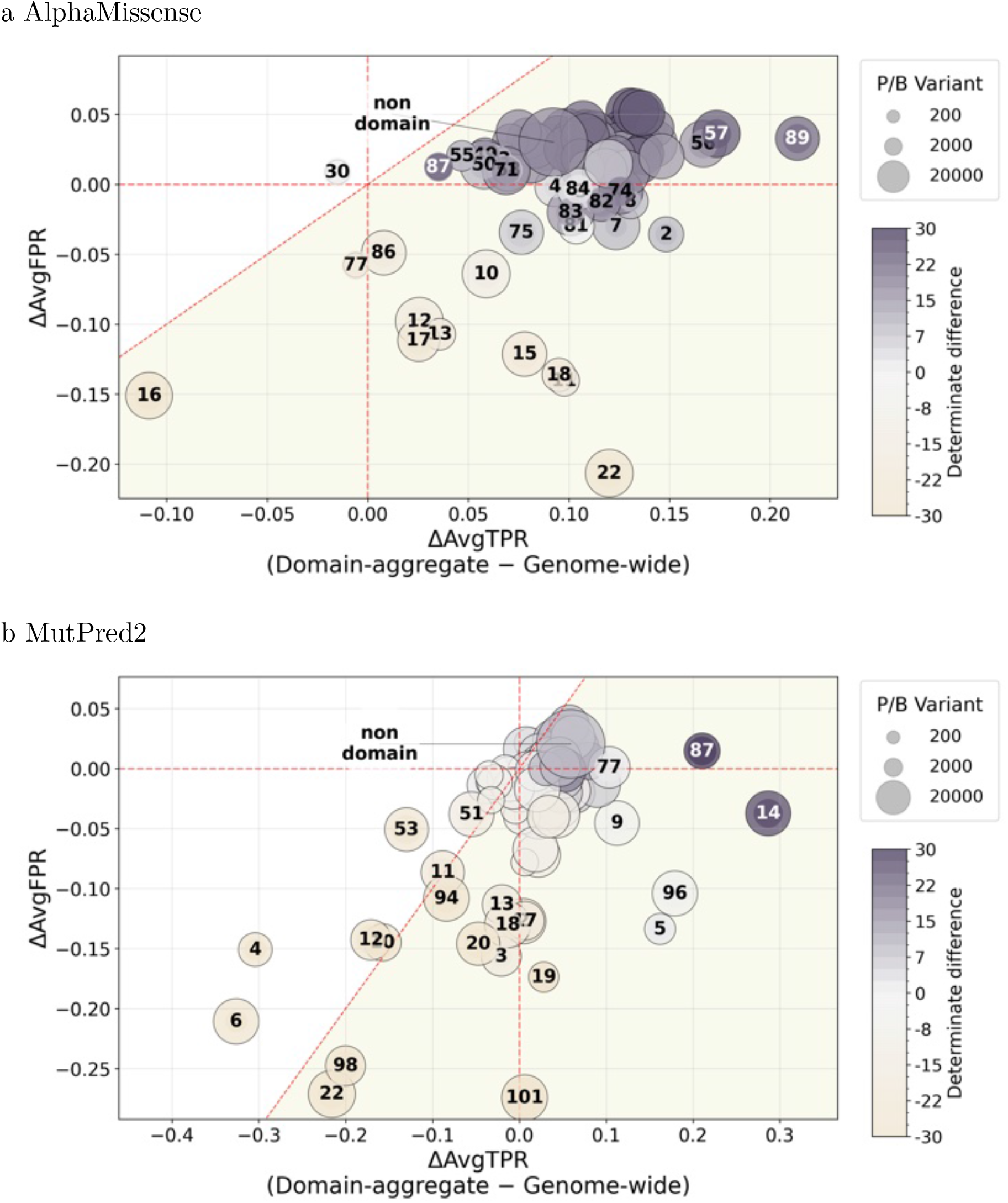
Calibration method comparison: AlphaMissense and MutPred2 (per-cluster) Cluster-level changes in sensitivity and false positive rate following domain-aggregate calibration. (a) Results using AlphaMissense scores. (b) Results using MutPred2 scores. Each point represents a cluster. The x-axis shows the average change in true positive rate (ΔAvgTPR) between domain-aggregate and genome-wide calibration (domain-aggregate minus genome-wide), and the y-axis shows the corresponding change in false positive rate (ΔAvgFPR). Point color encodes the change in evidence coverage (fraction of variants receiving at least -/+1 point of evidence), with deeper grey indicating higher coverage under the domain-aggregate method. Point size is proportional to the number of pathogenic and benign variants (*n*_PB_; P/LP and B/LB). The shaded area indicates increased performance (below the dashed red diagonal *x* = *y*).

**Extended Data Fig. 28:**
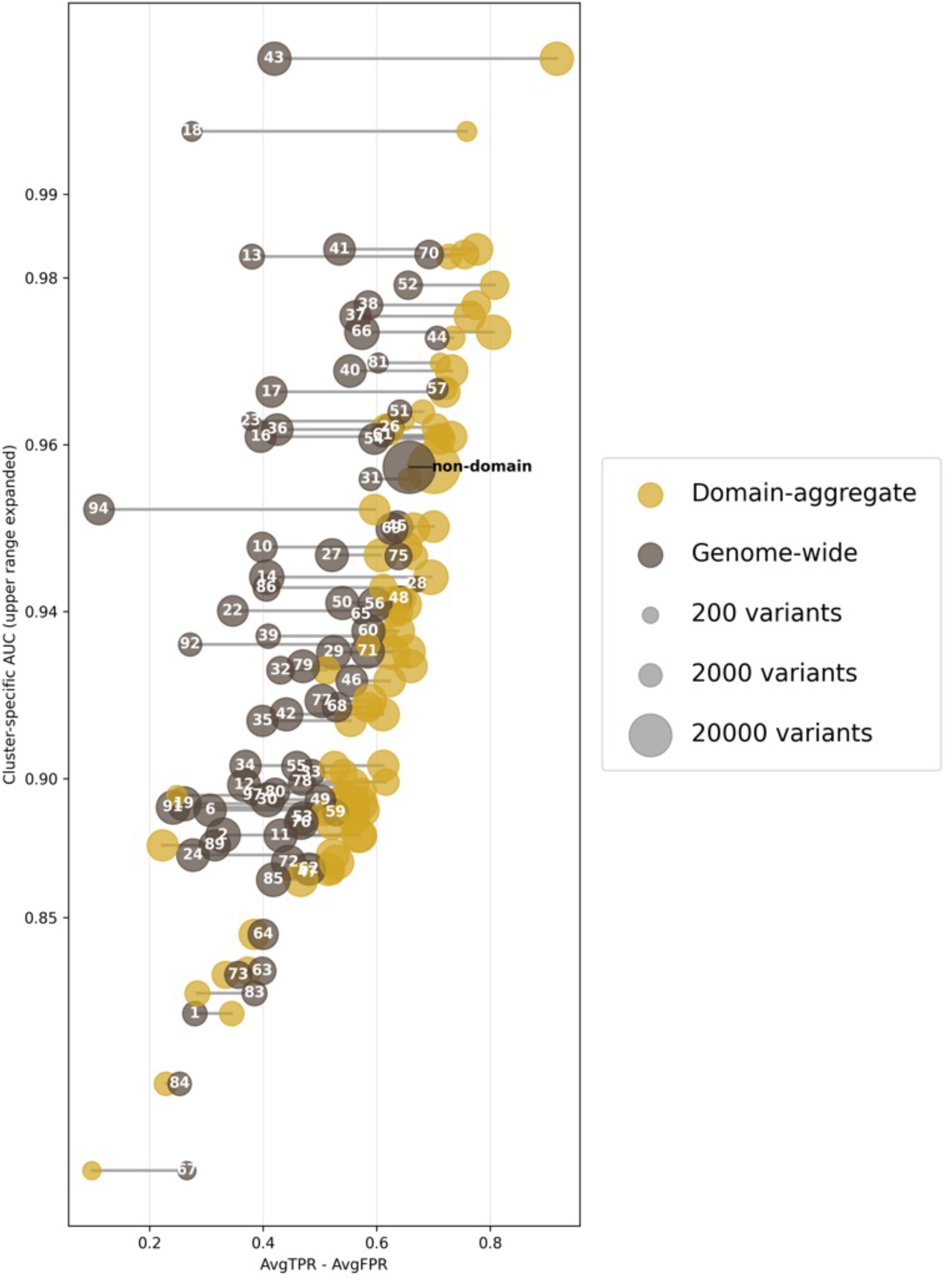
Cluster-specific calibration performance and REVEL AUC. Cluster-specific performance (AvgTPR − AvgFPR) versus cluster-specific REVEL AUC shown as a dumbbell plot. Each cluster is represented by a pair of connected points (domain-aggregate calibration vs. genome-wide calibration). The x-axis shows the domain-aggregate performance metric (AvgTPR − AvgFPR), and the y-axis shows the cluster-specific REVEL AUC. Higher values of AvgTPR − AvgFPR indicate better discrimination performance. The y-axis is displayed on a non-linear scale, with the upper range expanded to improve visualization of differences across clusters.

**Extended Data Fig. 29:**
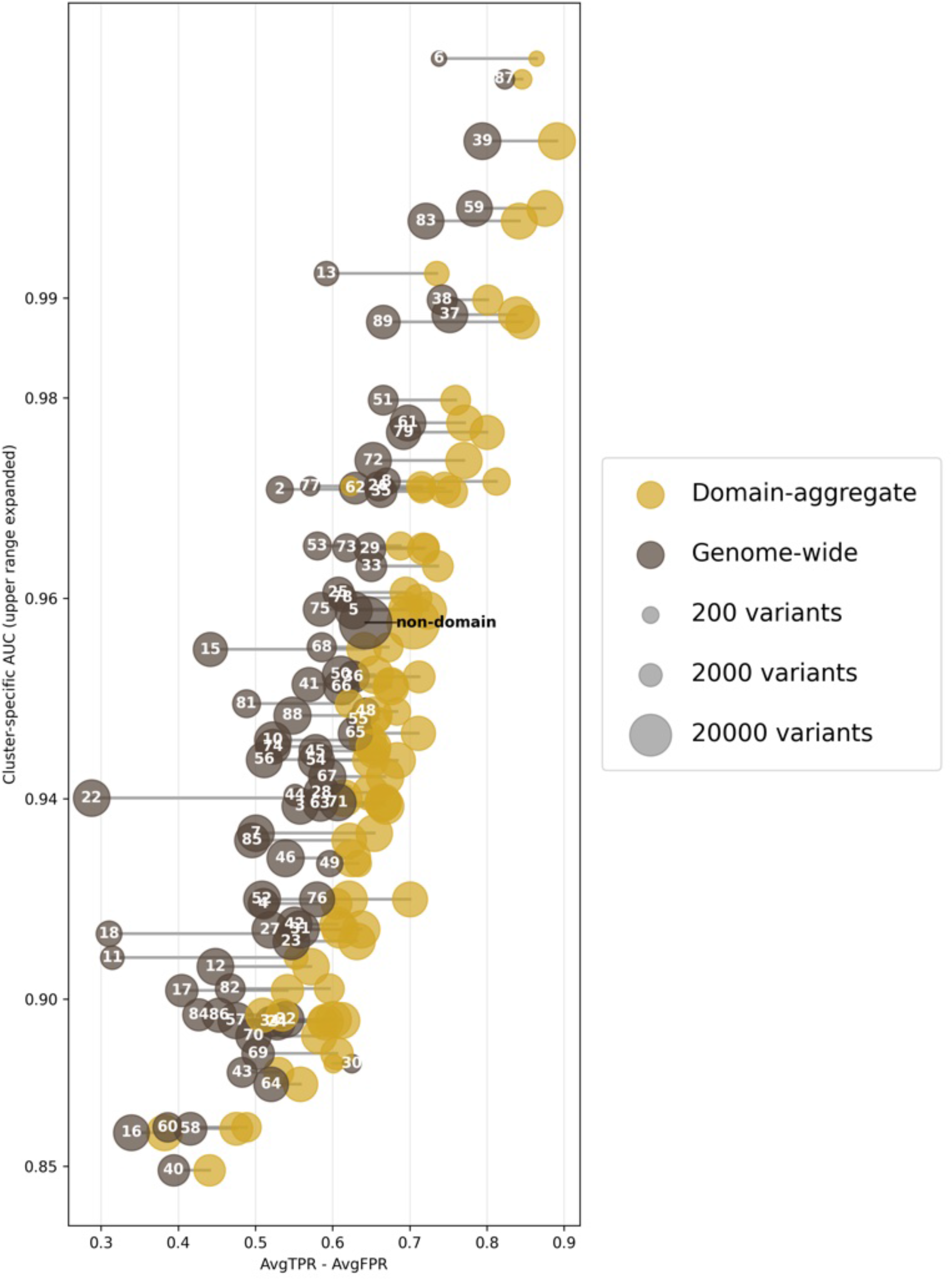
Cluster-specific calibration performance and AlphaMissense AUC. Cluster-specific performance (AvgTPR − AvgFPR) versus cluster-specific AlphaMissense AUC shown as a dumbbell plot. Each cluster is represented by a pair of connected points (domain-aggregate calibration vs. genome-wide calibration). The x-axis shows the domain-aggregate performance metric (AvgTPR − AvgFPR), and the y-axis shows the cluster-specific REVEL AUC. Higher values of AvgTPR − AvgFPR indicate better discrimination performance. The y-axis is displayed on a non-linear scale, with the upper range expanded to improve visualization of differences across clusters.

**Extended Data Fig. 30:**
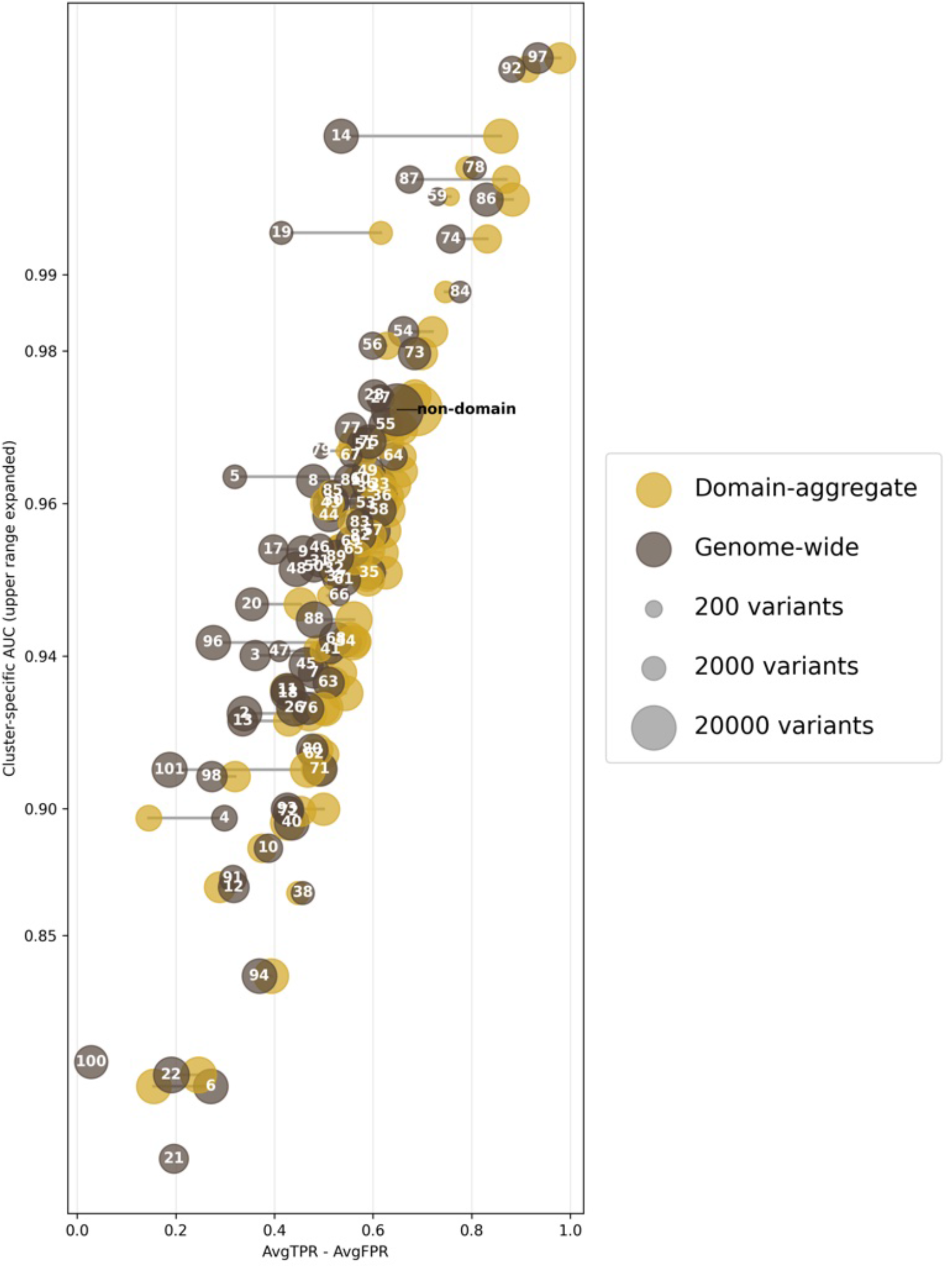
Cluster-specific calibration performance and MutPred2 AUC. Cluster-specific performance (AvgTPR − AvgFPR) versus cluster-specific MutPred2 AUC shown as a dumbbell plot. Each cluster is represented by a pair of connected points (domain-aggregate calibration vs. genome-wide calibration). The x-axis shows the domain-aggregate performance metric (AvgTPR − AvgFPR), and the y-axis shows the cluster-specific REVEL AUC. Higher values of AvgTPR − AvgFPR indicate better discrimination performance. The y-axis is displayed on a non-linear scale, with the upper range expanded to improve visualization of differences across clusters.

**Extended Data Fig. 31:**
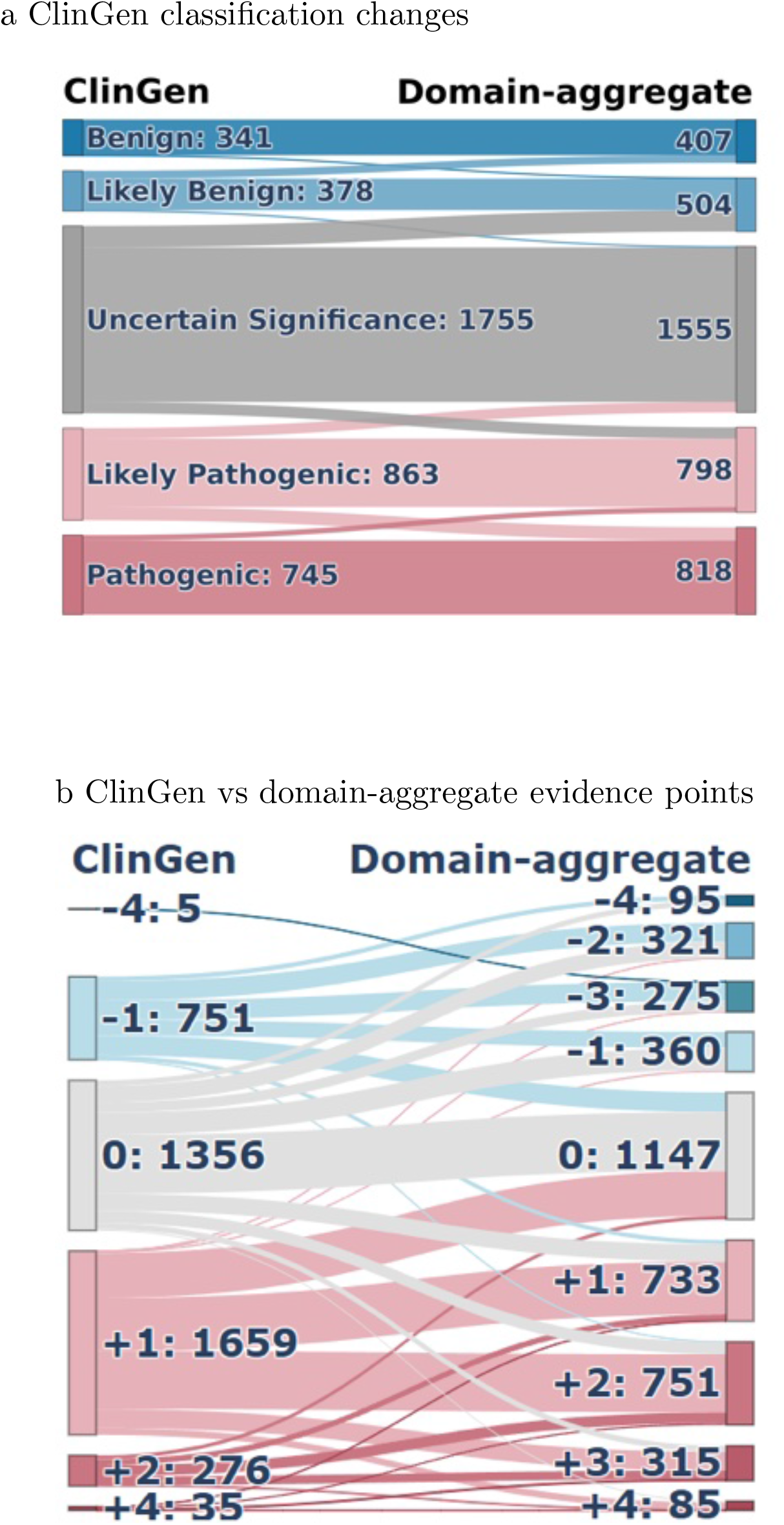
ClinGen Sankey by domain-aggregate calibration (REVEL) Sankey diagrams illustrating the impact of domain-aggregate calibration using REVEL scores. (a) Transitions in clinical classifications from original ClinGen assignments to recalibrated classifications. (b) Reassignment of computational evidence points under ClinGen guidelines compared with domain-aggregate calibration.

**Extended Data Fig. 32:**
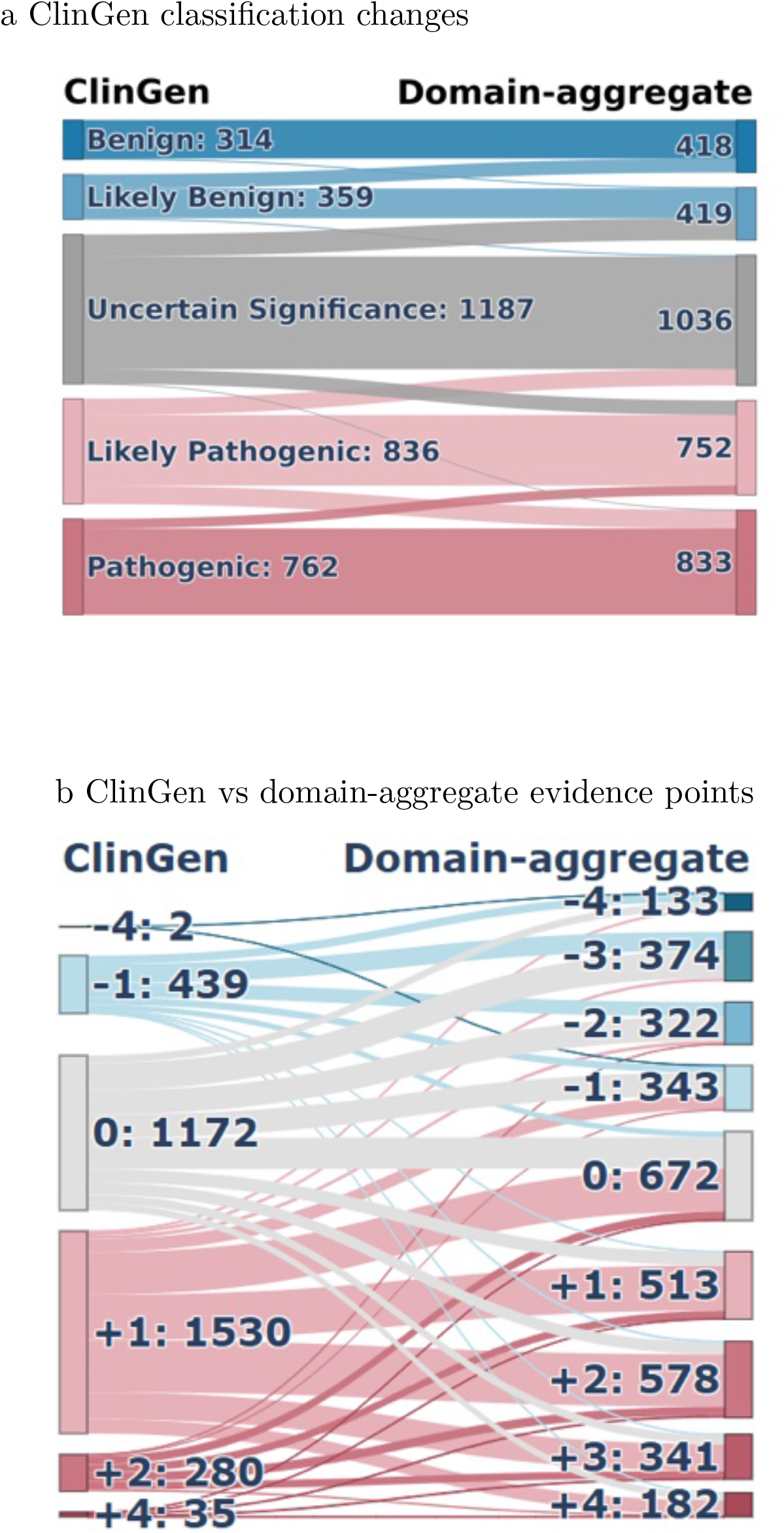
ClinGen Sankey by domain-aggregate calibration (AlphaMissense) Sankey diagrams illustrating the impact of domain-aggregate calibration using AlphaMissense scores. (a) Transitions in clinical classifications from original ClinGen assignments to recalibrated classifications. (b) Reassignment of computational evidence points under ClinGen guidelines compared with domain-aggregate calibration.

**Extended Data Fig. 33:**
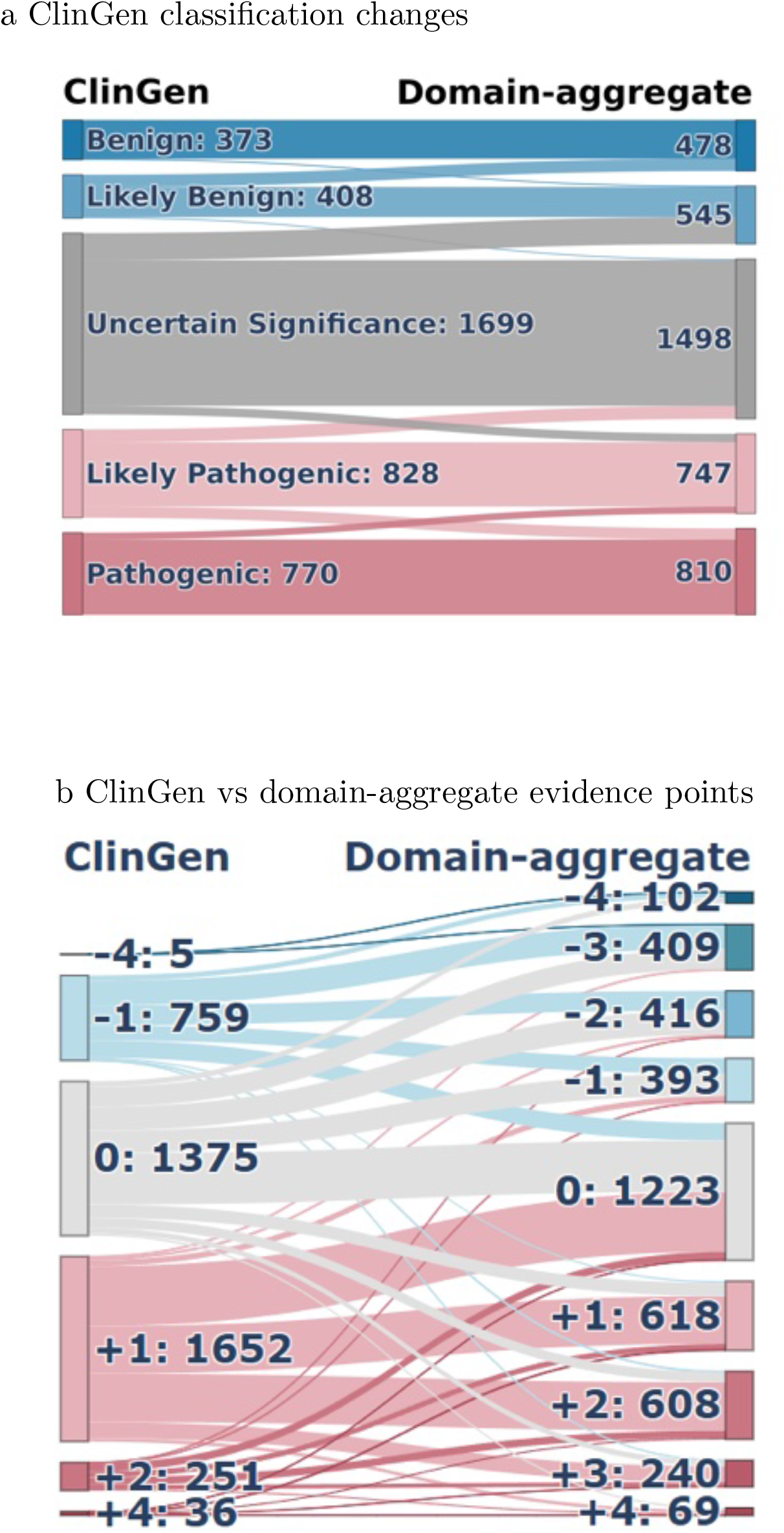
ClinGen Sankey by domain-aggregate calibration (MutPred2) Sankey diagrams illustrating the impact of domain-aggregate calibration using MutPred2 scores. (a) Transitions in clinical classifications from original ClinGen assignments to recalibrated classifications. (b) Reassignment of computational evidence points under ClinGen guidelines compared with domain-aggregate calibration.

**Extended Data Fig. 34:**
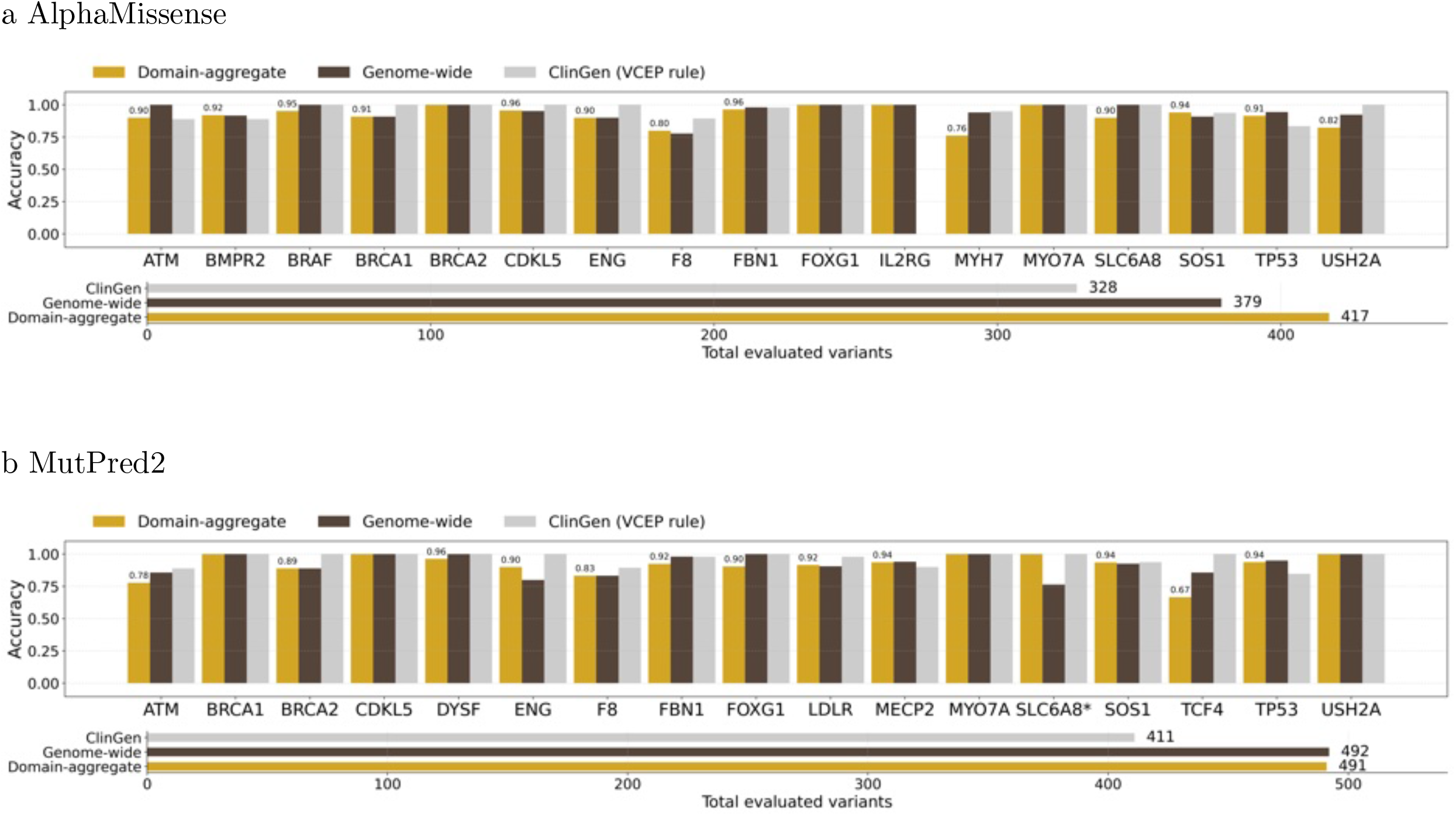
ClinGen non-circular set evaluation for domain-aggregate calibration (excluding zero-point assignments) Evaluation of per-gene classification accuracy on the ClinGen non-circular variant set using (a) AlphaMissense and (b) MutPred2 scores under three approaches: ClinGen-provided computational evidence strengths, genome-wide calibration thresholds, and domain-aggregate calibration thresholds. Top bar plots show the fraction of variants correctly classified relative to ClinGen reference classifications after recomputing variant classifications using only non-computational evidence to maintain non-circularity. Variants assigned zero computational evidence points are excluded from accuracy calculations. Bottom bar plots show the total number of variants assigned non-zero computational evidence points by each method.

**Extended Data Fig. 35:**
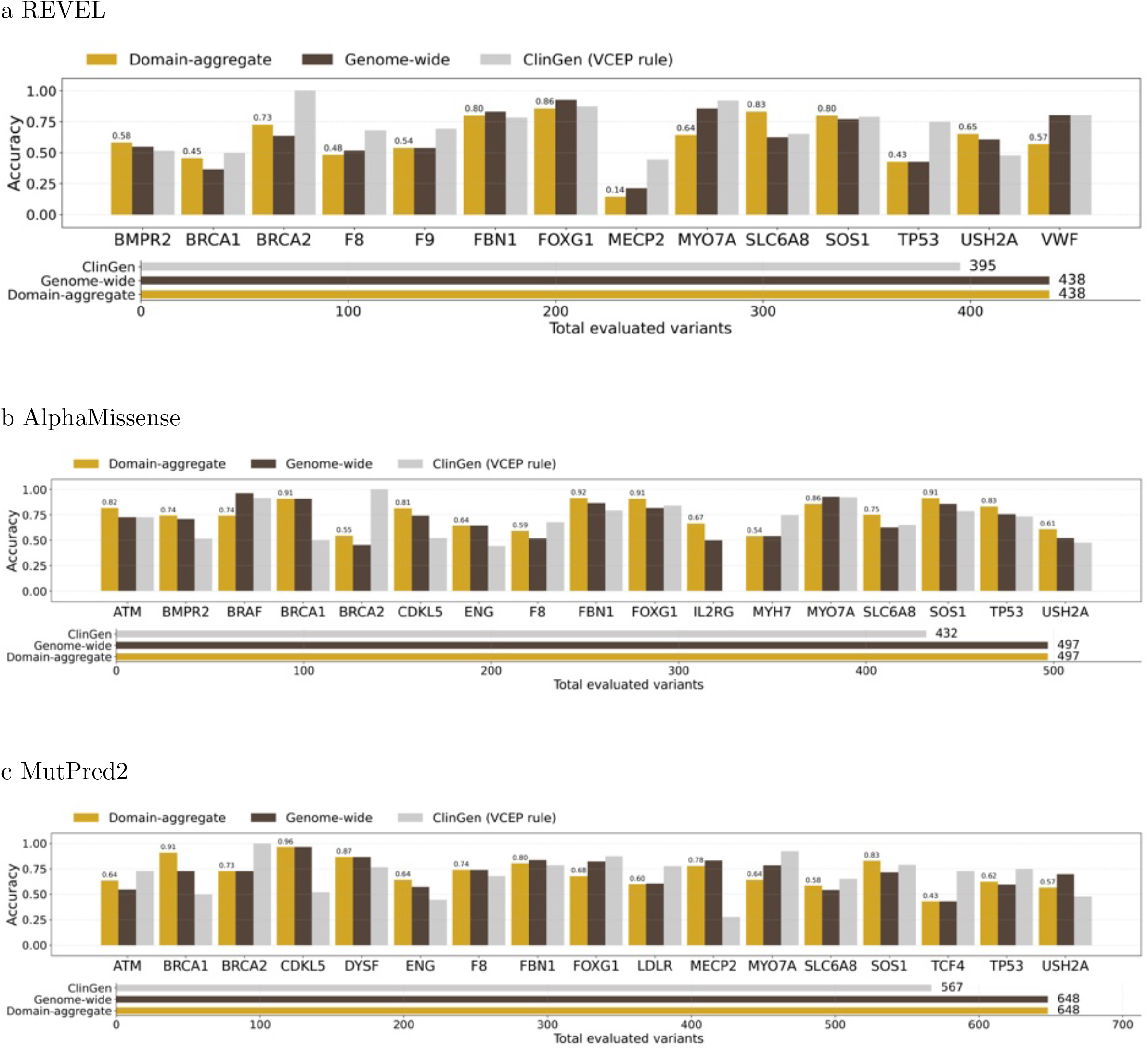
ClinGen non-circular set evaluation for domain-aggregate calibration (including zero-point assignments) Evaluation of per-gene classification accuracy on the ClinGen non-circular variant set using (a) REVEL, (b) AlphaMissense, and (c) MutPred2 scores under three approaches: ClinGen-provided computational evidence strengths, genome-wide calibration thresholds, and domain-aggregate calibration thresholds. Top bar plots show the fraction of variants correctly classified relative to ClinGen reference classifications after recomputing variant classifications using only non-computational evidence to maintain non-circularity. Variants assigned zero computational evidence points are included in the accuracy calculations. Bottom bar plots show the total number of variants assigned computational evidence points by each method.

**Extended Data Fig. 36:**
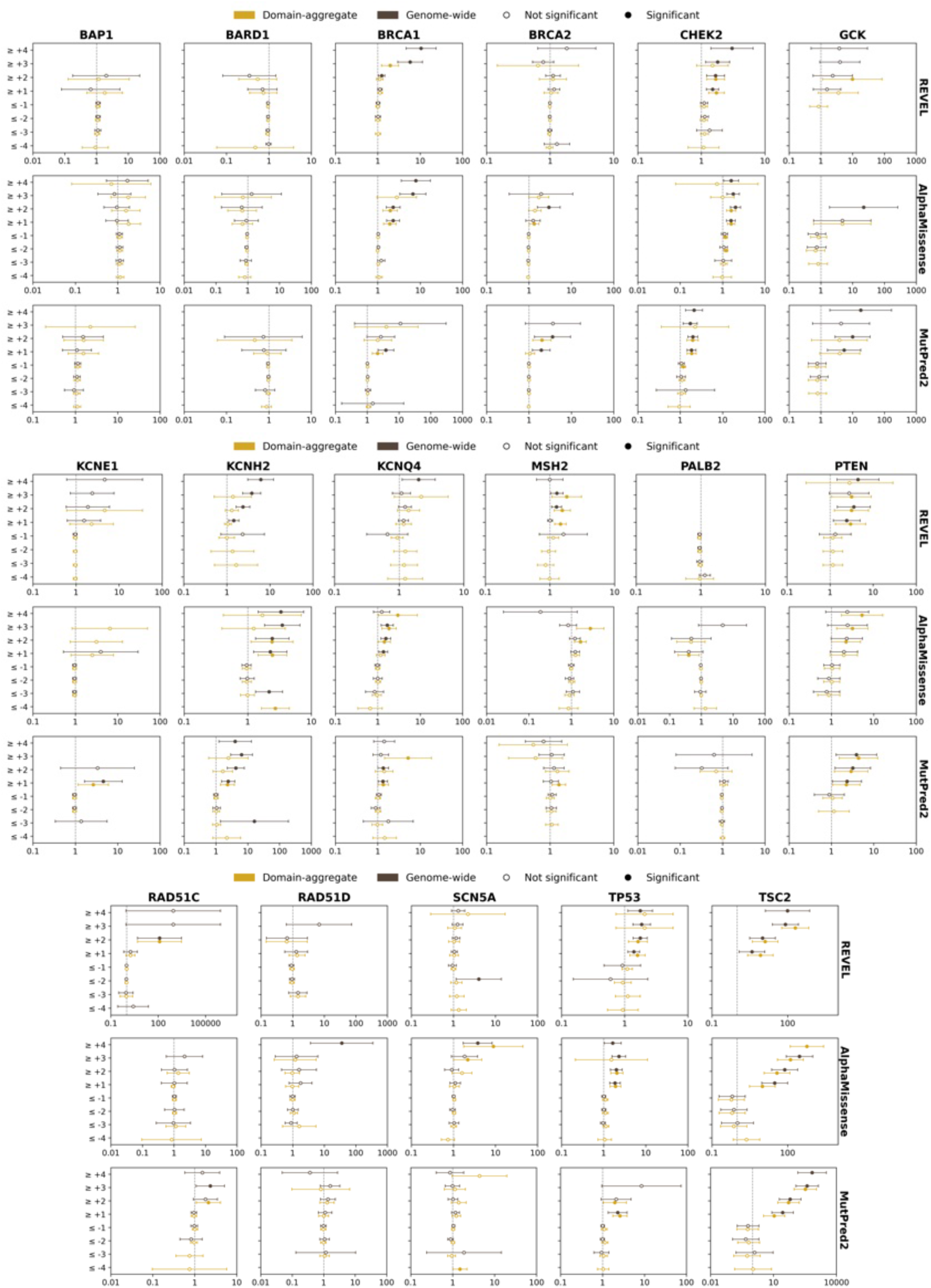
Odds ratios for disease occurrence in the All of Us biobank. Odds ratios for disease occurrence in the All of Us biobank for variants meeting different evidence strength thresholds in example genes using domain-aggregate calibration compared with genome-wide calibration. The x-axis shows the odds ratio (vertical dashed line indicates OR = 1), and the y-axis shows total evidence points for variant sets. Circles represent estimated odds ratios with 95% confidence intervals (whiskers); filled circles indicate statistically significant associations.

**Extended Data Fig. 37:**
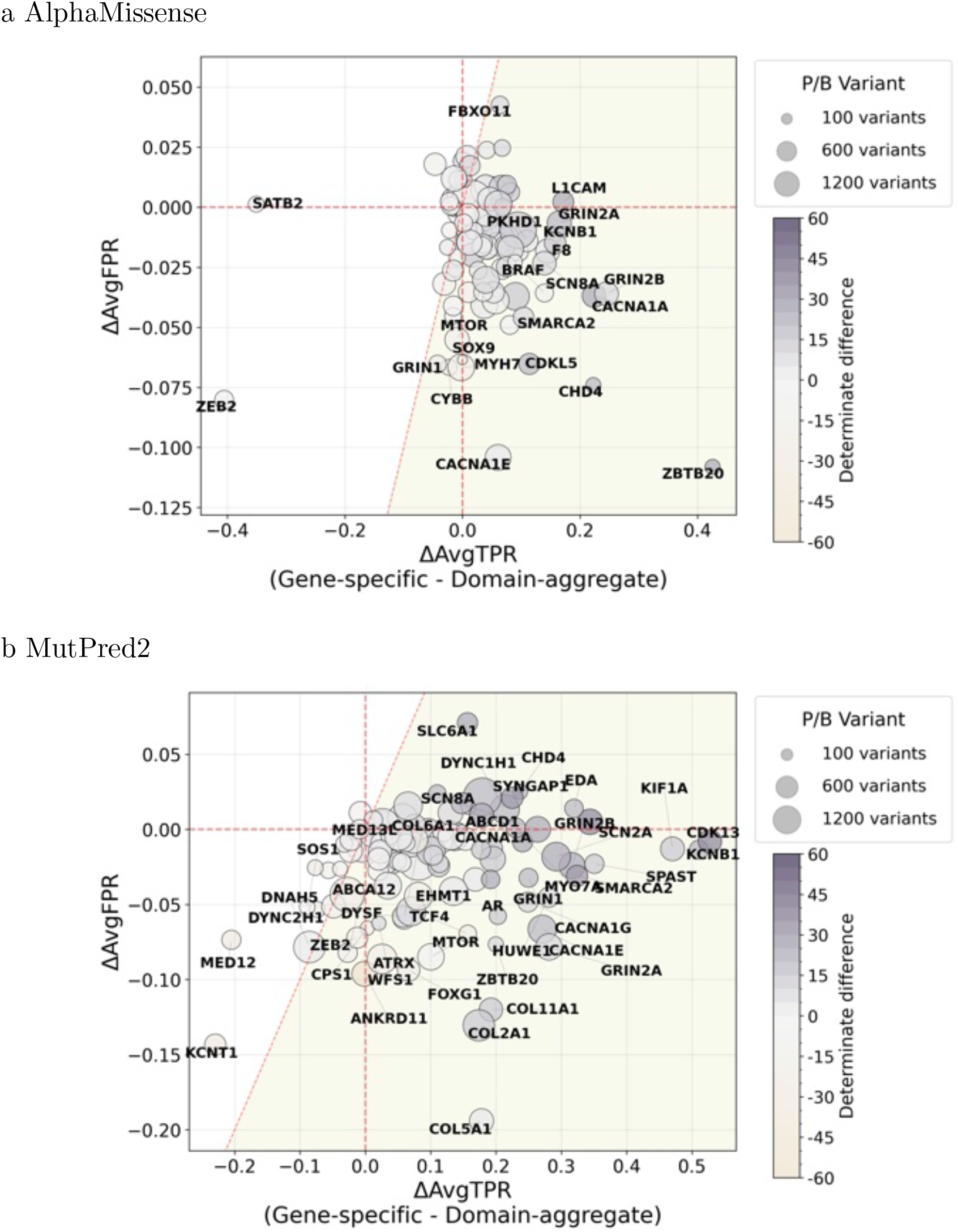
Calibration method comparison: AlphaMissense and MutPred2 (per-gene) Gene-level changes in sensitivity and false positive rate comparing gene-specific calibration versus domain-aggregate calibration. (a) Results using AlphaMissense scores. (b) Results using MutPred2 scores. Each point represents a gene. The x-axis shows the average change in true positive rate (ΔAvgTPR) between gene-specific calibration and domain-aggregate calibration (gene-specific minus domain-aggregate), and the y-axis shows the corresponding change in false positive rate (ΔAvgFPR). Point color encodes the change in evidence coverage (fraction of variants receiving at least -/+1 point of evidence), with deeper grey indicating higher coverage under the gene-specific method. Point size is proportional to the number of pathogenic and benign variants (*n*_PB_; P/LP and B/LB). The shaded area indicates increased performance (below the dashed red diagonal *x* = *y*).

**Extended Data Fig. 38:**
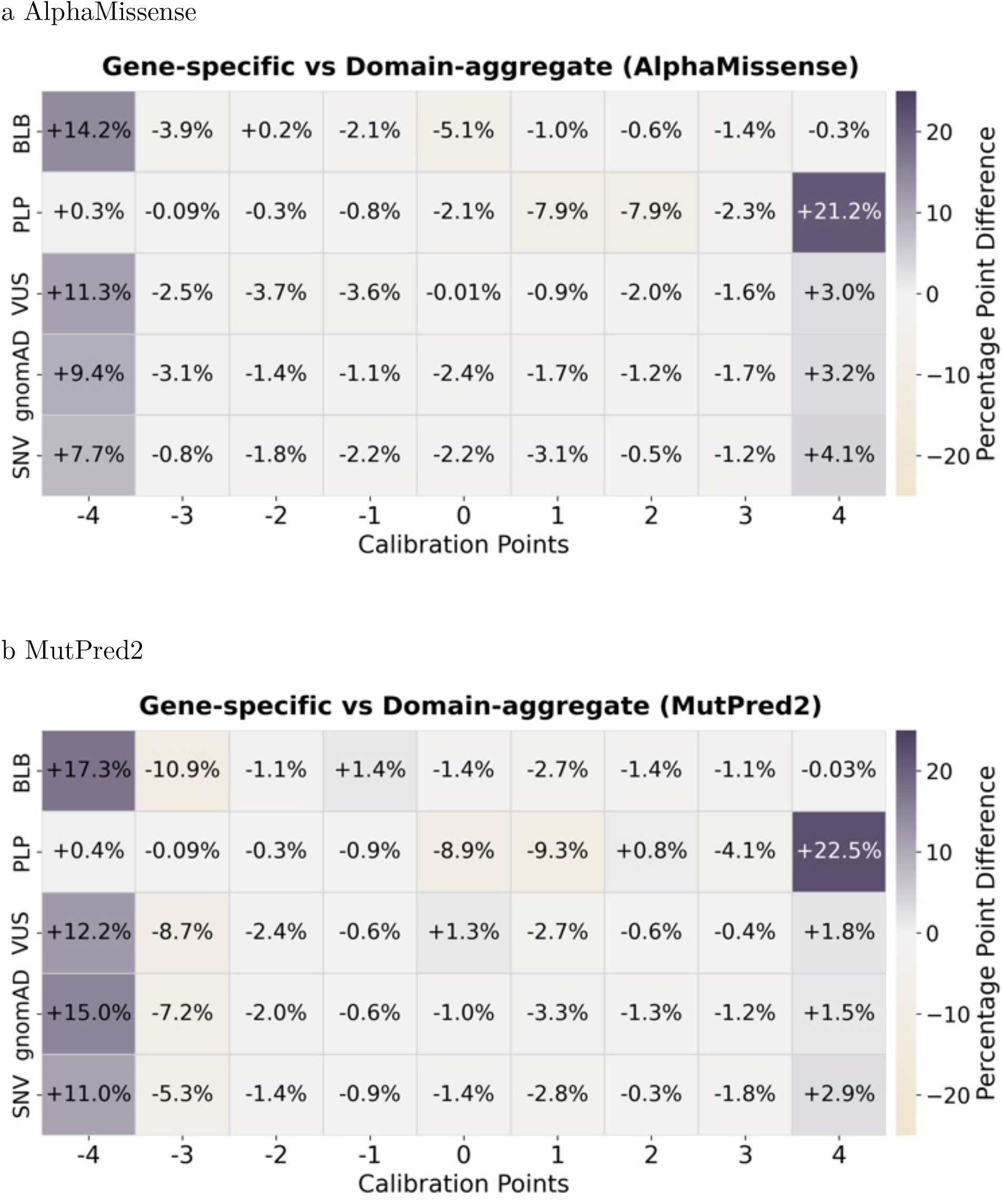
Evidence point assignment difference (gene-specific vs. domain-aggregate): AlphaMissense and MutPred2. Comparison of evidence point assignments between gene-specific and domain-aggregate calibration methods. (a) Results using AlphaMissense scores. (b) Results using MutPred2 scores. Heatmaps display percentage point differences in evidence point assignments stratified by variant classification. Darker grey shades indicate higher assignment rates under the gene-specific calibration method compared to domain-aggregate calibration.

**Extended Data Fig. 39:**
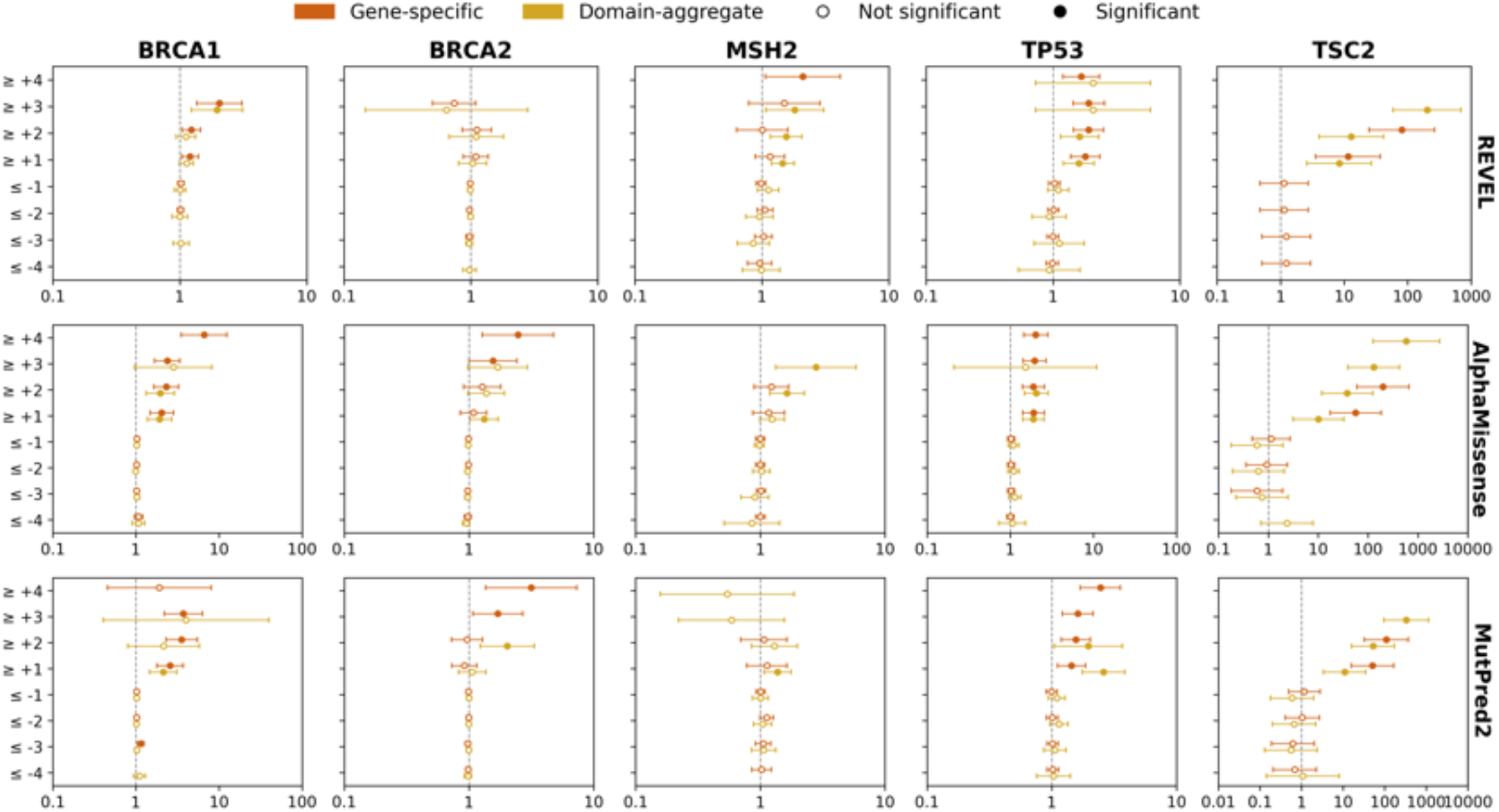
Odds ratios for disease occurrence in the All of Us biobank. Odds ratios for disease occurrence in the All of Us biobank for variants meeting different evidence strength thresholds in example genes using gene-specific calibration compared with domain-aggregate calibration. The x-axis shows the odds ratio (vertical dashed line indicates OR = 1), and the y-axis shows total evidence points for variant sets. Circles represent estimated odds ratios with 95% confidence intervals (whiskers); filled circles indicate statistically significant associations.

**Extended Data Fig. 40:**
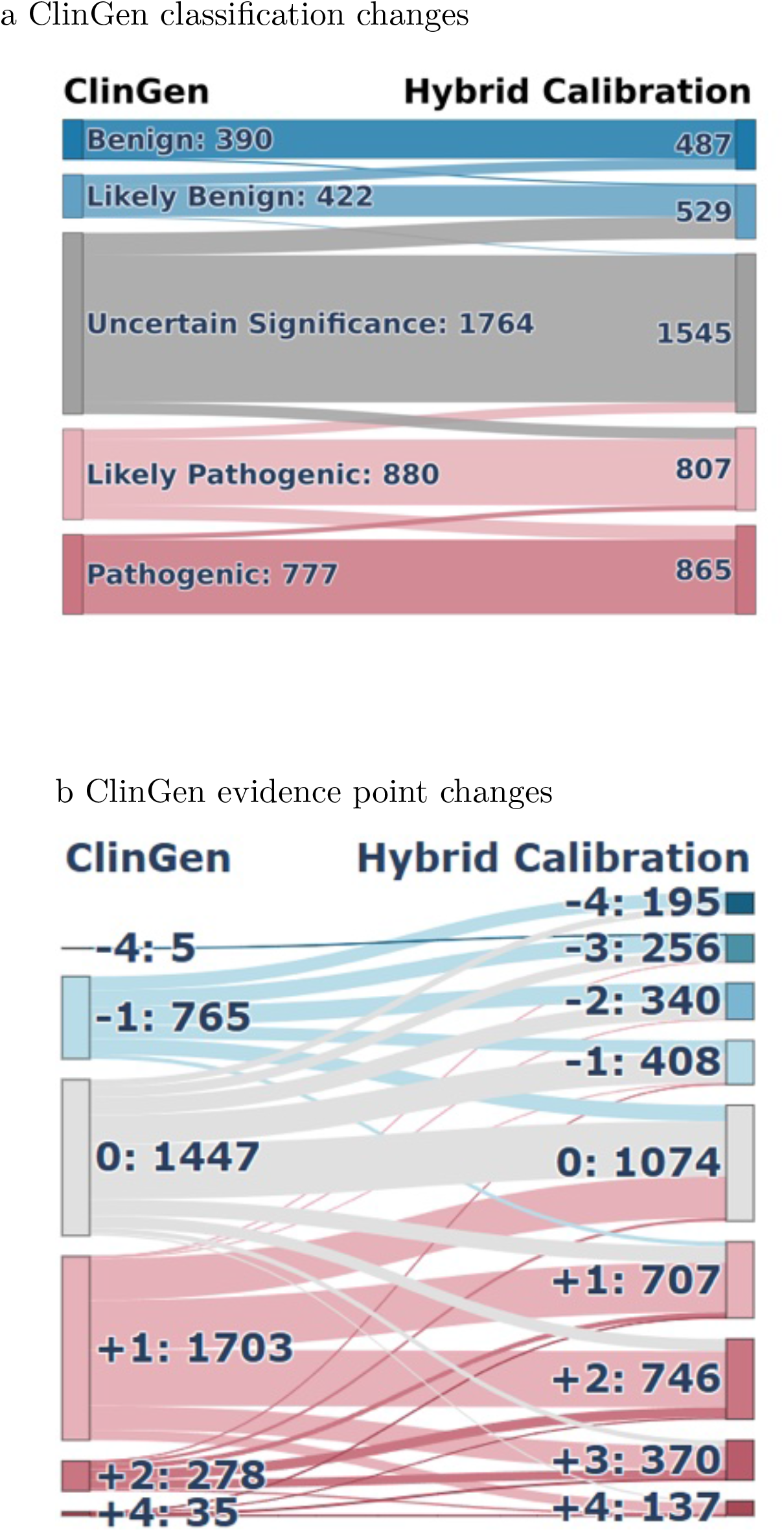
ClinGen Sankey by hybrid calibration (REVEL) Sankey diagrams illustrating the impact of hybrid calibration using REVEL scores. (a) Transitions in clinical classifications from original ClinGen assignments to recalibrated classifications. (b) Reassignment of computational evidence points under ClinGen guidelines compared with hybrid calibration.

**Extended Data Fig. 41:**
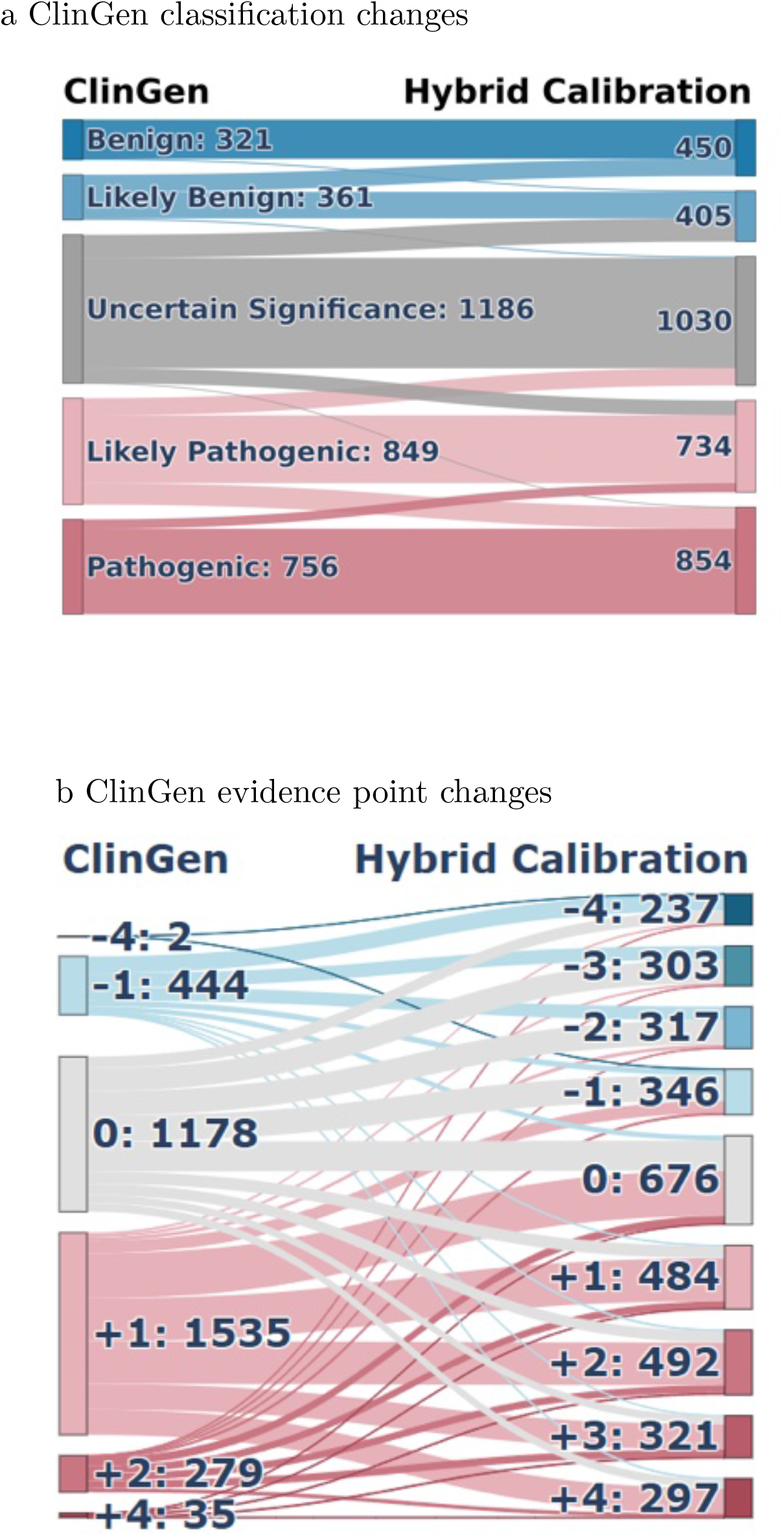
ClinGen Sankey by hybrid calibration (AlphaMissense) Sankey diagrams illustrating the impact of hybrid calibration using AlphaMissense scores. (a) Transitions in clinical classifications from original ClinGen assignments to recalibrated classifications. (b) Reassignment of computational evidence points under ClinGen guidelines compared with hybrid calibration.

**Extended Data Fig. 42:**
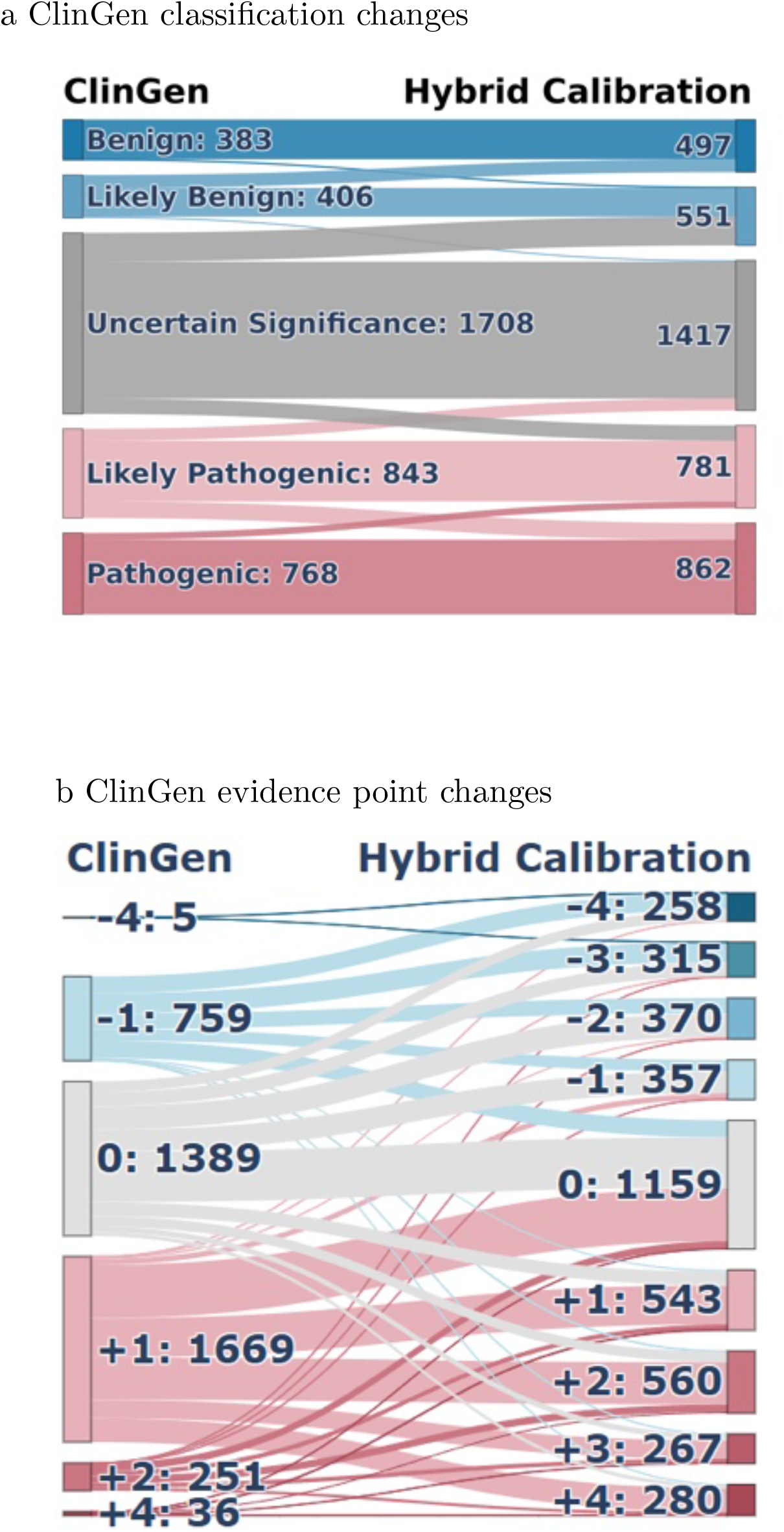
ClinGen Sankey by hybrid calibration (MutPred2) Sankey diagrams illustrating the impact of hybrid calibration using MutPred2 scores. (a) Transitions in clinical classifications from original ClinGen assignments to recalibrated classifications. (b) Reassignment of computational evidence points under ClinGen guidelines compared with hybrid calibration.

