## Supplementary Table Notes for "Gene- and domain-aware calibration increases the clinical utility of variant effect predictors"

**Supplementary Tables**

**Supplementary Table 1:** Gene-specific prior probability estimates for 648 genes inferred using the adaptive DistCurve algorithm.

**Supplementary Table 2:** Gene-specific calibration evidence point intervals for each gene derived separately for REVEL, AlphaMissense, and MutPred2. Each predictor is provided in a separate spreadsheet.

**Supplementary Table 3:** Comparison of interval likelihood ratios (LR) relative to the expected LR for each gene. For each predictor (REVEL, AlphaMissense, and MutPred2, provided in separate spreadsheets), the table reports the number of evidence intervals “won” by gene-specific calibration versus genome-wide aggregation. The Winner column indicates which method had the higher count of winning intervals; a tie indicates that both methods achieved the same number of wins for that gene.

**Supplementary Table 4:** Number of variants and genes compiled for different VEPs after filtering steps.

**Supplementary Table 5:** Cluster-specific prior probability estimates for clusters defined separately by REVEL, AlphaMissense, and MutPred2. Each predictor is provided in a separate spreadsheet.

**Supplementary Table 6:** Cluster-specific calibration evidence point intervals for each cluster derived separately for REVEL, AlphaMissense, and MutPred2. Each predictor is provided in a separate spreadsheet.

**Supplementary Table 7:** Comparison of interval likelihood ratios (LR) relative to the expected LR for each cluster. For each predictor (REVEL, AlphaMissense, and MutPred2, provided in separate spreadsheets), the table reports the number of evidence intervals “won” by domain aggregation calibration versus genome-wide aggregation. The Winner column indicates which method had the higher count of winning intervals; a tie indicates that both methods achieved the same number of wins for that cluster.

**Supplementary Table 8:** Hybrid calibration of ClinGen variants, comparing assigned evidence points. Genes were calibrated using gene-specific methods where available and domain-aggregate approaches otherwise, with the calibration strategy specified for each gene.

**Supplementary Table 9:** Comparison of interval likelihood ratios (LR) relative to the expected LR for each gene. For each predictor (REVEL, AlphaMissense, and MutPred2, provided in separate spreadsheets), the table reports the number of evidence intervals “won” by gene-specific calibration versus **acmgscaler**. The Winner column indicates which method had the higher count of winning intervals; a tie indicates that both methods achieved the same number of wins for that gene.

**Supplementary Table 10:** Summary of gene-level calibration performance and association statistics across variant classes. The table reports, for each combination of predictor (REVEL/AlphaMissense/MutPred2), gene, and calibration approach (gene-specific/domain-aggregate/genome-wide calibration), the defined variant classification ranges based on calibration point thresholds. For each class, the natural logarithm of the odds ratio (LogOR) and its 95% confidence interval (LogOR_LI, LogOR_UI) are provided, along with the corresponding two-sided Wald test p-value assessing deviation from the null hypothesis (LogOR = 0). Case inclusion and control exclusion phenotypes specify the criteria used to define case and control cohorts, respectively. Gene identifiers are provided using Ensembl gene IDs (ENSG).
